# PCRedux: A Data Mining and Machine Learning Toolkit for qPCR Experiments

**DOI:** 10.1101/2021.03.31.437921

**Authors:** Michał Burdukiewicz, Andrej-Nikolai Spiess, Dominik Rafacz, Konstantin Blagodatskikh, Jim Huggett, Matthew N. McCall, Peter Schierack, Stefan Rödiger

## Abstract

**Motivation:** Quantitative Real-time PCR (qPCR) is a widely used -omics method for the precise quantification of nucleic acids, in which the result is associated with the presence/absence or quantity of a specific nucleic acid sequence. As the amount of qPCR data increases worldwide, the manual assessment of results becomes challenging and difficult to reproduce. To overcome this, some automatable characteristics of amplification curves have been described in the literature, often with an appropriate “rule of thumb”.

**Results:** We developed *PCRedux* to analyze and calculate 90 numerical qPCR amplification curve descriptors (‘‘features”) from large datasets of qPCR amplification curves that are aimed for interpretable machine learning and development of decision support systems. In a case study of a diverse dataset with 3181 positive, negative and ambiguous amplification curves, as assessed by three human raters, we demonstrate a sensitivity >99 % and specificity >97 % in detecting positive and negative amplification. *PCRedux* is unique as it goes beyond traditional qPCR analysis to capture curvature properties that improve the characterization and classification of amplification curves. The calculation of the features is reproducible and objective, since *R* is used as a controllable working environment. *PCRedux* is not a black box, but open source software following on the principle of mathematically interpretable features. These can be combined with user-defined labels for automatic multi-category classification and regression in machine learning.

**Availability:** https://cran.r-project.org/package=PCRedux. Web server: http://shtest.evrogen.net/PCRedux-app/. Documentation: https://PCRuniversum.github.io/PCRedux/.

## Introduction

Quantitative Real-time PCR (qPCR) is widely employed for gene expression analysis (GEA) in pharmacology, medicine and forensics [2,8]. qPCR has become a staple tool for high-throughput technology validation such as validation of sequencing (SI 3, 5). Notwithstanding, it is not trivial to perform a reproducible qPCR data analysis since the majority of the experiment is assessed manually [20]. As the -omics data volume from high-throughput technologies is increasing, there is a demand for tools for data processing within automated pipelines [5].

Reproducible discrimination between positive, negative and unusual amplification curves (ACs), indicating presence, absence, or contamination of specific nucleic sequence, poses a challenge for many users and is not standardized (SI 3, 5). Others and we have developed algorithms to process qPCR data in an automated fashion (SI 3, 2 & 5), including amplification curve preprocessing, cycle of quantitation (Cq) calculation and relative GEA [5,7,8,13]. To this extent, our implementations calculate Cq values based on the maxima of the first or second derivative, and not on the variable cycle threshold (Ct), where the latter is not objective and reproducible [18,19]. Generally, it is not evident whether positive or negative calls pertain to sigmoid (high quality), jagged (poor quality), or flat (negative) shaped qPCR trajectories (SI 3, 2, SI 3, Fig. 1, 4 & 5). In this respect, most software packages do not consider that the entire shape of the curve adds information beyond Cq and amplification efficiency, as it is impacted by the chemistry, fluorochrome system [12] and cycler hardware [19]. For example, noise in the background and plateau region can be considered as an indicator for amplification curve data quality. For high-throughput experiments, a manual analysis of all amplification curves is not feasible, as it is laborious, error-prone and not reproducible (SI 3, Fig. 24). Although laboratory-internal guidelines seem to facilitate the accurate evaluation of amplification curves, they are not portable to other laboratories. To date, and to the best of our knowledge, no open-source software automatically and reproducibly classifies high-throughput amplification curves by machine learning (ML). *PCRedux* aims to remedy this situation by providing a basis for the automated evaluation of qPCR results using established and new descriptive curve features that could be used to classify amplification curves.

**Figure 1.**
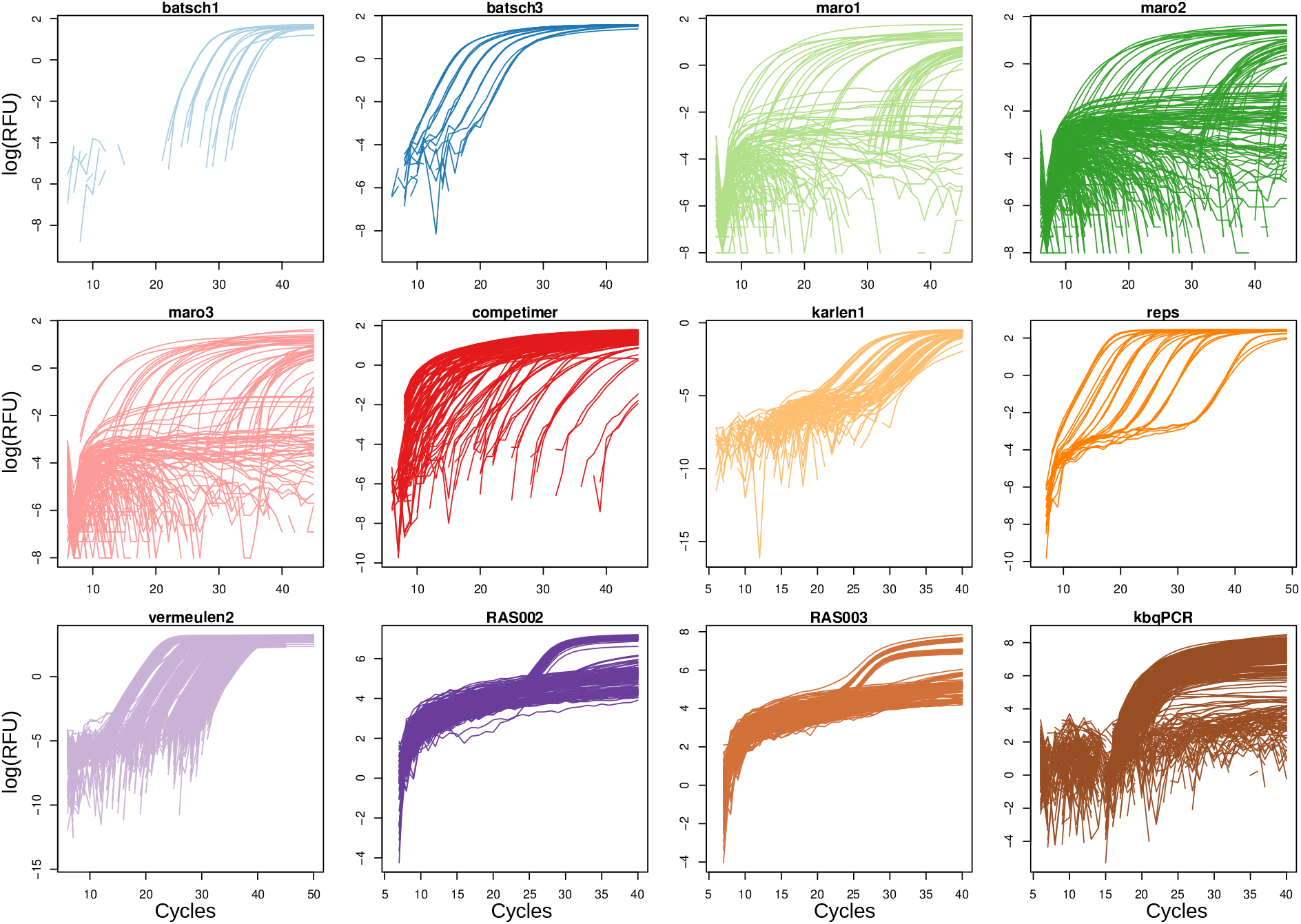
Plot of Cycles vs. logarithmized raw fluorescence values. Cycles vs. logarithmized raw fluorescence values (log(RFU)) for each of the 12 qPCR subsets used in this case study, which cover a broad Cq range (range: 6 - 50, median ± IQR: 29 ± 18, n = 3181). All raw RFU values have been baselined (median of the first 5 cycles) before log-transformation. Note that this dataset contains a total of 1658 positive, 1392 negative and 131 ambiguous amplifications, as evaluated from three different human raters. All amplification curves were selected based on a high diversity in curve shapes, where some curves are almost ideally sigmoid (e.g., batsch1, reps), while others – due to low amplification efficiency – are flat and elongated (e.g., karlen1). RFU = Raw fluorescence units.

With the availability of machine learning techniques, several groups have started to use these in qPCR experiments, such as Support Vector Machine based classifiers employed in genotyping sequence variants based on high resolution melting curves [3], or to generally improve Cq determination [6,9].

One may ask “Why not use deep learning on amplification curves?” as deep learning methods have proven to be very effective for different medical diagnostic tasks [4] and have even beaten human experts in some of these areas. Unfortunately, the black box character of the algorithms limits their medical diagnostic application, and only recent work aims at identifying those features that influence the decision for a model most. In fact, the difference in performance between complex classifiers (e.g., deep neural networks) and simple classifiers (e.g., logistic regression) is often not significant [11].

Conversely, our system is supposed to be interpretable from the beginning, so that the algorithms are based on mathematical procedures (e.g., derivatives, quotients), which can be recalculated. In addition, our approach allows domain-specific knowledge to be incorporated, such as the “hook effect” [1], and enables the user to engage in feature engineering. This way, the *PCRedux* software is accessible to the latest machine learning knowledge.

## Implementation and Results

### Software engineering

*PCRedux* ≥ v. 1.1 is a package for the R language. Unit testing and continuous integration are used for software quality control (SI 3.1). We integrated algorithms and tools published by us, including *qpcR* [10], *MBmca* [16], *chipPCR* [14], *RDML* [15], as well as other packages with references cited therein from the Comprehensive *R* Archive Network, and enhanced and extended these with further algorithms for feature extraction (SI 3, 4).

### Functions

An AC is divided into regions of interest for feature calculations (SI 3, Fig. 5). Here, *pcrfit_single()* is the core function for feature calculations of single ACs. A total of 90 features obtained from methods like non-linear regression, curve derivatives estimation, autocorrelation, Bayesian change-point analysis, robust local regression analysis, hook effect detection [1] and qPCR efficiency estimation [13] are employed for feature calculation (SI 3, 4.1). *encu()* is an extension of this function with integration of meta information (e.g., detection chemistry, thermocycler) for the analysis of large-scale data sets (SI 3, 4.1.2). The function produces a *data.frame* with 90 (binary-encoded boolean, numeric, integer, factors) features from an AC trajectory. All functions for feature calculation are hardened against missing and infinite values as described in [14]. The function *performeR()* calculates binary classification criteria (e.g., sensitivity, Cohen’s *κ*), while *qPCR2fdata()* is a helper function to convert AC data to the *fdata* format for Hausdorff distance analysis (SI 3, 4.1.1). For efficient classification labeling, the function *tReem()* for curve shape-based classification and multicategory labeling was developed (SI 3, 4.3.2).

### Datasets and Data Labeling

For the current study, a datast of 3181 curves compiled from 12 different qPCR studies and covering a wide range of Cq values (SI 2, Fig. 2), detection chemistries and qPCR hardware, as well as having a large proportion of negative and ambiguous curves (approx. 48 %) was used (Figure 1, baselined and logarithmized data; SI 2, Fig. 1, baselined raw data). The source and raw data for the present case study are given in detail in Supplemental Information 1 (‘‘Studies”, “Raw qPCR data”). Further data labels in this package for 14360 qPCR human rated curves are described in SI 3, 4.2. Human rating was conducted by three individuals, and the consensus classification adhered to the following criteria: 1) selecting the majority of the three votes, 2) in case of three divergent decisions (a/n/y), curves were classified as “a” (SI 1, “Human Rating”).

**Figure 2.**
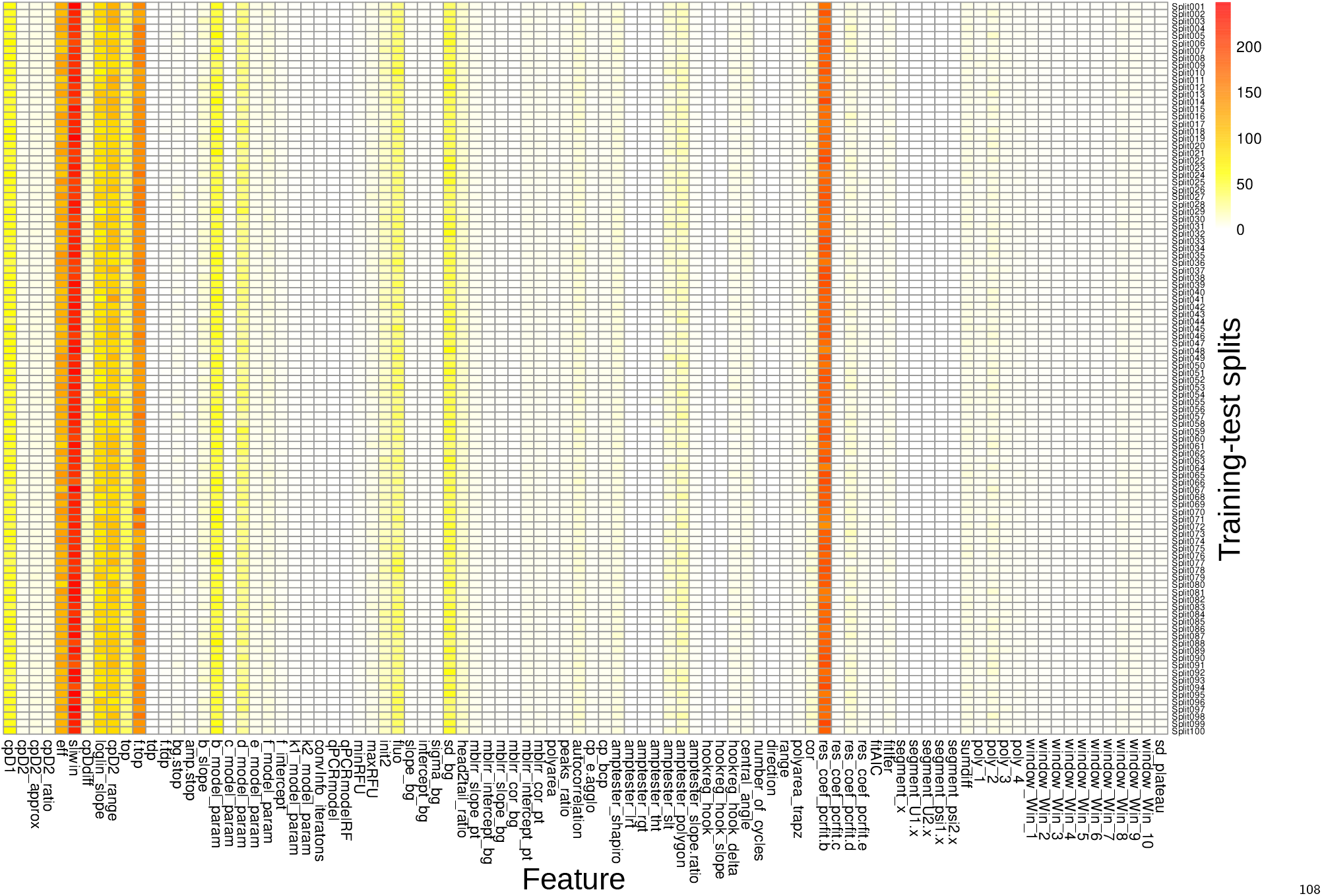
Heatmap representation of multiclass Random Forest-derived Variable Importance. Heatmap representation of multiclass Random Forest-derived Variable Importance (VI, as measured by the Mean Decrease Gini Index) of the 90 variables (curve features) extracted from the dataset represented in Fig. 1, for a total of 100 random 80 % / 20 % training-test splits (Split001 - Split100). Specifically, the encu() function was used to calculate these features from the amplification curves, including Cq values, amplification efficiencies (e.g., sliwin), noise components (e.g., sd_bg) and many other features. The highest VI’s (yellow to darkred) are displayed by features that describe the post-exponential linear region, e.g. cpD1, sliwin, res_coef_pcrfit.b, for discussion see main text. Note, that the VI is highly stable throughout the 100 splits.

### Graphical user interface

*PCRedux* also includes the graphical user interface *run_PCRedux()* with which the amplification curves can be analyzed in a graphical user interface (local or web-server) that outputs the features as a table and makes them accessible to other analysis tools. Users can upload amplification curve data and download calculated amplification curve features. Functions can be accessed through RScript or via graphical user interfaces for *R* (e.g, *RStudio* or *RKWard* [17]).

In a first step, all 90 implemented features were acquired for each of the 3181 curves (SI 1, ‘Features”), providing an extensive feature table. This feature table was then used as an input to a Random Forest-based classification approach (*R* package *randomForest*), in which 100 random subset samples (SI 2, Fig. 3) of 20 % (*test set*) were predicted from the remaining 80 % (*training set*), using the consensus classification as response target (SI 1, “RF Predictions”). At the same time, Variable Importances based on the Mean Decrease Gini Index were obtained for all features in each of the 100 splits (SI 1, ‘Variable Importance”), showing a wide range of values and indicating high importance on classification for approx. 15 variables for this specific classification task (Figure 2, yellow to darkred). Finally, for all 100 splits, a diverse set of performance measures (Accuracy, F1, Sensitivity, Specificity, multiclass-AUC) was acquired (SI 1, ‘Performance”) and summarized (Table 1; SI 1, ‘Performance Summary”; SI 2, Fig. 4).

**Table 1.**
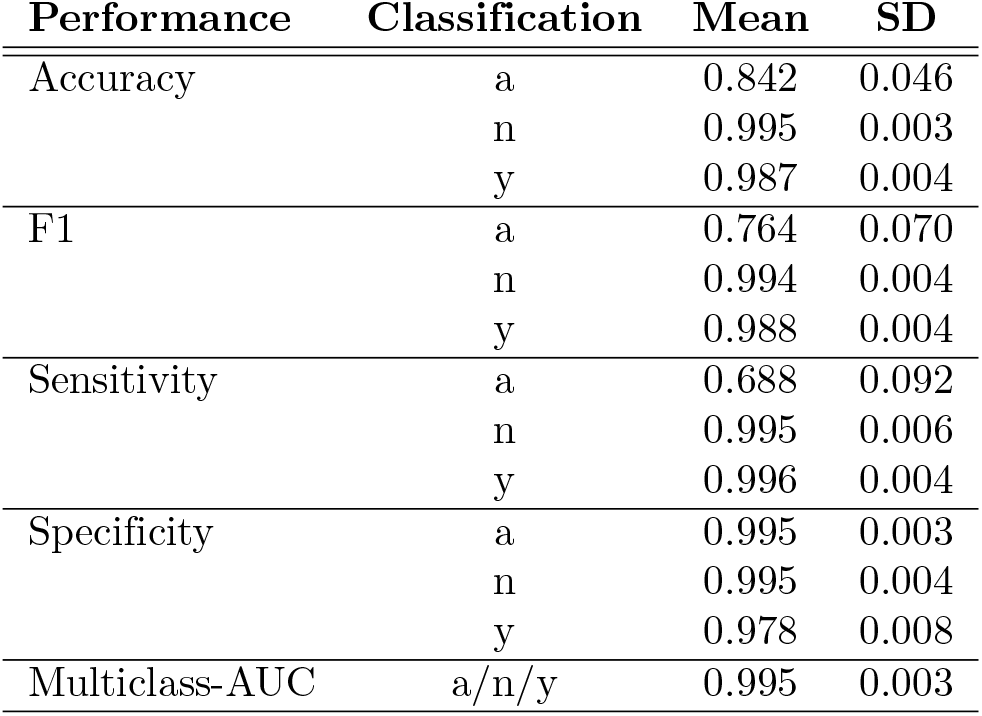
Summary of the five different performance measures Accuracy, F1, Sensitivity, Specificity and Multiclass-AUC (*R* package *pROC*) for the three classes (a)mbiguous, (n)o, (y)es, as obtained from the 80 % / 20 % class prediction of 100 random splits.

## Discussion

### Subsection heading

*PCRedux* is the first open-source software facilitating a comprehensive extraction of a diverse set of numerical and analytical explainable descriptive features from ACs for machine learning, for which we envision a broad spectrum of applications.

To illustrate how *PCRedux* can assist in classifying ACs (differentiation between positive and negative amplification curves) by single features, simple examples employing binomial logistic regression or k-means clustering are shown in the Supplemental Information (SI 3, 4.1.7 & 4.1.9).

In this work, we have used *PCRedux* to classify 3181 human-rated qPCR curves, trichotomized into the classes *ambiguous, no* and *yes*, by means of Random Forest-based machine learning and 100-fold 20 % subsampling. The summarized performance of this approach (Table 1; SI 2, Fig. 4) is quite conclusive: negative and positive amplifications were classified with >99 % sensitivity and >97 % specificity, with an F1-score (weighted average of precision and sensitivity) of approx. 99 % for both. Hence, the majority of positive and negative amplifications were classified according to the human rating. Unsurprisingly, ambiguous curves (“a”) exhibited a somewhat lower classification performance, in which only 68.8 % of these were classified according to human rating (sensitivity), i.e. 31.2 % were classified as either negatives or positives. However, with a specificity of 99.5 %, only 1 in 200 negatives/positives was classified as ‘ambiguous”. Interestingly, a closer inspection of the 68.8 %‘ambiguous” Random Forest-classified amplification curves revealed that these had been largely rated discordantly (a/n/y) between all three raters (SI 1, “RF ambiguous”).

It must be emphasized that our Random Forest-based classification approach is based on the complete space of 90 features without recursive feature elimination. Looking at the Variable Importances in Figure 2, it is conspicuous that features with high importance are those describing qPCR efficiencies (eff, sliwin), the exponential region (top, ftop, cpD2_range) and the slope or first derivative maximum of the post-exponential linear region (cpD1, res_coef_pcrfit.b). This finding strongly suggests that features describing the curve structure between the baseline and plateau phases can be exploited to automatically discriminate typical sigmoidal positive amplification curves from negative or ambiguous ones.

In our scenario, we have only distinguished positive, negative and ambiguous curves, however it should possible to exploit the feature list as to incorporate features that describe noise or other QC measures. Other possible applications include:

- models to assess the quality of amplification curves (e.g., batch control of chemicals),
- integration in decision support systems of diagnostic applications,
- integration in data validation tools (e.g., as a pre-screen for academic publishing services) and
- detection of false-negative and false-positive amplification curves.

Our tool should improve the quality and reproducibility of qPCR data analysis, as it systematically validates the input data. Moreover, it supports the development of control mechanisms necessary for other automated algorithms (e.g., Cq value determination) (SI 3, 2.3) and provides open data (SI 3, 5) to expedite the automatization of qPCR data analysis. Thus, *PCRedux* may reduce the workload involved in large-scale data analysis and deliver reproducible and more objective data assessment.

## Supporting information

Microsoft EXCEL file containing raw data (incl. amplification curves, ratings)

File containing raw amplification plots and performance plots

Comprehensive introduction to the PCRedux package with explanations of the features and applications of machine learning models

## Supporting Information

- SI 1: Microsoft EXCEL file containing raw data (incl. amplification curves, ratings).
- SI 2: File containing raw amplification plots and performance plots.
- SI 3: Comprehensive introduction to the *PCRedux* package with explanations of the features and applications of machine learning models.

## Acknowledgments

Grateful thanks belong to Paulina Przybyłek and Przemysław Chojecki (Warsaw University of Technology) for the improvement of the package and the *R* community.

