## Supplementary material for "PCRedux: A Data Mining and Machine Learning Toolkit for qPCR Experiments": File containing raw amplification plots and performance plots

to

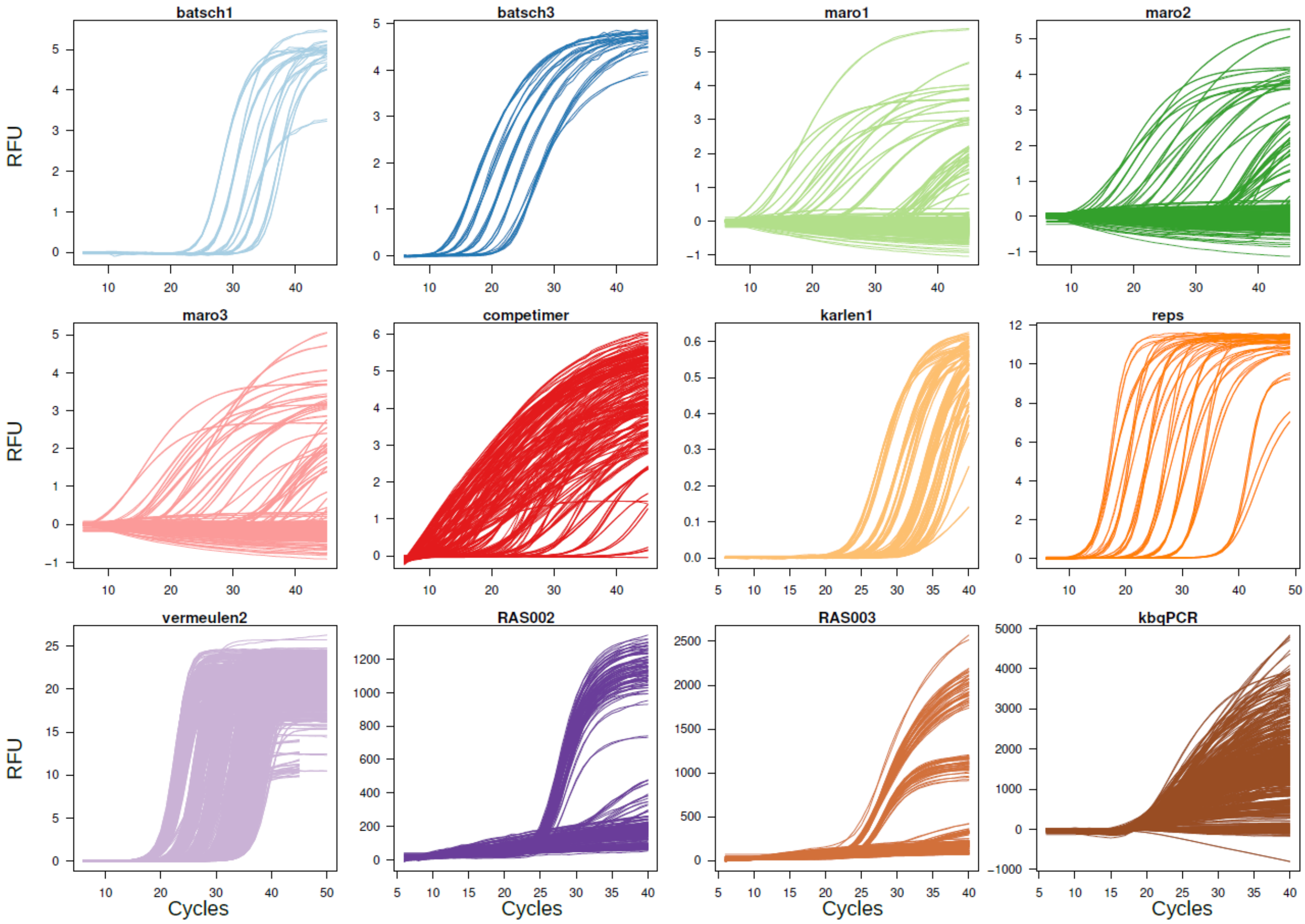


**S2 Figure 1**: Plot of Cycles vs. raw fluorescence values (RFU) for each of the 12 qPCR subsets, similar to Figure 1 of the main manuscript, but not logarithmized (n = 3181). All raw RFU values have been baselined (median of the first 5 cycles). RFU = raw fluorescence units.

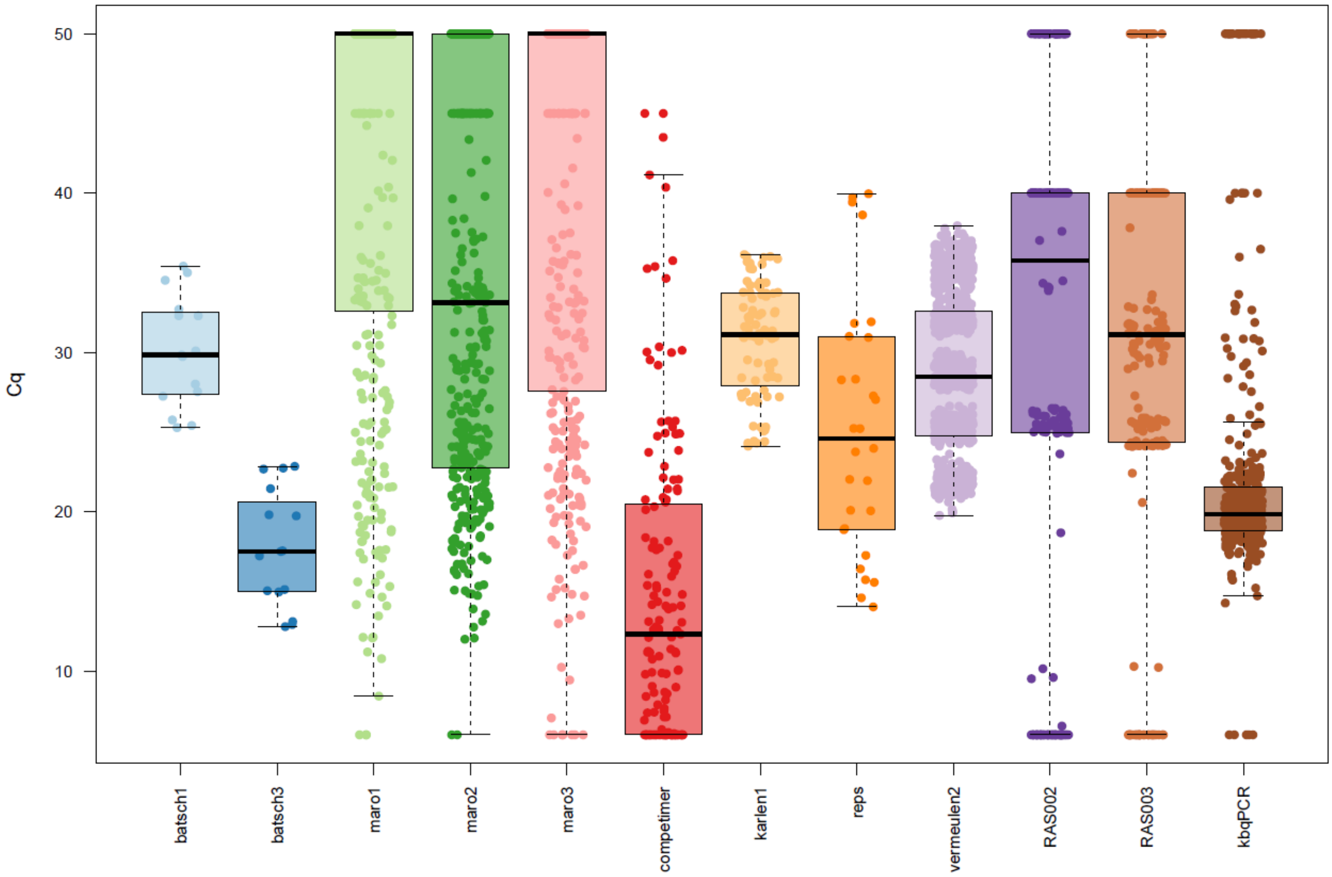


**S2 Figure 2**: Distribution of Cq values for the 12 different datasets used in this study. The datasets were selected to exhibit a wide range range of Cq values (range: 6 - 50, median ± IQR: 29 ± 18, n = 3181) with very heterogeneous distribution. The Cq values were estimated from the baselined raw data shown in S2 Figure 1 using the second derivative maximum of a 5-parameter logistic model (Ritz & Spiess, 2008). qPCR Curves for which no Cq value could be obtained (negatives with fitting error), were set to a Cq of 50.

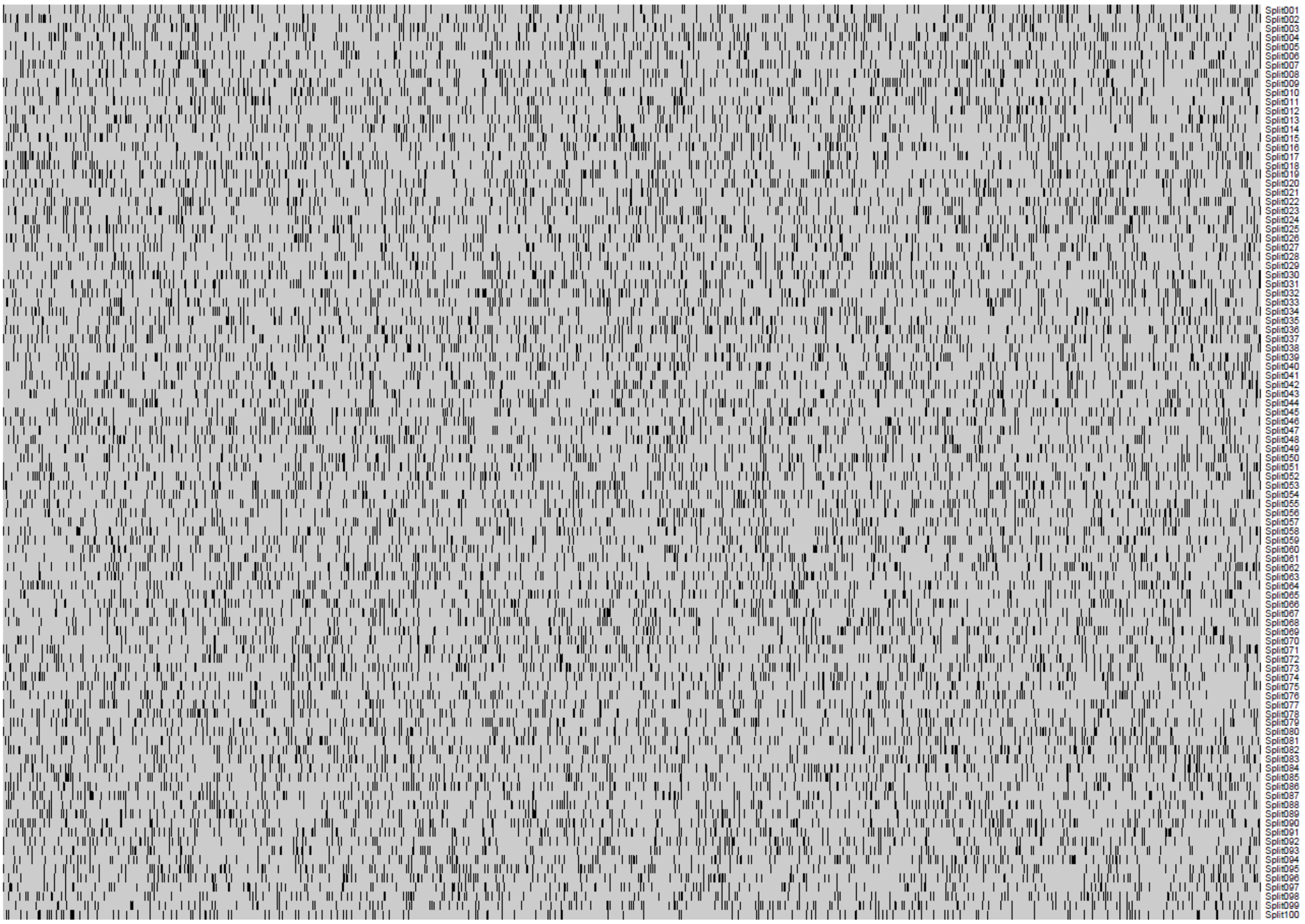


**S2 Figure 3**: 2-color matrix representation of the 100 random samplings (splits) used for obtaining the Random Forest-derived Variable Importance of the 90 features and calculating the performance measures.  Grey fields depict the 80% training set while black fields denote the 20% test set.

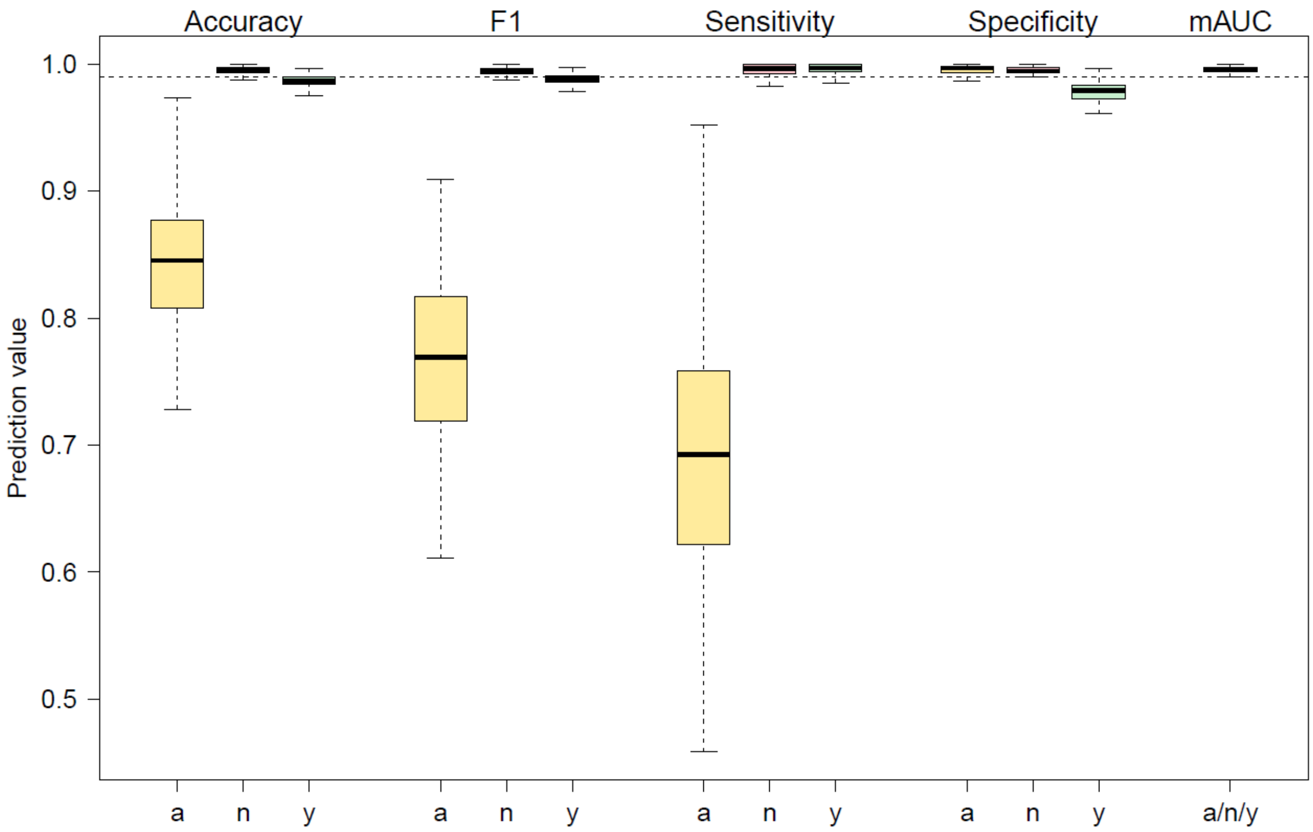


**S2 Figure 4**: Boxplot representation of the distribution of the various performance measures (Accuracy, F1, Sensitivity, Specificity and multiclass-AUC) as obtained from the 100 random sample splits of S2 Figure 3. Mean and s.d.'s of these values are given in Table 1 of the main manuscript.
