## Supplementary material for "PCRedux: A Data Mining and Machine Learning Toolkit for qPCR Experiments": Comprehensive introduction to the PCRedux package with explanations of the features and applications of machine learning models

### Supporting Information 3 (SI 3): PCRedux package - an introduction

The PCRedux package authors

2020-10-31

**5 Summary and Conclusions** **53**

**References** **56**

- A comprehensive PDF version (including domain knowledge about qPCRs and machine learning) of this document is available **online supplement**. This online document contains also the code that was used to generate the plot in this introduction.
- The code for all figures and analysis in the manuscript "PCRedux: A Data Mining and Machine Learning Toolkit for qPCR Experiments" is available at **https://github.com/PCRuniversum/PCRedux-supplements**.

### 1 Aims of the Project

A review of the literature (PubMed, Google Scholar; 1984-01-01 - 2020-10-31) and discussion with peers revealed that there is no open source software package to calculate predictors from quantitative PCR amplification curves for machine learning applications. A predictor is a quantifiable *informative* property of an amplification curve. In particular, there is no information available about predictors that can be used from amplification curves apart from measures that describe quantification points, amplification efficiencies and signal levels. Although several amplification curve data sets are available, no curated labeled data sets labels are described in the literature or repositories such as GitHub[1], Bitbucket[2], SourceForge[3] or Kaggle[4].

Therefore, the aim of the study was to:

1. create a collection of classified amplification curve data,
2. propose algorithms that can be used to calculate predictors from amplification curves,
3. evaluate pipelines that can be used for an automatic classification of amplification curves based on the curve shape and
4. to bundle the findings in a public repository open source software and open data package.

### 2 Introduction and important infromation about the `PCRedux` package

In the **online supplement** the reader finds an introduction to nucleic acids, including nucleic acid detection methods (e.g., melting curve analysis, photometric measurements) for the analysis of patient and forensic

[1] https://github.com/
[2] https://bitbucket.org/
[3] https://sourceforge.net/
[4] https://www.kaggle.com/

sample material. Special intention is paid to the quantitative Polymerase Chain Reaction (qPCR), since this method is the *de facto* standard for the detection and high-precision quantification of nucleic acids.

The focus of this study is the development of statistical and bioinformatical algorithms for the `PCRedux` software (version 1.0.7). This software can be used to automatically calculate putative predictors (*features*) from qPCR amplification curves. A predictor herein refers to a quantifiable *informative* property of an amplification curve, employable for data mining, machine learning applications and classification tasks.

On the basis of these observations, concepts for predictors (*features*) were developed and implemented in algorithms to describe amplification curves. The functions described in the following are aimed for experimental studies. It is important to note that the concepts for the predictors proposed herein emerged by a *critical reasoning* process and *domain knowledge* of the `PCRedux` package creator. The aim of the package is to propose a set of predictors, functions and data for an independent research.

#### 2.1 Development, Implementation, InstallationVersion Control and Continuous Integration

The **online supplement** deals with elements of the software engineering (e. g., continuous integration, Donald Knuth's *Literate Programming*, unit testing) used within the `PCRedux` software. The subsection 2.3 gives an introduction into qPCR data, their analysis and explains why there is a need for the `PCRedux` software. In addition, the data analysis using machine learning is concisely described, after which the work focuses on the analysis of the measured data.

The proposed algorithms were partially tested with machine learning methods. For this purpose, a brief introduction to the subject *machine learning* is given in the **online supplement**.

Information for statistical analyses of qPCR amplification curves are presented in section 4 ff. This covers the description of the curvature and the challenges of the calculations.

All scientific and engineering work depends on data. In particular, *open data* are becoming a cornerstone in science. As data sets of classified amplification curves were not available anywhere else, the **online supplement** summarizes the aggregation, maintenance, and distribution of classified qPCR amplification curve data sets. The manual classification of amplification curves is a time-consuming and error prone task when working with large data sets. To facilitate the manual analysis procedure, helper tools like `humanrater()` are presented in the the **online supplement**. A novel approach for *curve-shape based group classification* is shown too.

An achievement of this study is the extensive portfolio of statistical algorithms for predictor calculation. Central findings of the research are presented in subsection 4.1.

It is expected that these implementations will allow the automatic analysis of large data sets for machine learning applications. The expectations of the findings are critically discussed in section 5.

#### 2.2 PCRedux-app

`PCRedux-app` is a web server, based on the shiny technology (Chang et al. 2019) wrapped around the `encu()` function (subsubsection 4.1.2). An user can upload qPCR data and download obtained amplification curve features.

There are different ways to use the function.

- Through `RScript` (Scripting Front-End for R):
  - Enter the command `Rscript -e 'PCRedux::run_PCRedux()'` in a console and copy the pasted URL in a browser.
- Through Graphical User Interfaces:
  - The function can be started directly in `RStudio` or `RKWard` (Rödiger et al. 2012) by:

```
# run the Shiny app
PCRedux::run_PCRedux()
```

#### 2.3 Analysis of Sigmoid-Shaped Curves for Data Mining and Machine Learning Applications

The following sections describe `PCRedux` regarding the analysis, numerical description and predictor calculation from a sigmoid curve. A predictor herein refers to a quantifiable *informative* property of a sigmoid curve. The predictors (subsection 4.1), sometimes referred to as descriptors, can be used for applications such as data mining, machine learning and automatic classification (e. g., negative or positive amplification).

Machine learning is a scientific discipline that deals with the use of simple to sophisticated algorithms to learn from large volumes of data. A number of approaches to machine learning exist. Supervised learning algorithms are trained with data which contain correct answers (Zielesny 2011; Walsh, Pollastri, and Tosatto 2015; Fernandez-Delgado et al. 2014). This allows to create models that assign the data to the answers and use these for further processing and predictions (Tolson 2001). Unsupervised algorithms learn from data without answers. They use large, diverse data sets for self-improvement. Neural networks or artificial neural networks are a type of machine learning that roughly resembles the function of human neurons. They are computer programs that use several levels of nodes (neurons), work in parallel for learning, recognize patterns and make decisions in a human-like manner (Günther and Fritsch 2010). Deep Learning uses a deep neural network with many neuronal layers and an extensive volume of data (Shin et al. 2016). They solve complex, non-linear problems and are responsible for groundbreaking innovations through artificial intelligence, such as the processing of a natural language or images (Tolson 2001). Applications in the life sciences have already been described for each of these methods. Up to now there appears to be no study that uses machine learning for the classification of amplification curves in a scientific setting.

The determination of quantification points such as the Cq value is a typical task during the analysis of qPCR experiments. This is also covered by the `PCRedux` software in dedicated sections and the **online supplement**.

Characteristics of amplification curves that can be used for the statistical and analytical description are discussed in the **online supplement** in more in detail. The examples described focus on the concepts for **binary (dichotomous) classification** (Kruppa et al. 2014) as negative or positive. The mere binary classification into classes "positive" or "negative" is not necessarily the aim of the `PCRedux` package. Instead, it is aimed to provide a tool set for automatic **multicategory (polychotomus) classification** of amplification curves by any class conceivable. Such classification could be used for the quality of an amplification curve as negative, ambiguous and positive (Figure 1A & B). A definition for binary (dichotomous) classification and multicategory (polychotomus) classification is presented in Kruppa et al. (2014).

#### 2.4 Relation of Machine Learning to the Classification of Amplification Curves

A few scientific approaches have previously been shown in which machine learning was used for the analysis of amplification curves. The intention of Gunay, Goceri, and Balasubramaniyan (2016) was to improve the determination of Cq values, without dealing with classification. The authors postulated that they had developed an improved prediction of Cq values using a modified three-parameter model. One assumption of their approach was that their modified three-parameter model could be applied to any amplification curve. However, there are reasons why such an assumption is not valid.

In the **online supplement** it was described that a considerable proportion of amplification curves deviate clearly from a three-parameter model. Multiparametric models with more than four parameters are more frequently adapted to amplification curves. In addition, the multiparametric models tend to adapt to noise (Figure 2). Unsurprisingly, a Cq value is calculated for actually negative amplification curves, demonstrating that a three-parameter model alone cannot provide reliable predictions. However, a correct model is important for the extraction of Cq values, for the determination of predictors from the curves and consequently for classification.

Data mining and machine learning can be used for descriptive and predictive tasks during the analysis of complex data sets. Data mining uses specific methods from statistical inference, software engineering and domain knowledge to get a better understanding of the data, and to extract *hidden knowledge* from the preprocessed data (Kruppa et al. 2014; Herrera et al. 2016). All this implies that a human being interacts with the data at the different stages of the whole process as part of the workflow in data mining. Elements of

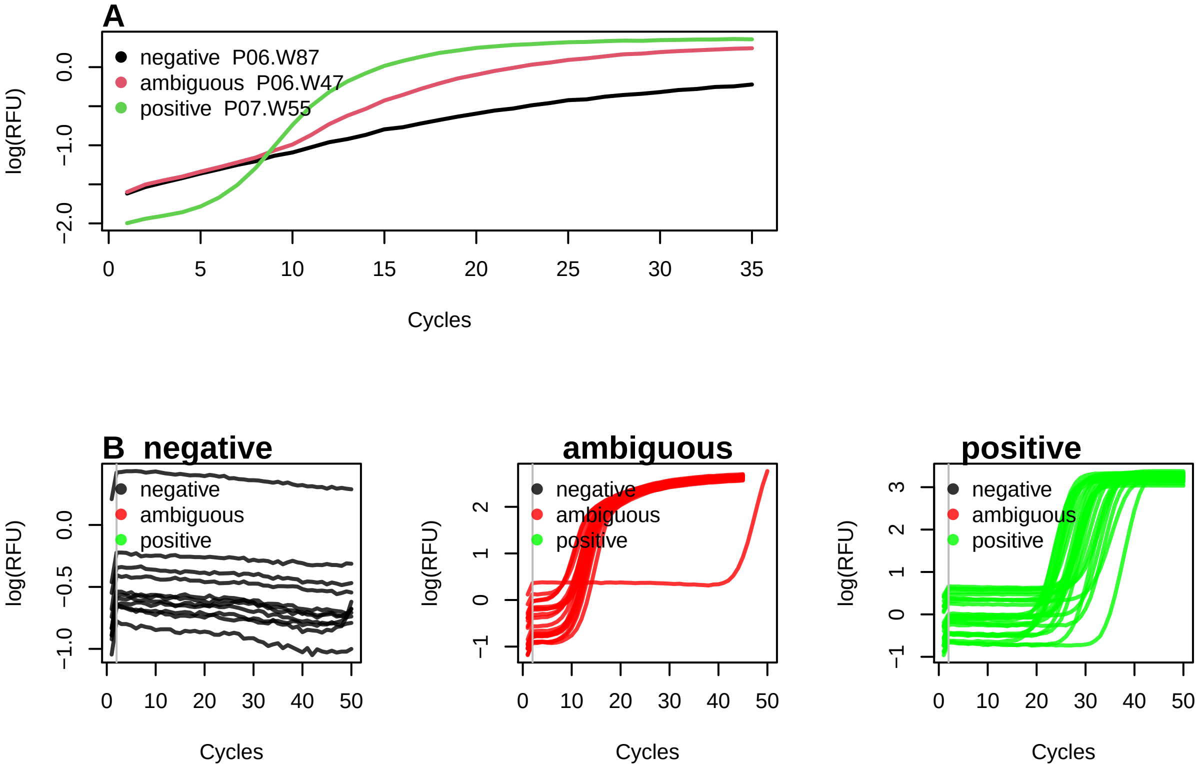

Figure 1: Examples of negative, ambiguous and positive amplification curves. A) A negative (black), ambiguous (red) and positive (green) amplification curve were selected from the ‘htPCR‘ data set. The negative amplification curve is non-sigmoid and has a positive trend. The ambiguous amplification curve is similar to a sigmoidic amplification curve, but shows a positive slope in ground phase (cycle 1 $\rightarrow$ 5). The positive amplification curve (green) is sigmoid. It starts with a flat baseline (cycle 5 $\rightarrow$ 25). This is followed by the exponential phase (cycle 5 $\rightarrow$ 25) and ends in a flat plateau phase (cycle 26 $\rightarrow$ 35). B) Amplification curves of the ‘vermeulen1‘ data set were divided into groups with *negative*, *ambiguous* and *positive* classification. Negative amplification curves have a low signal level. Interesting is the spontaneous increase (probably due to a sensor calibration) in cycles 1 to 2 followed by a linear signal decrease. In principle, the ambiguous amplification curves have a sigmoid curve shape. However, the plateau phase is fairly broad. When different qPCR users were asked what they find ambiguous, they responded that there is an additional change in the slope between cycles 15 to 25. This made some believe that the reaction is not valid. One of the ambiguous amplification curves begins to rise sharply at Cycle 45. The positive amplification curves have a characteristic sigmoid curve shape.

the data mining process are the preprocessing of the data, the description of the data, the exploration of the data and the search for connections and causes.

The availability of classified amplification curve data sets and technologies for the classification of amplification curves is of high importance to train and validate models. This is dealt with in the **online supplement**.

For machine learning, the type of learning task is the first thing that needs to be defined. The learning task can be a classification, clustering or regression problem. Next, suitable algorithms can be selected depending on the task. In the case of classification problems, it is attempted to predict a *discrete valued* output. The labels ($y$) are usually categorical and represent a finite number of classes (e. g. "negative", "positive" $\rightarrow$ binary classification). With regression tasks, it is attempted to predict a *continuously valued* output. Clustering is primarily about forming groups (clusters) based on their similarities. More examples are presented in the **online supplement**.

In contrast, machine learning uses instructions and data in software modules to create models that can be used to make predictions on novel data. In machine learning, the human being is much less necessary in the entire process. Processes (algorithms) are used to create models with tunable parameters. These models automatically adapt their performance to the information (predictors) from the data. Well-known examples of machine learning technologies are Decision Trees (DT), Boosting, Random Forests (RF), Support Vector Machines (SVM), generalized linear models (GLM), logistic regression (LR) and deep neural networks (DNN) (Lee 2010). The three following concepts of machine learning that are frequently described in the literature are *Supervised learning*, *Unsupervised learning* and *Reinforcement Learning* which are described in detail in the **online supplement**.

### 3 Why is there a need for the `PCRedux` software?

The classification of an amplification curve is feasible using bioanalytical methods such as melting curve analysis (Rödiger, Böhm, and Schimke 2013) or electrophoretic separation (Westermeier 2004). However, this is not always possible or desirable.

- Melting curve analysis is used in some qPCRs as a post-processing step to identify samples which contain the specific target sequence (*positive*) based on a specific melting temperature. However, some detection probe systems like hydrolysis probes do not permit such classification. Moreover, nucleic acids with similar biochemical properties but different sequences may have the same melting temperature.
- An electrophoretic separation (classification of target DNA sequences by size and quantity) often requires too much effort for experiments with high sample throughput.
- There are mathematical qPCR analysis algorithms such as `linreg` (Ruijter et al. 2009) that require information on whether an amplification curve is negative or positive for subsequent calculation.
- Raw data of amplification curves can be fitted with sigmoid functions. Sigmoid functions are non-linear, real-valued, have an S-shaped curvature (Figure 3) and can be differentiated (e. g., first derivative maximum, with one local minimum and one local maximum). With the obtained model, predictions can be made. For example, the position of the second derivative maximum can be calculated from this (section 4). In the context of amplification curves, the second derivative maximum is commonly used to describe the relationship between the cycle number and the PCR product formation (section 4). All software assume that the amplification resembles a sigmoid curve shape (ideal positive amplification reaction), or a flat low line (ideal negative amplification reaction). For example, Ritz and Spiess (2008) published the `qpcR` `R` package that contains functions to fit several multi-parameter models. This includes the five-parameter Richardson function (Richards 1959) (Equation 3). The `qpcR` package (Ritz and Spiess 2008) contains an amplification curve test via the `modlist()` function. The parameter `check="uni2"` offers an analytical approach, as part of a method for the kinetic outlier detection. It tries to checks for a sigmoid structure of the amplification curve. Then, `modlist()` tests for the location of the first derivative maximum and the second derivative maximum. However, multi-parameter functions fit "successful" in most cases including noise and give false positive results. This will be shown in later sections. This is exemplary shown in later sections in combination with the `amptester()` [`chipPCR`] function (Rödiger, Burdukiewicz, and Schierack 2015), which uses fixed thresholds and frequentist

inference to identify amplification curves that exceed the threshold ($\mapsto$ classified as positive). However, the analysis can also lead to false-positive classifications as exemplified in the example below and in Figure 2. Therefore, additional classification concepts would be beneficial.

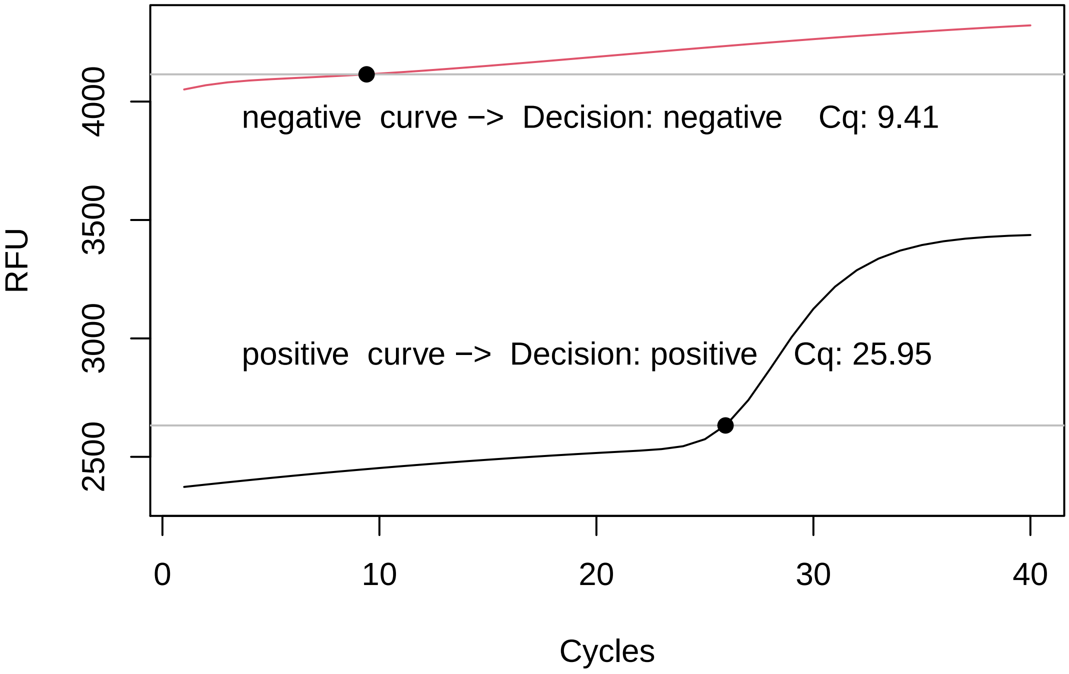

Figure 2: Incorrect model adjustment for amplification curves. A positive (black), and a negative (red) amplification curve were randomly selected from the 'RAS002' data set. The positive amplification curve has a baseline signal of about 2500 RFU (Raw Fluorescene Unit) and has a definite sigmoidal shape. The negative amplification curve has a baseline signal of approx. 4200 RFU, but only moderately positive slope (no sigmoidal shape). A logistic function with seven parameters ('l7') has been fitted to both amplification curves. A Cq value of 25.95 was determined for the positive amplification curve. The negative amplification curve had a Cq value of 9.41. However, it can be seen that the latter model fitting is not appropriate for calculating a trustworthy Cq value. An automatic calculation without user control would give a false-positive result. Note: This plot was shown in linear scale to demonstrate typical pitfalls.

- The analysis and classification of sigmoid data (e. g., quantitative PCR) is a manageable task if the data volume is low, or dedicated analysis software is available. An example for a low number of amplification curves is shown in Figure 4A. All 65 curves exhibit a sigmoid curve shape. It is trivial to classify them as positive by hand. In contrast, the vast number of amplification curves in Figure 4B is barely manageable with a reasonable effort by simple visual inspection. These data originate from a high-throughput experiment that encompasses in total 8858 amplification curves of which only 200 are shown. A manual analysis of the data is time-consuming and prone to errors. Even for an experienced user, it is difficult to classify the amplification curves unambiguously and reproducible as will be later shown in subsection 4.3.
- qPCRs are performed in thermo-cyclers, which are equipped with a real-time monitoring technology. There are numerous commercial manufactures producing thermo-cyclers (Table 4). An example for a thermo-cycler that originated in a scientific project is the VideoScan technology (Rödiger, Schierack, et al. 2013). Most of the thermo-cyclers have a thermal block with wells at certain positions. Reaction vessels containing the PCR mix are inserted into the wells. There are also thermo-cyclers that use capillary tubes that are heated and cooled by air (e. g., Roche Light Cycler 1.0). The thermo-cycler raises and lowers the temperature in the reaction vessels in discrete, pre-programmed steps so that PCR cycling can take place. Instruments with a real-time monitoring functionality have sensors to measure changes of the fluorescence intensity in the reaction vessel. All thermo-cycler systems use software to processes the amplification curves. Plots of the fluorescence observations versus cycle number obtained from two different qPCR systems are shown in Figure 4A and B. The thermo-cyclers

produce different amplification curve shapes even with the same sample material and PCR mastermix because of their technical design, sensors, and software. These factors need to be taken into account during the development of analysis algorithms.

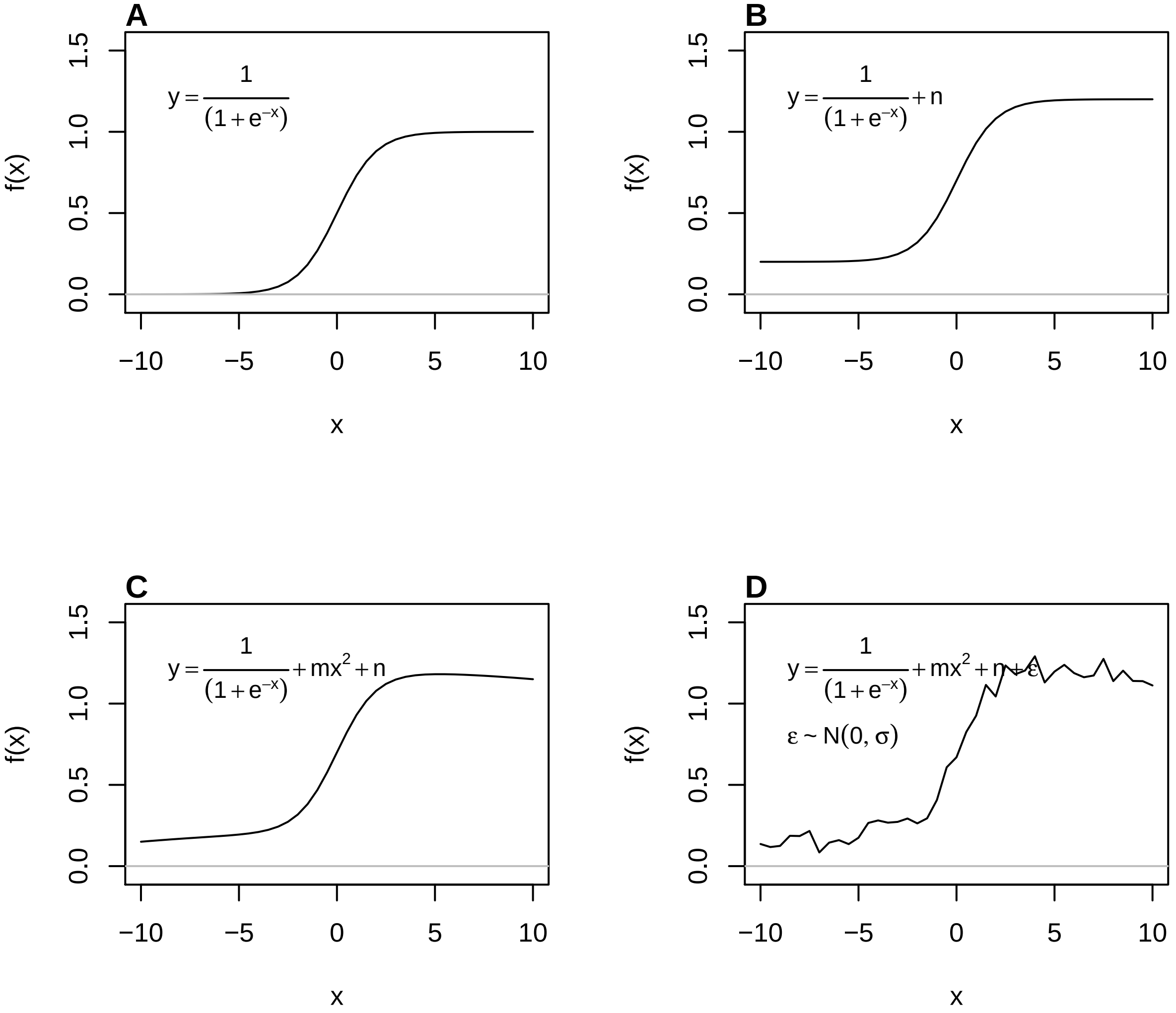

Figure 3: A) Model function of a one-parameter sigmoid function. B) Model function of a sigmoid function with an intercept $n = 0.2$ RFU (shift in base-line). C) Model function of a sigmoid function with an intercept ($n$ ~0.2 RFU) and a square portion $m * x^2, m = -0.0005, n = 0.2RFU$ (hook-effect-like). D) Model function of a sigmoid function with an intercept ($n$) and a square portion of $m * x^2$ and additional noise $\epsilon$ (normal distributed, $\mu = 0.01, \sigma = 0.05$). Note: This plot was shown in linear scale to demonstrate typical pitfalls.

There are several open source and closed source software tools for the analysis of qPCR data (Pabinger et al. 2014). The software packages deal for example with challenges like missing values and non-detects (McCall et al. 2014), quantification cycle estimation (Ritz and Spiess 2008; Ruijter et al. 2013), relative gene expression analysis (Dvinge and Bertone 2009; Pabinger et al. 2009; Neve et al. 2014) and data analysis pipelines (Pabinger et al. 2009; Ronde et al. 2017; Mallona, Weiss, and Egea-Cortines 2011; Mallona et al. 2017). More information can be found in the **online supplement**.

However, a bottleneck of qPCR data analysis is the lack of predictors and software to build classifiers for amplification curves. A classifier herein refers to a vector of **interpretable** predictors that can be used to distinguish the amplification curves only by their shape. A predictor, also referred to as *feature*, is an entity that characterizes an object. A few potential predictors for amplification curves are described in the literature.

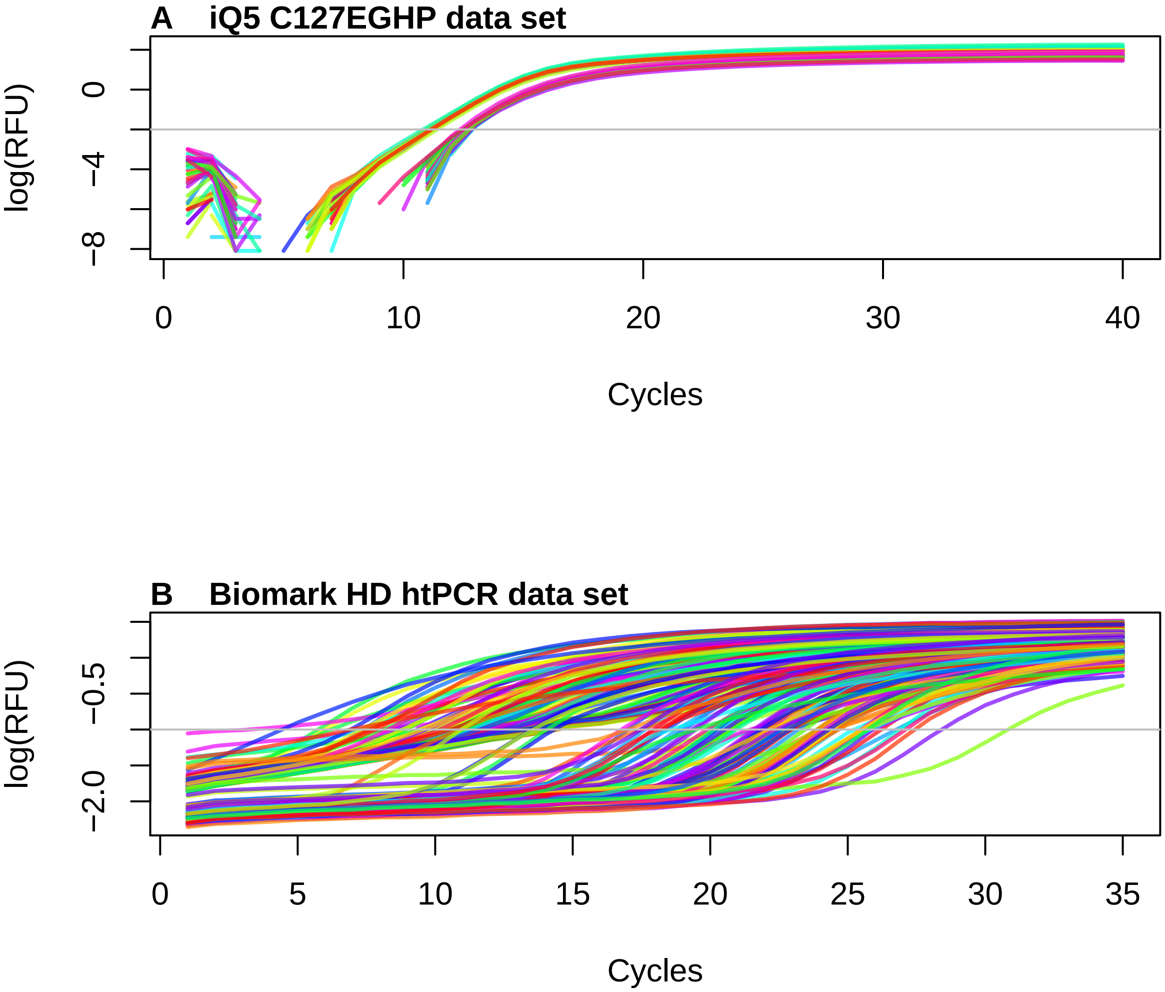

Figure 4: Amplification curve data from an iQ5 (Bio-Rad) thermo-cycler and a high throughput experiment in the Biomark HD (Fluidigm). A) The ‘C127EGHP‘ data set with 64 amplification curves was produced in a conventional thermo-cycler with a 8 x 12 PCR grid. B) The ‘htPCR‘ data set, which contains 8858 amplification curves, was produced in a 95 x 96 PCR grid. Only 200 amplification curves are shown. In contrast to ‘A)‘ have all amplification curves in ‘B)‘ a stong off-set (intercept) between -2.5 and 0 log(RFU). This needs proper baselineing.

### 4 Principles of Amplification Curve Data Analysis and Predictor Calculation

The shape of a positive amplification curve is in most cases sigmoidal. Many factors such as the sample quality, qPCR chemistry, and technical problems (e. g., sensor errors) contribute to various curve shapes (Ruijter et al. 2014). The curvature of the amplification curve can be used as a quality measure. For example, fragmentation, inhibitors and sample material handling errors during the extraction can be identified. The kinetic of fluorescence emission is proportional to the quantity of the synthesized DNA. Typical amplification curves have three phases.

1. **Ground phase**: This phase occurs during the first cycles of the PCR, where the fluorescence emission is in most cases flat. Here, noise but no product formation is detected by the sensor system and the PCR product signal is an insignificantly small component of the total signal. This is often referred to as base-line or background signal. Apparently, there is only a phase shift or no signal at all, primarily due to the limited sensitivity of the instrument. Even in a perfect PCR reaction (double amplification per cycle), qPCR instruments cannot detect the fluorescence signal from the amplification. Fragmentation, inhibitors and sample handling errors would result in a prolonged ground phase. Nevertheless, this may indicate some typical properties of the qPCR system or probe system. In many instruments, this phase is used to determine the base-line level for the calculation of the Cycle threshold (Ct). The Ct value is considered statistically relevant as an increase outside of the noise range (threshold) when coming from the amplicon. In some qPCR systems, a flat amplification signal is expected in this phase. Slight deviations from this trend are presumably due to changes (e. g., disintegration of probes) in the fluorophores. Background correction algorithms are often used here to ensure that flat amplification curves without slope are generated. However, this can result in errors and inevitably leads to a loss of information via the waveform of the raw data (Nolan, Hands, and Bustin 2006). The slope, level and variance of this phase can serve as predictors.
2. **Exponential phase**: This phase follows the ground phase and is also called *log-linear phase*. It is characterized by a strong increase of the emitted fluorescence as the DNA amount roughly doubles in each cycle under ideal conditions and when the amount of the synthesized fluorescent labeled PCR product is high enough to be detected by the sensor system. This phase is used for the calculation of the quantification point (Cq) and curve specific amplification efficiency. The most important measurement from qPCRs is the Cq, which signifies the PCR cycle for which the fluorescence exceeds a `threshold value`. However, there is an ongoing debate as to what a significant and robust threshold value is. An overview and performance comparison of Cq methods is given in Ruijter et al. (2013). There are several mathematical methods to calculate the Cq.
    - The 'classical' threshold value (cycle threshold, Ct) is the intersection between a manually defined straight horizontal line with the quasi-linear phase in the exponential amplification phase (Figure 6A & B). This simple to implement method requires that amplification curves are properly baselined prior to analysis. The Ct method makes the assumption that the amplification efficiency (~ slope in the log-linear phase) is equal across all compared amplification curves (Ruijter et al. 2013). Evidently, this is not always case as exemplified in Figure 5C. The Ct method is widely used presumably due to the familiarity of users with this approach (e. g., chemical analysis procedures). However, this method is statistically unreliable (Ruijter et al. 2013; Spiess et al. 2015, 2016). Moreover, the Ct method gives no stable in predictions if different users are given the same data set to be analyzed. *Therefore, this method is not used within the `PCRedux` package.*
    - Another Cq method uses the maximum of the second derivative (SDM) (Rödiger, Burdukiewicz, et al. 2015) (Figure 6C). In all cases, the Cq value can be used to calculate the concentration of target sequence in a sample (low Cq →high target concentration). In contrast, negative or ambiguous amplification curves loosely resemble noise. This noise may appear linear or exhibit a curvature similar to a specific amplification curve (Figure 1). This however, may result in faulty interpretation of the amplification curves. Fragmentation, inhibitors and sample handling errors would decrease the slope of the amplification curve (Spiess, Feig, and Ritz 2008; Ritz and Spiess 2008). The slope and its variation can be considered as predictors. Since the Cq depends on the initial template amount and amplification efficiency, there seemingly is no immediate use of the

Cq as an predictor.

3. **Plateau phase**: This phase follows the exponential phase and is a consequence of the exhaustion of limited reagents (incl. primers, nucleotides, enzyme activity) in the reaction vessel, limiting the amplification reaction, so that the theoretical maximum amplification efficiency (doubling per cycle) no longer prevails. This turning point, and the progressive limitation of resources, finally leads to a plateau. In the plateau phase, there is in some cases a signal decrease called *hook effect* (section 3 and (Barratt and Mackay 2002; Isaac 2009; Burdukiewicz et al. 2018)). The slope (*hook effect*), level and variation can be considered as predictors.

If the amplification curve has only a slight positive slope and no perceptible/measureable exponential phase, it can be assumed that the amplification reaction did not occur (Figure 5B). Causes may include poor specificity of the PCR primers (non-specific PCR products), degraded sample material, degraded probes or detector failures. If a lot of input DNA is present in a sample, the amplification curve starts to increase in early PCR cycles (1 - 12 cycles). Some PCR devices have a software that corrects this feature without rechecking, resulting in an amplification curve with a negative trend.

The discussed phases are considered as regions of interest (ROI). As an example, the *ground phase* is in the head area, while the *plateau phase* is in the tail area. The *exponential phase* is located between these two ROIs.

The amplification curve shape, the amplification efficiency and the Cq value are important measures to judge the outcome of a qPCR reaction. In all phases of PCR, the curves should be smooth. Possible artifacts in the curves may be due to unstable light sources from the instrument or problems during sample preparation, such as the presence of bubbles in the reaction vessel, incorrectly assigned dye detectors, errors during the calibration of dyes for the instrument, errors during the preparation of the PCR master mix, sample degradation, lack of a sample in the PCR, too much sample material in the PCR mix or a low detection probe concentration (Ruijter et al. 2009, 2014; Spiess et al. 2015). Smoothing and filtering cause alterations to the raw data that affects the Cq value and the amplification efficiency.

Most commercial qPCR systems do not display the raw data of the amplification curves on the screen. Instead, raw data are often processed by the instrument software to remove fluorophore-specific effects and noise in all ROIs. Commonly employed preprocessing step of qPCR is smoothing and filtering to remove noise, where the latter can have different causes (Spiess et al. 2015).

The ordinate often does not display the measured fluorescence, but rather the change in fluorescence per cycle ($\Delta RFU = RFU_{cycle+1} - RFU_{cycle}$). Some qPCR systems display periodicity in the amplification curve data, thereby exposing the risk of introducing artificial shifts in the Cq values (Spiess et al. 2016).

In particular the cycle threshold method (Ct method) (section 4) is affected by these factors (Spiess et al. 2015, 2016). Therefore, it is advisable to clarify in advance, which processing steps the amplification curves have been subjected to. Failure to do so may result in misinterpretations and incorrect amplification curve fitting models (Nolan, Hands, and Bustin 2006; Rödiger, Burdukiewicz, et al. 2015; Rödiger, Burdukiewicz, and Schierack 2015; Spiess et al. 2015).

#### 4.1 Data Analysis Functions of the `PCRedux` Package

##### 4.1.1 Helper Functions of the `PCRedux` Package

The `PCRedux` package contains functions for analyzing amplification curves. These are distinguished into helper functions and analysis functions. The details about the: - `performeR()` - Performance Analysis for Binary Classification and - `qPCR2fdata()` - A Helper Function to Convert Amplification Curve Data to the `fdata` format can by found in the **online supplement**.

##### 4.1.2 `pcrfit_single()` and `encu()`- Predictor calculation from an Amplification Curve

The following sections give a concise description of the algorithms used to calculate predictor vectors by the `pcrfit_single()` function. Based on considerations and experience, the algorithms of the `pcrfit_single()` function are restricted to ROIs (Figure 5) to calculate specific predictors.

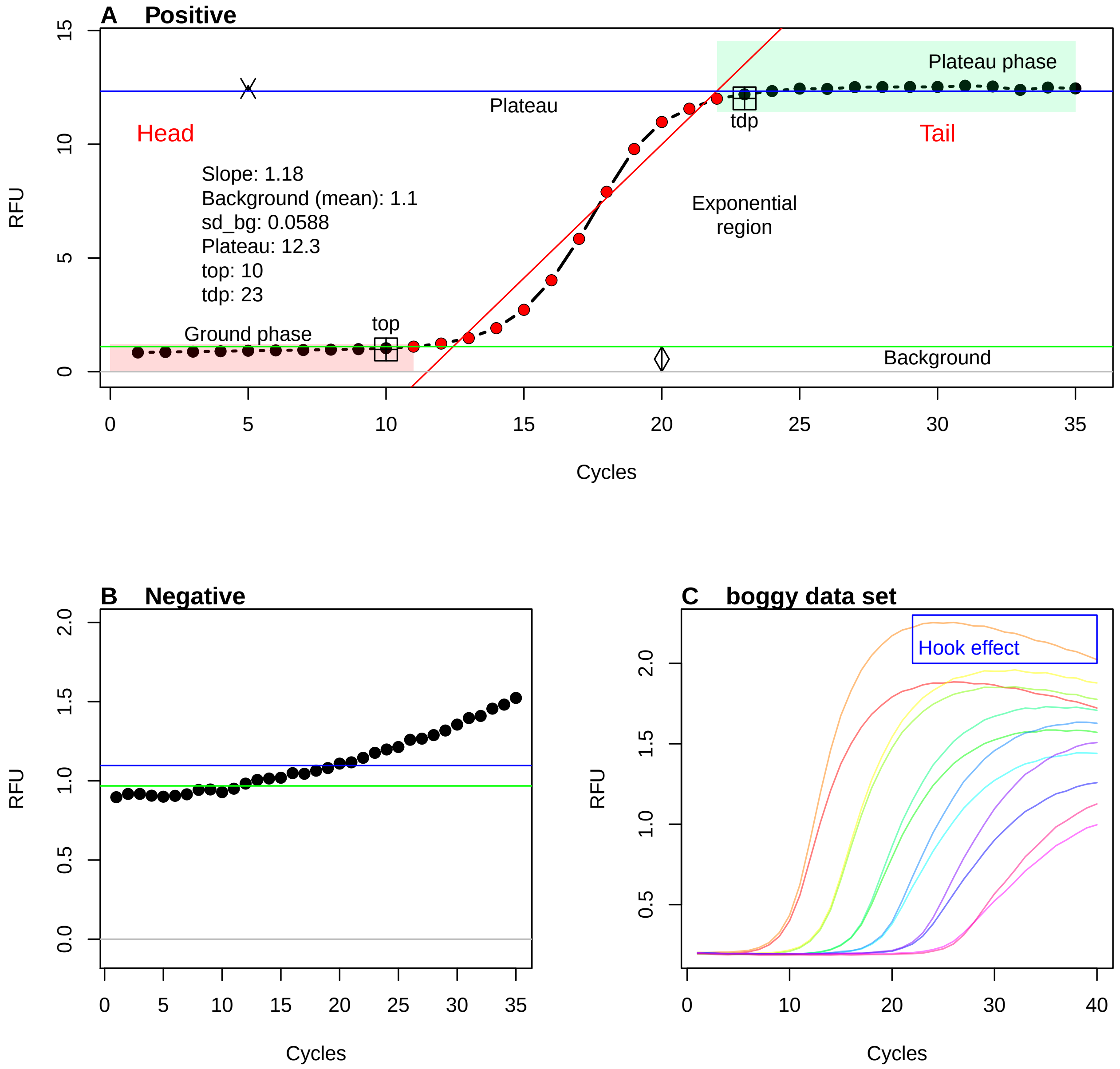

Figure 5: Phases of amplification curves as Region of Interest (ROI). For amplification curves, the fluorescence signal (RFU, relative fluorescence units) of the reporter dye is plotted against the cycle number. Positive amplification curves possess three ROIs: ground phase, exponential phase and plateau phase. These ROIs can be used to determine predictors such as the takedown point (‘tdp‘) or the standard deviation within the ground phase (‘sd_bg‘). The exponential range (red dots) is used to determine the Cq values and amplification efficiency (not shown). A linear regression model (red) can be used to calculate the slope in this region. B) PCRs without amplification reaction usually show a flat (non-sigmoides) signal. C) The exponential phase of PCR reactions can vary greatly depending on the DNA starting quantity and other factors. Amplification curves that appear in later cycles often have a lower slope in the exponential phase.

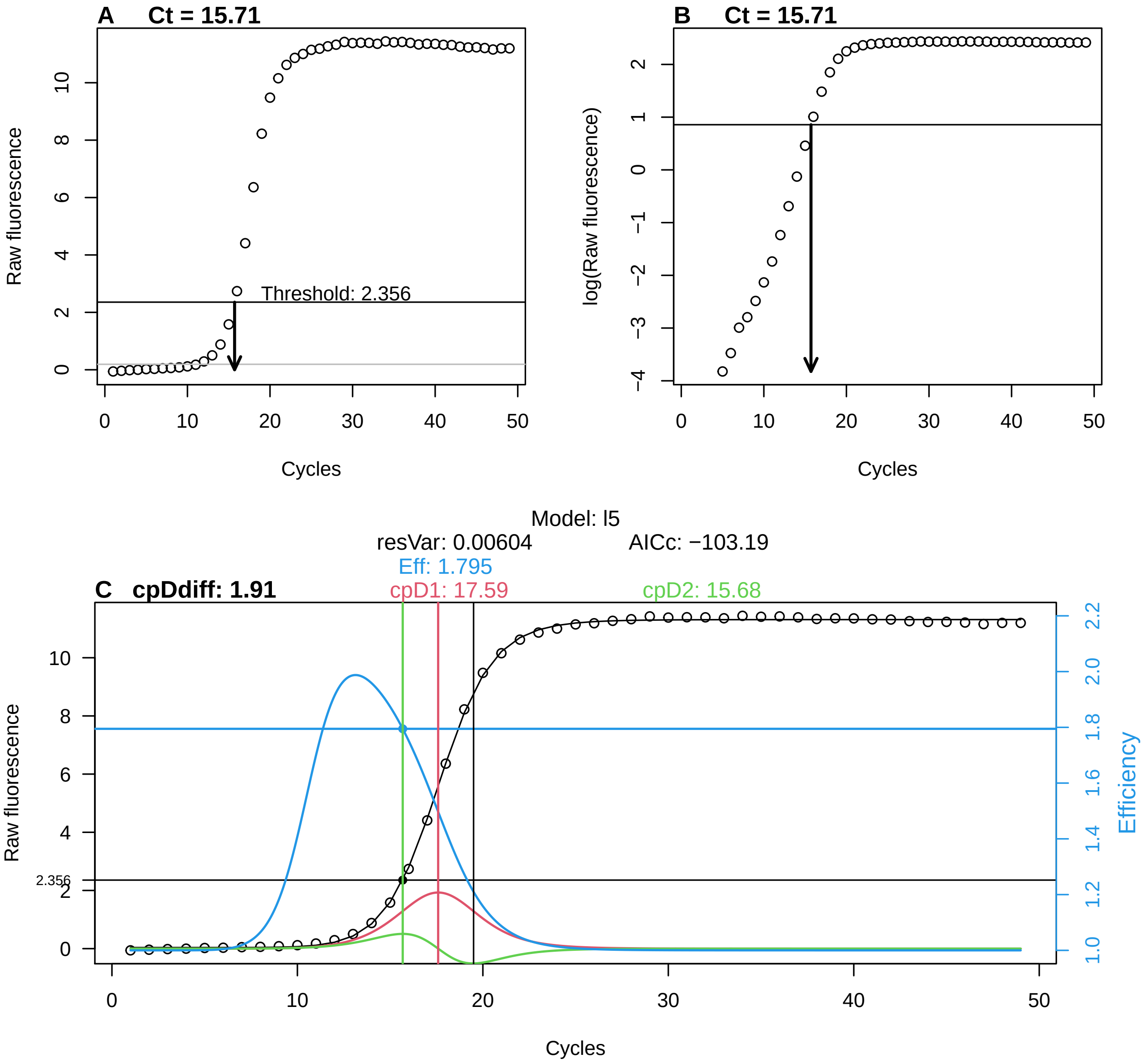

Figure 6: Frequently used methods for the analysis of quantification points. A) The amplification curve is intersected by a gray horizontal line. This is the background signal ($3\sigma$) determined from the *68-95-99.7 rule* from the fluorescence emission of cycles 1 to 10. The black horizontal line is the user-defined threshold (Ct value) in the exponential phase. Based on this, the cycle at which the amplification curve differs significantly from the background is calculated. B) The amplification curve can also be analyzed by fitting a multi-parametric model (black line, five parameters). The red line is the first derivative of the amplification curve with a maximum of 17.59 cycles. The first derivative maximum ('cpD1') is used as a quantification point (Cq value) in some qPCR systems. The green line shows the second derivative of the amplification curve, with a maximum at 15.68 cycles a minimum at 19.5 cycles. The maximum of the second derivative ('cpD2') is used as the Cq value in many systems. The blue line shows the amplification efficiency estimated from the trajectory of the exponential region. The 'Eff' value of 1.795 means that the amplification efficiency is approximately 89%. 'cpDdiff' is the difference between the first and second derivative maximum ($cpDdiff = cpD1 - cpD2$).

The `encu()` function is a wrapper for the `pcrfit_single()` function. `encu()` can be used to process large records of amplification curve data arranged in columns. The progress of processing is displayed in the form of a progress bar and the estimated run-time. Additionally, `encu()` allows to specify which monitoring chemistry (e. g., DNA binding dye, sequence specific probes) and which thermo-cycler was used. Ruijter et al. (2014) demonstrated that the monitoring chemistry and the type of input DNA (single stranded, double stranded) are important when analysing qPCR data, because they have an influence on the shape of the amplification curve. For simplicity, the documentation will describe the `pcrfit_single()` only.

The underlying hypotheses and concepts of the predictors are formulated and supported by *exemplary applications*. Different representative data sets were used to support a concept or predictors. For example, the `RAS002` data set represents a typical qPCR. This means that the positive amplification curves start with a flat plateau phase and then transition into the sigmoid shape with a plateau. The negative amplification curves display no significant peculiarities. For both positive and negative amplification curves, there is a shift from the origin. The `htPCR` data set serves as a problem example in several places, since it contains many observations (amplification curves from high-throughput experiments). In addition, the amplification curves have a high diversity of curve shapes that cannot be uniquely and reproducibly classified even by experienced users. Other data sets are used in the documentation, but these are not discussed in detail.

To underscore the usability of the algorithms and their predictors, 3302 observations (471 negative amplification curves, 2831 positive amplification curves) from the `batsch1`, `boggy`, `C126EG595`, `competimer`, `dil4reps94`, `guescini1`, `karlen1`, `lievens1`, `reps384`, `rutledge`, `testdat`, `vermeulen1`, `VIMCFX96_60`, `stepone_std`, `RAS002`, `RAS003`, `HCU32_aggR` and `lc96_bACTXY` were analyzed with the `encu()` function and the results (predictors) were combined in the file **`data_sample.rda`**. Users of this function should independently verify and validate the results of the methods for their own applications.

A new data set called `data_sample_subset_balanced` has been compiled from the `data_sample` data set for some of the applications. Selection criteria included:

- both positive and negative amplification curves had to be included in a similar ratio,
- there should not be a dominating thermal cycler platform,
- the amplification curves should represent typical amplification curves (subjective criterion). The compilation of the data sets `batsch1`, `HCU32_aggR`, `lc96_bACTXY`, `RAS002`, `RAS003` and `stepone_std` met this requirement satisfactorily.

```
## [1] 651  62
```

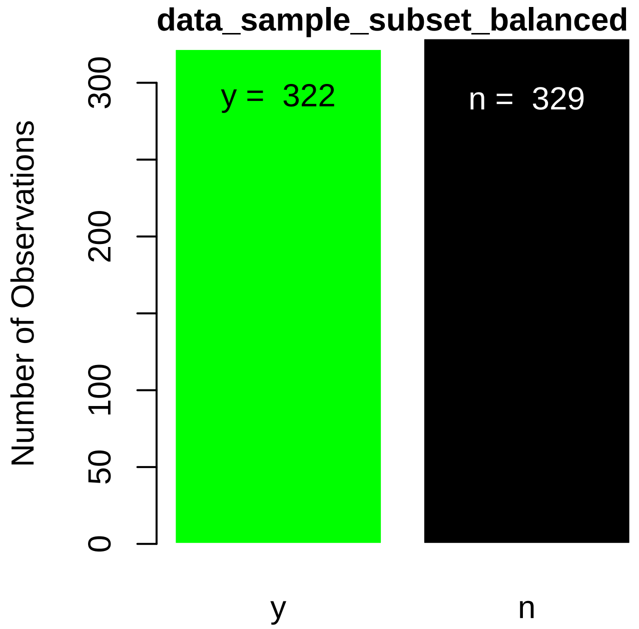

For the comparison of predictors, the data set was enlarged. Selection criteria for the data sets were comparatively less stringent.

```
data_sample_subset <- data_sample[data_sample$dataset %in% c("stepone_std",
                                                              "RAS002", "RAS003",
                                                              "lc96_bACTXY",
                                                              "C126EG595",
                                                              "dil4reps94",
                                                              "testdat",
                                                              "boggy"), ]

# Dimension of data_sample_subset
dim(data_sample_subset) ## Observations predictors
```

```
## [1] 979  62
```

```
# Show the counts of negative and positive amplification
# curves in a bar plot
# Build a contingency table of the counts at each
# combination of factor levels.

dec_table<- table(data_sample_subset[["decision"]])
barplot(dec_table, ylab = "Number of Observations", col = c("green", "black"),
        border = "white")
text(c(0.7, 1.9), rep(min(dec_table) *0.9, length(dec_table)),
     c(paste("y = ", dec_table[1]), paste("n = ", dec_table[2])),
     col = c("black", "white"))
mtext("data_sample_subset", cex = 1, side = 3, adj = 0, font = 2,
      las = 0)
```

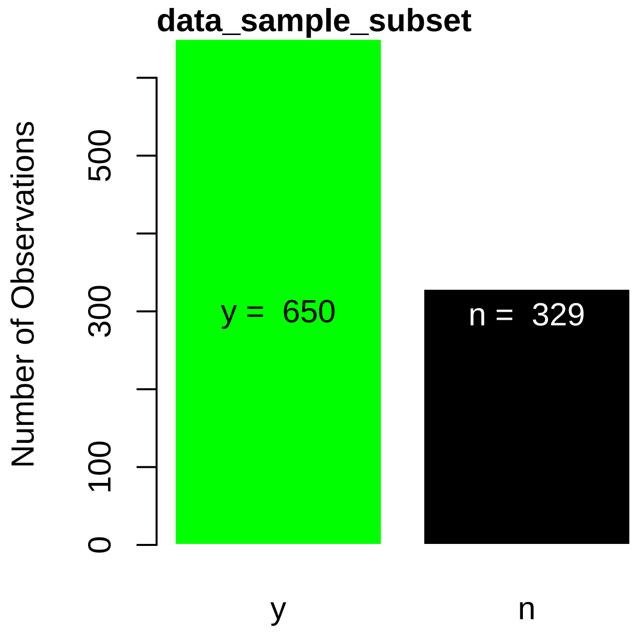

The goal is to demonstrate the basic functionality of the algorithms for predictor calculation. Similar concepts are presented in groups. The algorithms are divided into the following broad categories:

- algorithms that determine slopes, signal levels,
- algorithms that determine turning points and
- algorithms that determine areas.

The algorithms in

- `earlyreg()` (subsubsection 4.1.9),
- `head2tailratio()` (subsubsection 4.1.10),
- `hookreg()` & `hookregNL()` (subsubsection 4.1.11) and
- `mblrr()` (subsubsection 4.1.12),
- `autocorrelation_test()` (subsubsection 4.1.13)

were implemented as standalone functions to make them available for other applications.

The output below shows the predictors and their data type (`num`, numeric; `int`, integer; `Factor`, factor; `logi`, boolean) that were determined with the `pcrfit_single()` function.

```
library(PCRedux)
# Calculate predictor vector of column two from the RAS002 data set.
str(pcrfit_single(RAS002[, 2]))
```

```
## 'data.frame':    1 obs. of  90 variables:
##  $ cpD1                  : num 28.3
##  $ cpD2                  : num 25.6
##  $ cpD2_approx           : num 26
##  $ cpD2_ratio            : num 0.986
##  $ eff                   : num 1.48
##  $ sliwin                : num 1.37
##  $ cpDdiff               : num 2.67
##  $ loglin_slope          : num 118
##  $ cpD2_range            : num 4.62
##  $ top                   : num 25
##  $ f.top                 : num 160
##  $ tdp                   : num 33
##  $ f.tdp                 : num 907
##  $ bg.stop               : num 15
##  $ amp.stop              : num 40
##  $ b_slope               : num -13.6
##  $ b_model_param         : num -13.6
##  $ c_model_param         : num -62.9
##  $ d_model_param         : num 818
##  $ e_model_param         : num 26
##  $ f_model_param         : num 3.17
##  $ f_intercept           : num 3.17
##  $ k1_model_param        : num 10.3
##  $ k2_model_param        : num -0.133
##  $ convInfo_iteratons    : int 13
##  $ qPCRmodel             : Factor w/ 1 level "l7": 1
##  $ qPCRmodelRF           : Factor w/ 1 level "l7": 1
##  $ minRFU                : num -76.5
##  $ maxRFU                : num 1016
##  $ init2                 : num 0.00846
##  $ fluo                  : num 206
##  $ slope_bg              : num 22.6
##  $ intercept_bg          : num -102
##  $ sigma_bg              : num 15.7
##  $ sd_bg                 : num 0.055
##  $ head2tail_ratio       : num -0.0149
##  $ mblrr_slope_pt        : num 20.2
##  $ mblrr_intercept_bg    : num -38.6
```

```
##  $ mblrr_slope_bg       : num 6.96
##  $ mblrr_cor_bg         : num 0.91
##  $ mblrr_intercept_pt   : num 237
##  $ mblrr_cor_pt         : num 0.942
##  $ polyarea             : num 5960
##  $ peaks_ratio          : num 0.0164
##  $ autocorrelation      : num 0.725
##  $ cp_e.agglo           : int 6
##  $ cp_bcp               : int 5
##  $ amptester_shapiro    : num 0
##  $ amptester_lrt        : num 1
##  $ amptester_rgt        : num 1
##  $ amptester_tht        : num 1
##  $ amptester_slt        : num 1
##  $ amptester_polygon    : num -10025
##  $ amptester_slope.ratio: num 0.00386
##  $ hookreg_hook         : num 0
##  $ hookreg_hook_slope   : num 0
##  $ hookreg_hook_delta   : num 0
##  $ central_angle        : num 0.999
##  $ number_of_cycles     : int 40
##  $ direction            : num 0
##  $ range                : num 1093
##  $ polyarea_trapz       : num 12363
##  $ cor                  : num 0.898
##  $ res_coef_pcrfit.b    : num -14.1
##  $ res_coef_pcrfit.c    : num 30.8
##  $ res_coef_pcrfit.d    : num 1024
##  $ res_coef_pcrfit.e    : num 28.6
##  $ fitAIC               : num 408
##  $ fitIter              : int 13
##  $ segment_x            : num 0
##  $ segment_U1.x         : num 0
##  $ segment_U2.x         : num 0
##  $ segment_psi1.x       : num 0
##  $ segment_psi2.x       : num 0
##  $ sumdiff              : num 0.95
##  $ poly_1               : num 321
##  $ poly_2               : num 2238
##  $ poly_3               : num 899
##  $ poly_4               : num -32
##  $ window_Win_1         : num 29.8
##  $ window_Win_2         : num 7.73
##  $ window_Win_3         : num 7.17
##  $ window_Win_4         : num 6.64
##  $ window_Win_5         : num 5.1
##  $ window_Win_6         : num 16.6
##  $ window_Win_7         : num 122
##  $ window_Win_8         : num 125
##  $ window_Win_9         : num 34.4
##  $ window_Win_10        : num 10.3
##  $ sd_plateau           : num 0.0387
```

##### 4.1.3 Amplification Curve Preprocessing

The `pcrfit_single()` function performs preprocessing steps before each calculation, including checking whether an amplification curve contains missing values. Missing values (NA) are measuring points in a data set where no measured values are available or have been removed arbitrarily. NAs may occur if no measurement has been carried out (e. g., defective detector) or lengths of the vectors differ (number of cycles) between the observations. Such missing values are automatically imputed by spline interpolation as described in Rödiger, Burdukiewicz, and Schierack (2015).

Values of an amplification curve are normalized to their 99% quantile or rare cases to the maximum for many calculations. The normalization is used to equalize the amplitudes differences of amplification curves from thermo-cyclers (sensor technology, software processing) and detection chemistries. To compare amplification curves from different thermo-cyclers, the values should always be scaled systematically using the same method. Although there are other normalization methods (e. g., minimum-maximum normalization, see Rödiger, Burdukiewicz, and Schierack (2015), the normalization by the 99% quantile preserves the information about the level of the background phase. A normalization to the maximum is not used to avoid strong extenuation by outliers. The data in Figure 16D show that the `maxRFU` values after normalization are approximately 1. There is no statistical significant difference between `maxRFU` values of positive and negative amplification curves.

Selected algorithms of the `pcrfit_single()` function use the `CPP()` [`chipPCR`] function to preprocess (e. g., base-lining, smoothing, imputation of missing values) the amplification curves. Further details are given in Rödiger, Burdukiewicz, and Schierack (2015). Until package version 0.2.6-4 was the `visdat_pcrfit()` part of the package. `visdat_pcrfit()` was used for visualizing the content of data from an analysis with the `pcrfit_single()` function. There are other more powerful packages such as `visdat` by Tierney (2017), `assertr` by Fischetti (2019) and `xray` by Seibelt (2017).

During the analysis, several values are determined to describe the amplitude of an amplification curve. The resulting potential predictors are `minRFU` (minimum of the amplification curve, which is determined at the 1% quantile to minimize the influence of outliers), `init2` (the initial template fluorescence from an exponential model) and `fluo` (raw fluorescence value at the second derivative maximum). The `minRFU`, `init2` and `fluo` values differ significantly between negative and positive amplification curves (Figure 16C, E & F).

##### 4.1.4 Handling of Missing Predictors

Missing values (NA) can occur if a calculation of a predictor is impossible (e. g., if a logistic function cannot be adapted to noisy raw data). The lack of a predictor is nevertheless an useful information (no predictor calculate $\mapsto$ amplification curves deviate from sigmoid shape). The NAs were left unchanged in the `PCRedux` package up to version 0.2.5-1. Since version 0.2.6 the NAs are replaced by numerical values (e. g., total number of cycles) or factors (e. g., *lNA* for non-fitted model). Under the term "imputation", there are a number of procedures based on statistical methods (e. g., neighboring median, spline interpolation) or on user-defined rules (Williams 2009; Cook and Swayne 2007; Hothorn and Everitt 2014). Rules are mainly used in the functions of `PCRedux` to relieve the user from the decision as to how to deal with missing values. For example, slope parameters of a model are set to zero when it cannot be determined. The disadvantage is that rules do not necessarily concur to real world values.

##### 4.1.5 Multi-parametric Models for Amplification Curve Fitting

Both the `pcrfit_single()` function and the `encu()` function use four multi-parametric models based on the findings of Spiess, Feig, and Ritz (2008) and Ritz and Spiess (2008). The `pcrfit_single()` function starts by adjusting a seven-parameter model since this adapts *easier* and more frequent to a data set (Figure 7).

- **l7**:

$$f(x) = c + k1 \cdot x + k2 \cdot x^2 + \frac{d - c}{(1 + exp(b(log(x) - log(e))))^f} \quad (1)$$

From that model, the `pcrfit_single()` function estimates the variables `b_slope` and `c_intercept`, describing the slope and the y-intercept. The number of iterations required to adapt the model is also stored. That value is returned by the `pcrfit_single()` function as `convInfo_iteratons`. The higher the `convInfo_iteratons` value, the more iterations are necessary to converge from the start parameters (Figure 12L). A low `convInfo_iteratons` value is an indicator for

- a sigmoid curve shape or
- close start parameters.

High iterations numbers imply

- noisy amplification curves or
- non-sigmoid amplification curves.

The amplification curve fitting process continues with the four-parameter model (*l4*, Equation 2). This is followed by a model with five parameters (*l5*, Equation 3) and six parameters (*l6*, Equation 4).

- **l4**:

$$f(x) = c + \frac{d - c}{1 + exp(b(log(x) - log(e)))} \tag{2}$$

- **l5**:

$$f(x) = c + \frac{d - c}{(1 + exp(b(log(x) - log(e))))^f} \tag{3}$$

- **l6**:

$$f(x) = c + k \cdot x + \frac{d - c}{(1 + exp(b(log(x) - log(e))))^f} \tag{4}$$

The optimal model is selected on the basis of the Akaike information criterion and used for all further calculations. The `pcrfit_single()` function returns `qPCRmodel` as a factor (*l4*, *l5*, *l6*, *l7*). In case no model could be fitted, an *lNA* is returned.

The model is an indicator of the amplification curve shape. Model with many parameters deviate more from an ideal sigmoid model. For instance, a four-parameter model, unlike the six-parameter model, does not have a linear component. A negative linear slope in the plateau phase is an indicator of a *hook effect* (Burdukiewicz et al. 2018).

###### 4.1.6 `winklR()` - A function to calculate the central angle based on the first and the second derivative of an amplification curve data

`winklR()` is a function to calculate the in the trajectory of the first negative and the second negative derivatives maxima and minima (Figure 9) of an amplification curve data from a quantitative PCR experiment. For the determination of the angle, the origin is the maximum of the first derivative. On this basis, the vectors to the approximate minimum and maximum of the second derivatives are determined. The vectors result from the relation of the maximum of the first derivative to the minimum of the second derivative and from the maximum of the first derivative to the maximum of the second derivative. In a simple trigonometric approach, the scalar product of the two vectors is formed first. Then the absolute values are calculated and multiplied by each other. Finally, the value is converted into an angle with the cosine. The assumption is that flat (negative amplification curves) have a large angle and sigmoid (positive amplification curves) have a smaller angle. Another assumption is that this angle is independent of the rotation of the amplification curve. This means that systematic off-sets, such as those caused by incorrect background correction, are of no consequence. The cycles to be analyzed is defined by the user. The output contains the angle and the coordinates of the minima and maxima.

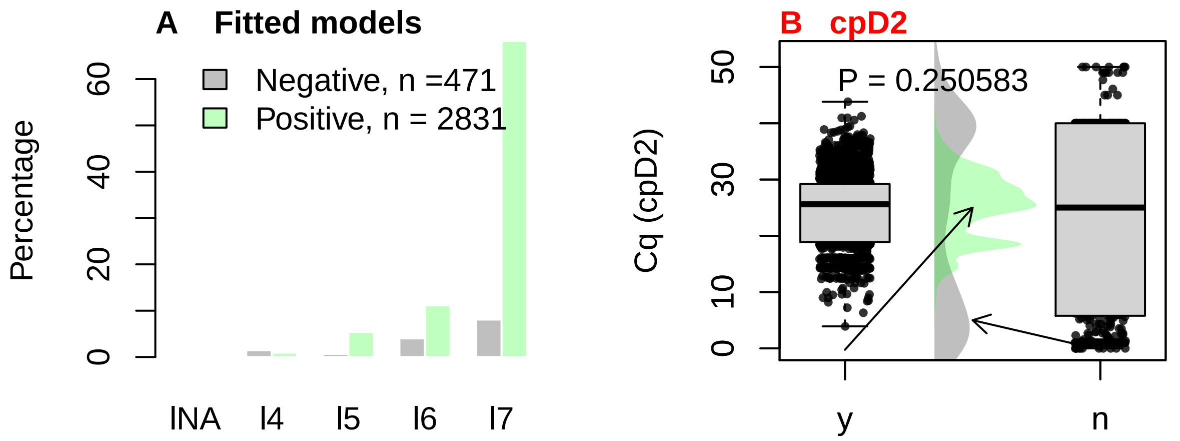

Figure 7: Frequencies of the fitted multiparametric models and Cq values. The amplification curves (n = 3302) of the 'data_sample' data set were analyzed with the encu() function. The amplification curves have been stratified according to their classes (negative: grey, positive: green). A) The optimal multiparametric model was selected for each amplification curve based on the Akaike information criterion. lNA stands for 'no model' and l4 . . . l7 for a model with four to seven parameters. B) All Cq values were calculated from optimal multiparametric models. Cqs of positive amplification curves accumulate in the range between 15 and 30 PCR cycles (50%). For the negative amplification curves, the Cqs are distributed over the entire span of the cycles. Note: The Cqs of the negative amplification curves are false-positive!

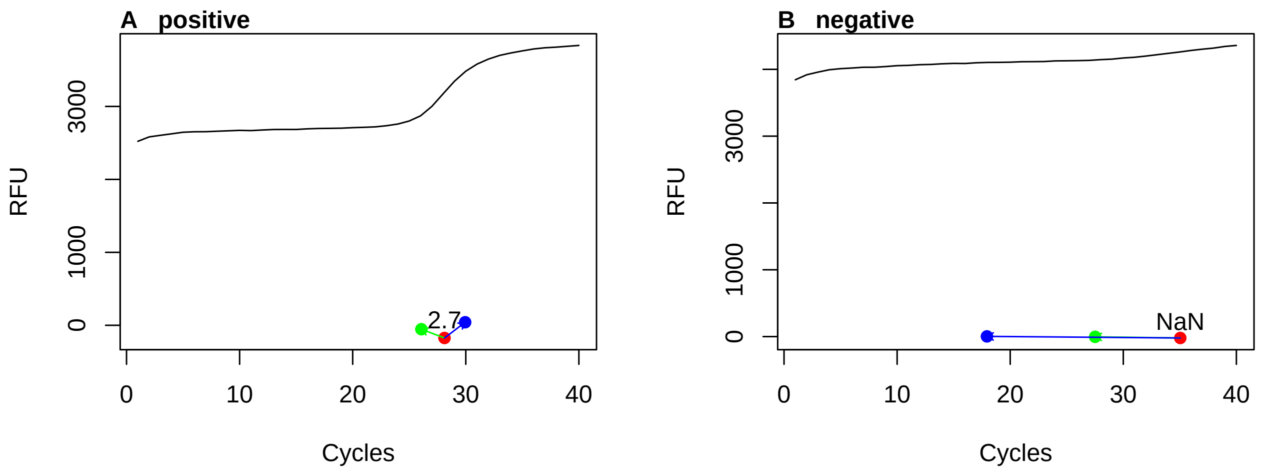

Figure 8: Concept of the winklR() function. Analysis of the amplification curves of the 'RAS002' data set with the winklR() function. Two amplification curves (A: positive, B: negative) were used. The red point shows the origin (first negative derivative maximum) while the green and blue points show the minimum and maximum of the second negative derivative. The angle is calculated from these points. Positive curves have smaller angles than negative curves.

```
## dec
##   y   n
##  42 150
```

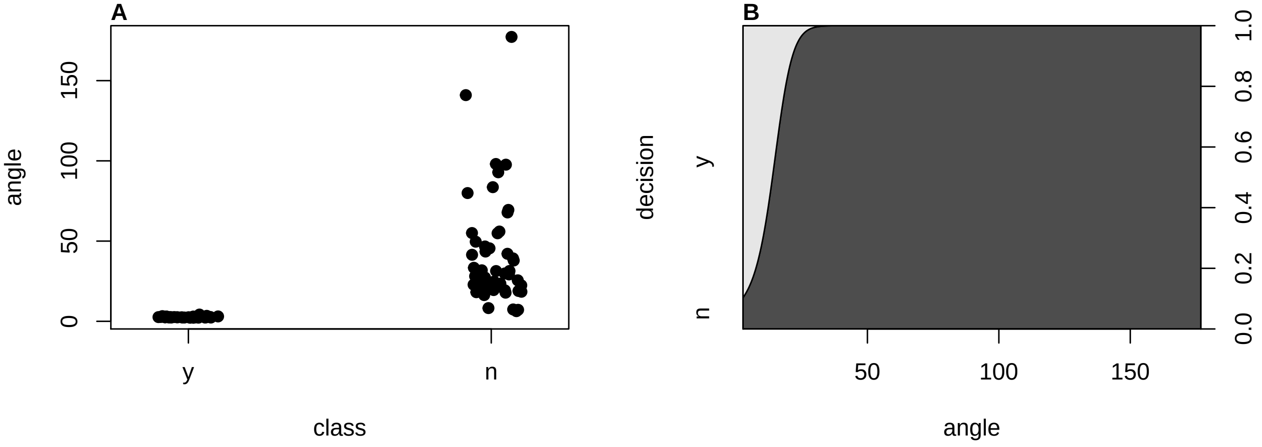

Figure 9: Analysis of the amplification curves of the ‘RAS002‘ data set with the winklR() function. All amplification curves of the data set ‘RAS002‘ were analyzed. Negative amplification curves are shown in red and positive amplification curves in black. The winklR() function was used to analyze all amplification curves. B) The stripchart of the analysis of positive and negative amplification curves shows separation. C) The cdplot calculates the conditional densities of x based on the values of y weighted by the boundary distribution of y. The densities are derived cumulatively via the values of y. The probability that the decision is negative (n) when the angle equals 30 is approximately 100%.

###### 4.1.7 Quantification Points, Ratios and Slopes

The `pcrfit_single()` function calculates `cpD1` and `cpD2` and uses them for further analysis. Both the `cpD1` and `cpD2` value are used to describe the amplification reaction quantitatively. For example, low `cpD1` and `cpD2` values (< 5 cycles) indicate that the PCR reaction was negative or that the amount of input DNA was to high Figure 7B. Since the `pcrfit_single()` function gives the parameters of all models (subsubsection 4.1.5) they are part of the feature set for completeness (Figure 10). In particular, the results of the five-parameter function and of the seven-parameter function are reported.

Further predictors from the `pcrfit_single()` function are:

- `eff` is the optimized PCR efficiency found within a sliding window (Figure 6C). A linear model of cycles versus log(Fluorescence) is fit within a sliding window (for details see `sliwin()` [`qpcR`] function). The comparison of positive and negative amplification curves in Figure 12A demonstrates that the classes are significantly different from each other. The `eff` values differ significantly between negative and positive amplification curves (Figure 12A).
- `sliwin` is the PCR efficiency by the ‘window-of-linearity’ method (Spiess, Feig, and Ritz 2008) (Figure 12B). The `sliwin` values differ significantly between negative and positive amplification curves (Figure 12B).
- `cpDdiff` is the difference between the first (`cpD1`) and the second derivative maximum `cpD2` ($cpDdiff = cpD1 - cpD2$) **from the fitted model** (Figure 6C). Provided that a model can be exactly fitted, the estimates of the difference are reliable. Higher `cpDdiff` values indicate a negative amplification reaction or a very low amplification efficiency. The comparison of positive and negative amplification curves in Figure 12C demonstrates that the classes are significantly different from each other. In the event that the `cpDdiff` value cannot be determined (NA), it is replaced by zero. The `cpDdiff` values differ significantly between negative and positive amplification curves (Figure 12C).
- `cpD2_range` is the absolute value of the difference between the minimum and the maximum of the second derivative maximum ($cpD2\_range = |cpD2m - cpD2|$) from the `diffQ2()` function (**no model fitted**) (Figure 11E). The `cpD2_range` value does not require an adjustment of a multiparametric model. The approximate first and second derivatives are determined using a five-point stencil (Rödiger,

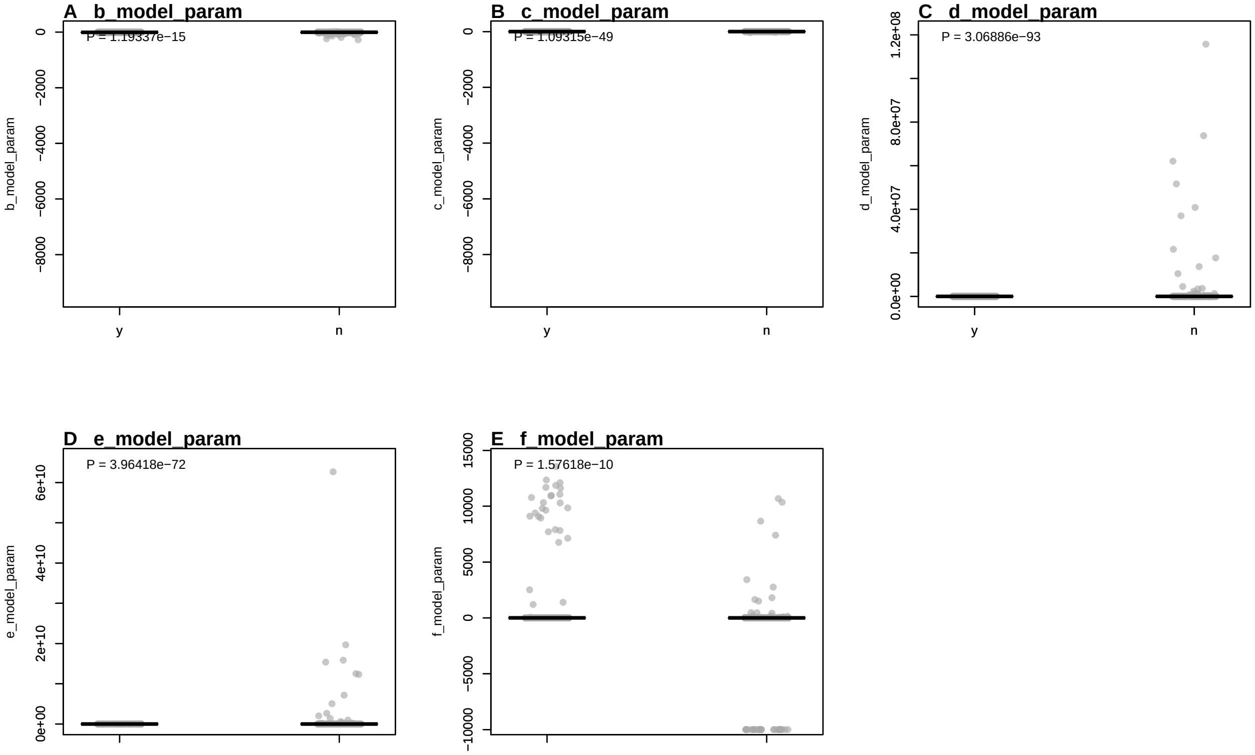

Figure 10: Values of predictors calculated from negative and positive amplification curves. Amplification curve predictors from the ‘data_sample_subset‘ data set were used as they contain positive and negative amplification curves, as well as amplification curves that exhibit a *hook effect* or non-sigmoid shapes. A) ‘c_model_param‘, is the c model parameter of the seven parameter model. B) ‘d_model_param‘, is the d model parameter of the seven parameter model. C) ‘e_model_param‘, is the e model parameter of the seven parameter model. D) ‘f_model_param‘, is the f model parameter of the seven parameter model. The classes were compared using the Wilcoxon Rank Sum Test.

Burdukiewicz, and Schierack 2015). The comparison of positive and negative amplification curves in Figure 12E shows that the classes differ significantly from each other. In the event that the `cpD2_range` value cannot be determined (NA), it is replaced by zero. The `cpD2_range` values differ significantly between negative and positive amplification curves (Figure 12E).

- `cpD2_approx` is the approximate second derivative maximum. In most cases the value should be close to the `cpD2`. Deviations indicate noise in the data, negative amplification curves or positive amplification curves that deviate from a typical sigmoid amplification curve.
- `cpD2_ratio` is is the ratio between the the approximate second derivative maximum `cpD2_approx` and the second derivative maximum `cpD2` ($cpD2_ratio = cpD2/cpD2_approx$). In the event that the `cpD2_ratio` value cannot be determined (NA, Inf), it is replaced by zero. Provided that a model can be exactly fitted and that the function has little noise the ratio between both values should be close to 1. Note: Empirical data suggest that an interval of 0.85 and 1.1 indicates positive amplification curves. These can be set to 1. Values outside this interval indicate non-sigmoidal (e. g.„ negative) amplification curves. These can be set to 0.

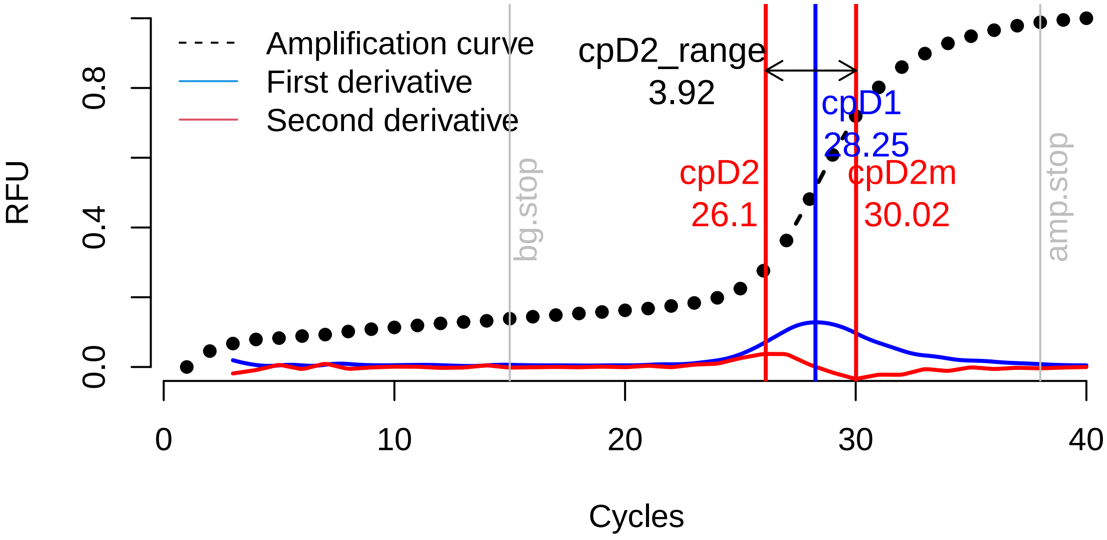

Figure 11: Location of the predictors ‘cpD2_range‘, ‘bg.start‘, ‘bg.stop‘ within an amplification curve. The minimum (cpD2m) and maximum (cpD2) of the second derivative were calculated numerically using the diffQ2() function. This function also returns the maximum of the first derivative (cpD1). The ‘cpD2_range‘ is defined as $cpD2\_range = |cpD2 - cpD2m|$. Large ‘cpD2_range‘ values indicate a low amplification efficiency or a negative amplification reaction. The predictor ‘bg.start‘ is an estimate for the end of the ground phase. ‘bg.start‘ is an approximation for the onset of the plateau phase.

- `bg.stop` is the end of the ground phase and `amp.stop` is the end of the exponential phase estimated by the `bg.max()` [chipPCR] function (Rödiger, Burdukiewicz, and Schierack 2015). A graphical presentation of the locations in the amplification curve are shown in Figure 11. The `bg.stop` and `bg.stop` values differ significantly between negative and positive amplification curves (Figure 12J & K).

- `top` is the *takeoff point* as proposed by Tichopad et al. (2003). The `top` is calculated using externally studentized residuals, which tested to be an outlier in terms of the t-distribution. The `top` signifies to first PCR cycle entering the exponential phase. `tdp` is the *takedown point*. This is an implementation in the `pcrfit_single()` function, which uses the rotated $f(x) \mapsto f_1(f(x))$ and flipped $g(x) = -(x)$ amplification curve for calculation. Figure 5A describes the location of `top` and `tdp`. The position (`f.top`, `f.tdp`) on the ordinate is also determined from these points. If an amplification curve is negative or neither `top` nor `tdp` can be calculated, then `top` & `tdp` will be assigned the number of cycles and `f.top` & `f.tdp` the value 1. The distribution of `top`, `tdp`, `f.top` and `f.tdp` is shown in Figure 12F-I. The `top`, `tdp`, `f.top` and `f.tdp` values differ significantly between negative and positive amplification curves. Potentially they enable a qualitative classification of the amplification reaction. An interesting aspect is that the positive `f.top` values are markedly lower than the negative `f.top` values The same applies inversely to the `tdp` values. In this way, amplification curves can be classified according to these values.

- `peaks_ratio` is based on a sequential chaining of functions. The `diffQ()` [`MBmca`] function determines numerically the first derivative of an amplification curve. This derivative is passed to the `mcaPeaks()` [`MBmca`] function. In the output all minima and all maxima are contained. The ranges are calculated from the minima and maxima. The Lagged Difference is determined from the ranges of the minima and maxima. Finally, the ratio of the differences (maximum/minimum) is calculated. The `peaks_ratio` values differ significantly between negative and positive amplification curves (Figure 22B).

- `loglin_slope` is calculated from the slope determined by a linear model of the data points from the cycle dependent fluorescence at the minimum of the second derivative and maximum of the second derivative (Figure 13), provided that the locations of the minimum of the second derivative and the maximum of the second derivative yield a *suitable* interval. As a precaution, the algorithm checks, for example, whether the distance between the minimum of the second derivative and the maximum of the second derivative is not more than nine PCR cycles. Failing this, the `loglin_slope` value is set to zero (no slope), as in the example of Figure 13. The `loglin_slope` values differ significantly between negative and positive amplification curves (Figure 12D).

The predictor `loglin_slope` is used in the following to test if the slope within this ROI can be used to distinguish positive and negative amplification curves. The hypothesis is that positive amplification curves have a higher `loglin_slope` than negative amplification curves. As shown in Figure 12D, there is a statistically significant difference between positive and negative amplification curves.

The `loglin_slope` values from the `data_sample_subset_balanced` data set (subsubsection 4.1.2) were used to save computing time. A binomial logistic regression (see subsection 2.4)) was used to analyze the relationship between the `loglin_slope` value and the class (negative, positive). The data set was split into two chunks. This is an important step during such applications. One chunk is for adapting, i. e. training, the model and the other chunk for testing the model. By convention, 70% to 80% of the data is used for training (Walsh, Pollastri, and Tosatto 2015; Kuhn 2008). The binomial logistic regression model was adapted using the `glm()` [`stats`] function by using the parameter `family = binomial(link = 'logit')`. To objectify the splitting, the `sample()` [`stats`] function was used.

```
library(PCRedux)

data <- data_sample_subset_balanced

n_positive <- sum(data[["decision"]] == "y")
n_negative <- sum(data[["decision"]] == "n")

dat <- data.frame(loglin_slope = data[, "loglin_slope"],
                  decision = as.numeric(factor(data$decision,
                                               levels = c("n", "y"),
                                               label = c(0, 1))) - 1)

# Select randomly observations from 70% of the data for training.
# n_train is the number of observations used for training.

n_train <- round(nrow(data) * 0.7)

paste0("Percentage of observations (", n_train, ") = ",
       signif((n_train/nrow(data)*100),3), "%")
```

```
## [1] "Percentage of observations (456) = 70%"
```

```
# index_test is the index of observations to be selected for the training
index_test <- sample(1L:nrow(dat), size = n_train)
```

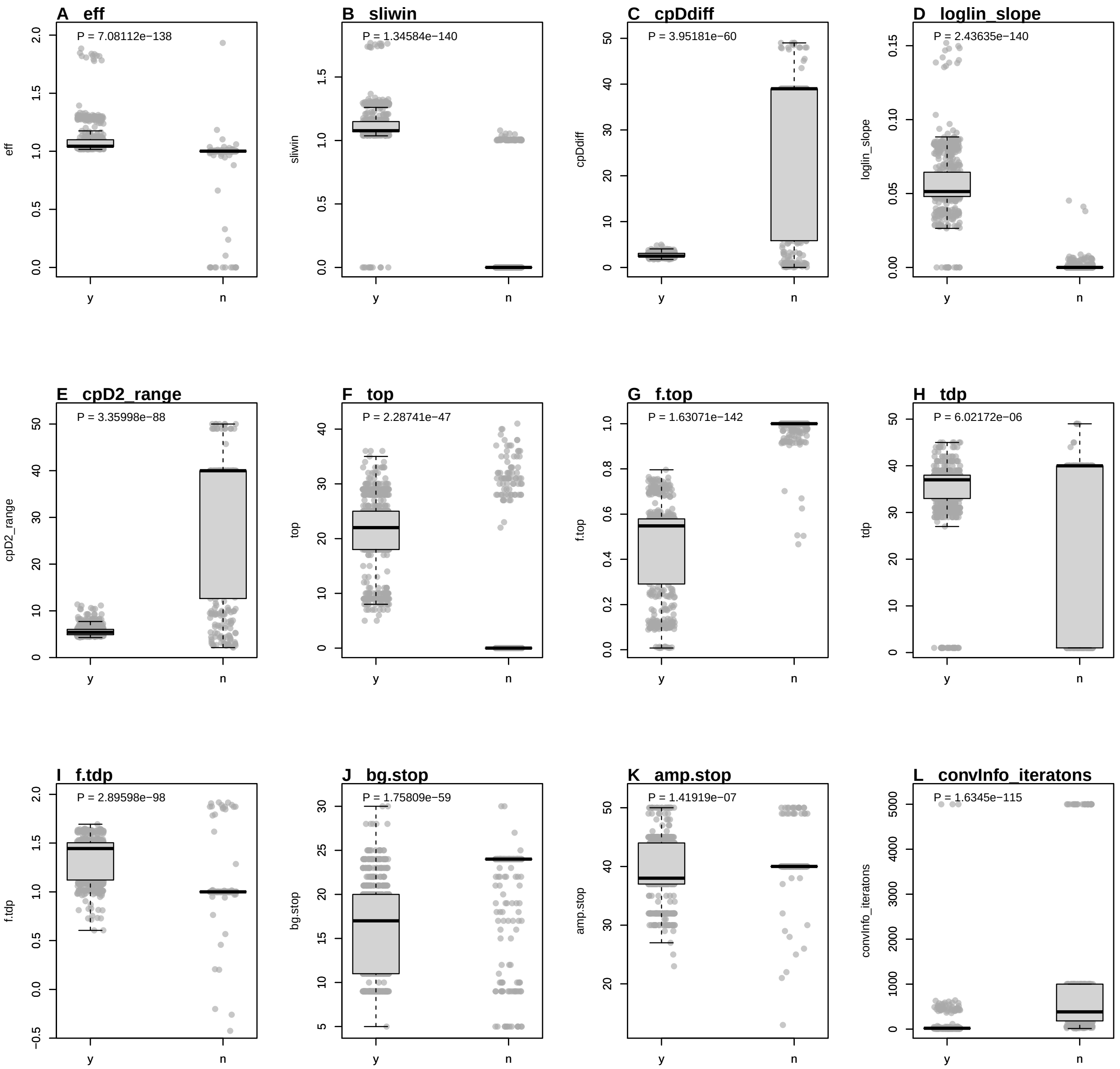

Figure 12: Values of predictors calculated from negative and positive amplification curves. Amplification curve predictors from the ‘data_sample_subset‘ data set were used as they contain positive and negative amplification curves, and amplification curves that exhibit a *hook effect* or non-sigmoid shapes. A) ‘eff‘, optimized PCR efficiency found within a sliding window. B) ‘sliwin‘, PCR efficiency by the ‘window-of-linearity‘ method. C) ‘cpDdiff‘, difference between the Cq values calculated from the first and the second derivative maximum. D) ‘loglin_slope‘, slope from the cycle at the second derivative maximum to the second derivative minimum. E) ‘cpD2_range‘, absolute value of the difference between the minimum and the maximum of the second derivative maximum. F) ‘top‘, takeoff point. G) ‘f.top‘, fluorescence intensity at takeoff point. H) ‘tdp‘, takedown point. I) ‘f.tdp‘, fluorescence intensity at takedown point. J) ‘bg.stop‘, estimated end of the ground phase. K) ‘amp.stop‘, estimated end of the exponential phase. L) ‘convInfo_iteratons‘, number of iterations until convergence.

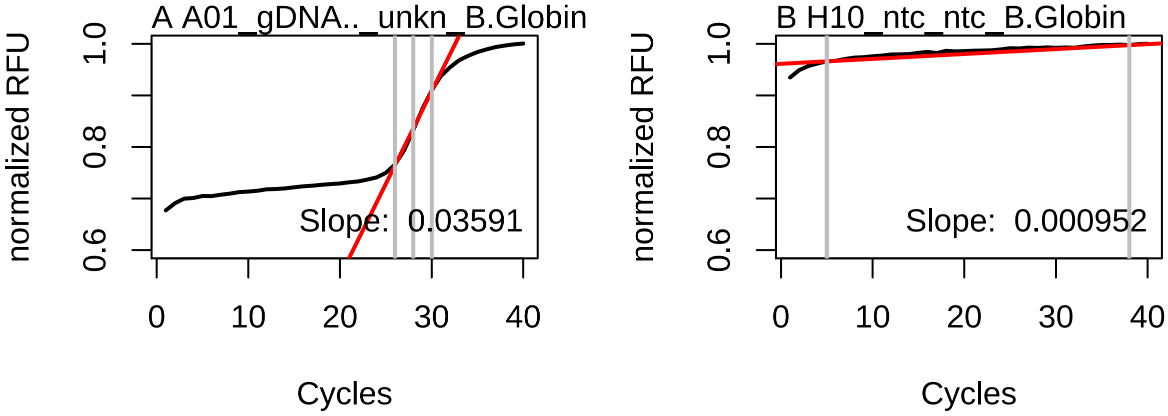

Figure 13: Concept of the 'loglin_slope' predictor. The algorithm determines the fluorescence values of the raw data at the approximate positions of the maximum off the first derivative, the minimum of the second derivative and the maximum of the second derivative, which are in the exponential phase of the amplification curve. The data were taken from the 'RAS002' data set. A linear model is created from these parameter sets and the slope is determined. A) Positive amplification curves have a clearly positive slope. B) Negative amplification curves usually have a low, sometimes negative slope.

```
# index_test is the index of observations to be selected for the testing
index_training <- which(!(1L:nrow(dat) %in% index_test))

# train_data contains the data used for training

train_data <- dat[index_test, ]

# test_data contains the data used for training

test_data <- dat[index_training, ]

# Fit the binomial logistic regression model

model_glm <- glm(decision ~ loglin_slope, family=binomial(link='logit'),
                 data = train_data)

predictions <- ifelse(predict(model_glm,
                              newdata = test_data, type = 'response') > 0.5,
                      1, 0)

res_performeR <- performeR(predictions, test_data[["decision"]])[, c(1:10, 12)]
```

The `summary()` function returns the results of the model fitting. This can be analysed and interpreted.

```
summary(model_glm)
```

```
##
## Call:
## glm(formula = decision ~ loglin_slope, family = binomial(link = "logit"),
##     data = train_data)
##
## Deviance Residuals:
##      Min        1Q    Median        3Q       Max
```

```
## -3.05451  -0.27685  -0.27685   0.01242   2.56156
##
## Coefficients:
##              Estimate Std. Error z value Pr(>|z|)
## (Intercept)   -3.2425     0.3296  -9.836   <2e-16 ***
## loglin_slope 174.6584    18.9726   9.206   <2e-16 ***
## ---
## Signif. codes:  0 '***' 0.001 '**' 0.01 '*' 0.05 '.' 0.1 ' ' 1
##
## (Dispersion parameter for binomial family taken to be 1)
##
##     Null deviance: 632.07  on 455  degrees of freedom
## Residual deviance: 112.82  on 454  degrees of freedom
## AIC: 116.82
##
## Number of Fisher Scoring iterations: 8
```

Based on the results it can be concluded that the parameters `(intercept)` and `loglin_slope` are statistically significant ($P < 2\text{e-}16$). This indicates an association between `loglin_slope` and the probability that an amplification curve is positive.

In order to apply the model to a new data set, further steps are necessary. `predict()` [`stats`] is a generic function for predicting the results of a model fitting function (Figure 14A). All previously split test data is passed to the function argument `newdata`. By setting the `type = 'response'` parameter, the `predict()` function returns probabilities in the form of $P(y = 1|X)$. In the case in hand, it was decided that a decision limit of 0.5 is to be applied. If $P(y = 1|X) < 0.5$ then $y = 0$ (amplification curve negative), otherwise $y = 1$ (amplification curves positive).

```r
library(PCRedux)

# Create graphic device for the plot(s)
# Plot train_data (grey points) and the predicted model (blue)
par(mfrow = c(1,2))

plot(train_data$loglin_slope, train_data$decision, pch = 19,
     xlab = "loglin_slope", ylab = "Probability",
     col = adjustcolor("grey", alpha.f = 0.9), cex = 1.5)
mtext("A", cex = 1, side = 3, adj = 0, font = 2, las = 0)
abline(h = 0.5, col = "grey")

curve(predict(model_glm, data.frame(loglin_slope = x), type = "resp"),
      add = TRUE, col = "blue")

# Plot test_data (red)

points(test_data$loglin_slope, test_data$decision, pch = 19,
       col = adjustcolor("red", alpha.f = 0.3))
legend("right", paste("Positive: ", n_positive,
                      "\nNegative: ", n_negative), bty = "n")

# Plot the sensitivity, specificity and other measures to describe
# the prediction.
```

```r
position_bp <- barplot(as.matrix(res_performeR), yaxt = "n",
                       ylab = "Probability", main = "", las = 2,
                       col = adjustcolor("grey", alpha.f = 0.5),
                       border = "white")

par(srt = 90)
text(position_bp, rep(0.8, length(res_performeR)),
     paste(signif(res_performeR, 2)*100, "%"), cex = 0.6)
axis(2, at = c(0, 1), labels = c("0", "1"), las = 2)
abline(h = 0.85, col = "grey")

mtext("B", cex = 1, side = 3, adj = 0, font = 2, las = 0)
```

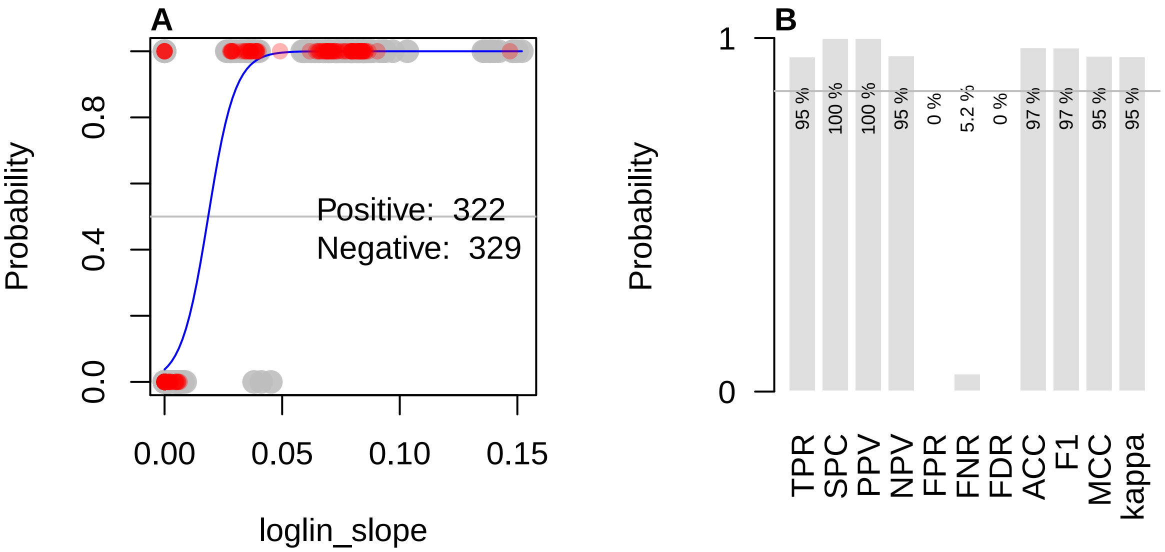

Figure 14: Machine classification by means of binomial logistic regression using the ‘loglin_slope‘ predictor. A) For the calculation of a binomial logistic regression model, the categorical response variable $Y$ (decision with classes: negative and positive) must be converted to a numerical value. With binomial logistic regression, the probability of a categorical response can be estimated using the $X$ predictor variable. In this example, the predictor variable ‘loglin_slope‘ is used. Grey measurement points (70% of the data set) were used for training. Red dots represent the values used for testing. The regression curve of the binomial logistic regression is shown in blue. The grey horizontal line at 0.5 marks the threshold of probability above which it is determined whether an amplification curve is negative or positive. B) The performance indicators were calculated using the performeR() function. Sensitivity, TPR; Specificity, SPC; Precision, PPV; Negative prediction value, NPV; Fall-out, FPR; False negative rate, FNR; False detection rate, FDR; Accuracy, ACC; F1 score, F1; Matthews correlation coefficient, MCC, Cohens kappa (binary classification), kappa ($\kappa$).

Sensitivity, specificity and further parameters for estimating predictions were calculated using the `performeR()` function (subsubsection 4.1.1). The results indicate that the sensitivity and specificity for the test data set provides a good result. However, in this case, they depend heavily on the computer-aided random sampling of the training data, and the total size of the data set (Figure 14B). Over-fitting and under-fitting and other problems need to be addressed (Walsh, Pollastri, and Tosatto 2015).

To proof the results, further methods such as Likelihood Ratio Test, McFadden’s $R^2$, k-fold cross-validation, Receiver Operating Characteristic (ROC) analysis and model interpretation should be used (Arlot and Celisse 2010; McFadden 1974; Sing et al. 2005).

- `sd_bg` is the standard deviation from the first PCR cycle to the takeoff point (Figure 5A). Manufacturers of thermo-cyclers use different sensors and data processing algorithms. The same applies to the detection chemistry used in experiments section 3. The signal variation in the ground phase differs between the different systems (Figure 3D). If no takeoff point can be determined from an amplification curve, the value for `sd_bg` is calculated from the first to the eighth PCR cycle. The results for the predictor `sd_bg` were broken down by the thermo-cycler and the output of the amplification reaction (negative, positive). It can be seen that the signal variation between the thermo-cyclers seems to be different. There is also a difference between negative and positive amplification curves Figure 15. The `sd_bg` values differ significantly between negative and positive amplification curves (Figure 16J).

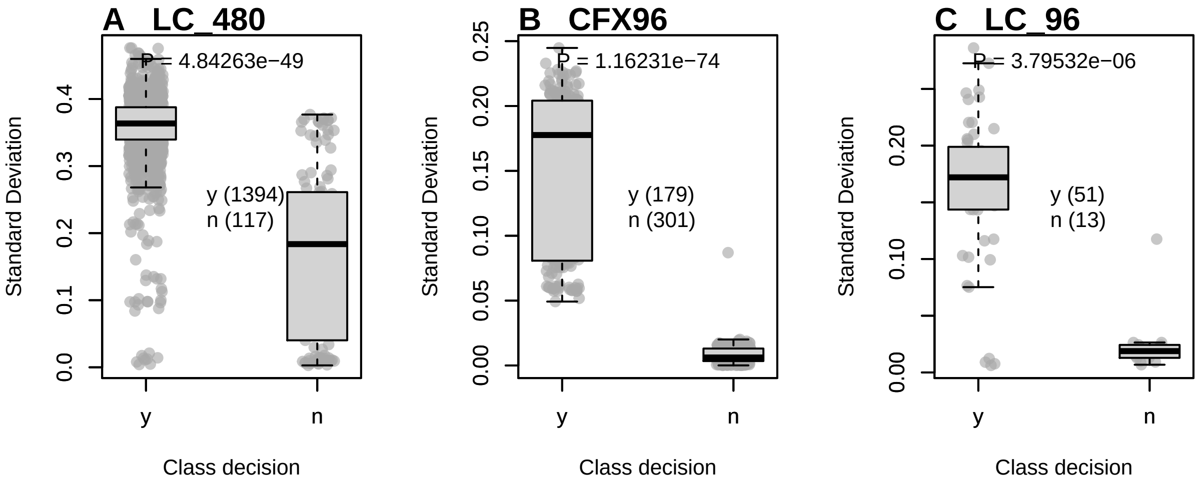

Figure 15: Standard deviation in the ground phase of various qPCR devices. The ‘sd_bg‘ predictor was used to determine if the standard deviation between thermo-cyclers and between positive and negative amplification curves was different. The standard deviation was determined from the fluorescence values from the first cycle to the takeoff point. If the takeoff point could not be determined, the standard deviation from the first cycle to the eighth cycle was calculated. The Mann-Whitney test was used to compare the medians of the two populations (y, positive; n, negative). The differences were significant for A) LC_480 (Roche), B) CFX96 (Bio-Rad) and C) LC96 (Roche).

###### 4.1.8 Integration of the `amptester()` function in `PCRedux`

`amptester_polygon` is another method to calculate the area under an amplification curve. `amptester_polygon`[5] is part of the `amptester()` [`chipPCR`] package (Rödiger, Burdukiewicz, and Schierack 2015). In contrast to the implementation in `amptester()`, `amptester_polygon` has values normalized to the total number of cycles, thereby allowing comparable predictions (Figure 22Ds).

###### 4.1.9 `earlyreg()` - A Function to Calculate the Slope and Intercept in the Ground Phase of an Amplification Curve

The signal height and the slope in the first cycles (1 - 10) of amplification curves are potentially useful because some qPCR systems calibrate themselves by fluorescence intensity of the first cycles. This is noticeable as strong signal changes which appear spontaneously between the first and second cycle (e. g., Figure 1B). Furthermore, the signal level can be used to determine which background signal is present and whether the ground phase already has a slope. Moreover, characteristics of the detection probe system are noticeable (see section 4). From the slope, it may be deduced whether amplification has already started (see section 4).

Consequently, the `earlyreg()` function was developed. This function uses an ordinary least squares linear regression within a limited number of cycles. As ROI, the first 10 cycles were defined. This restriction is based on the developers experience, suggesting that during the first ten cycles only a significant increase in signal strength can be measured within few qPCRs. However, `earlyreg()` does not ignore the first cycle, as many thermo-cyclers use this cycle for sensor calibration. Extreme values are therefore included. As standard,

[5] This predictor is determined from the points in an amplification curve (like a polygon, in particular non-convex polygons) in a ‘clockwise‘ order. The sum over the edges result in a positive value if the amplification curve is ‘clockwise‘ and is negative if the curve is ‘counter-clockwise‘.

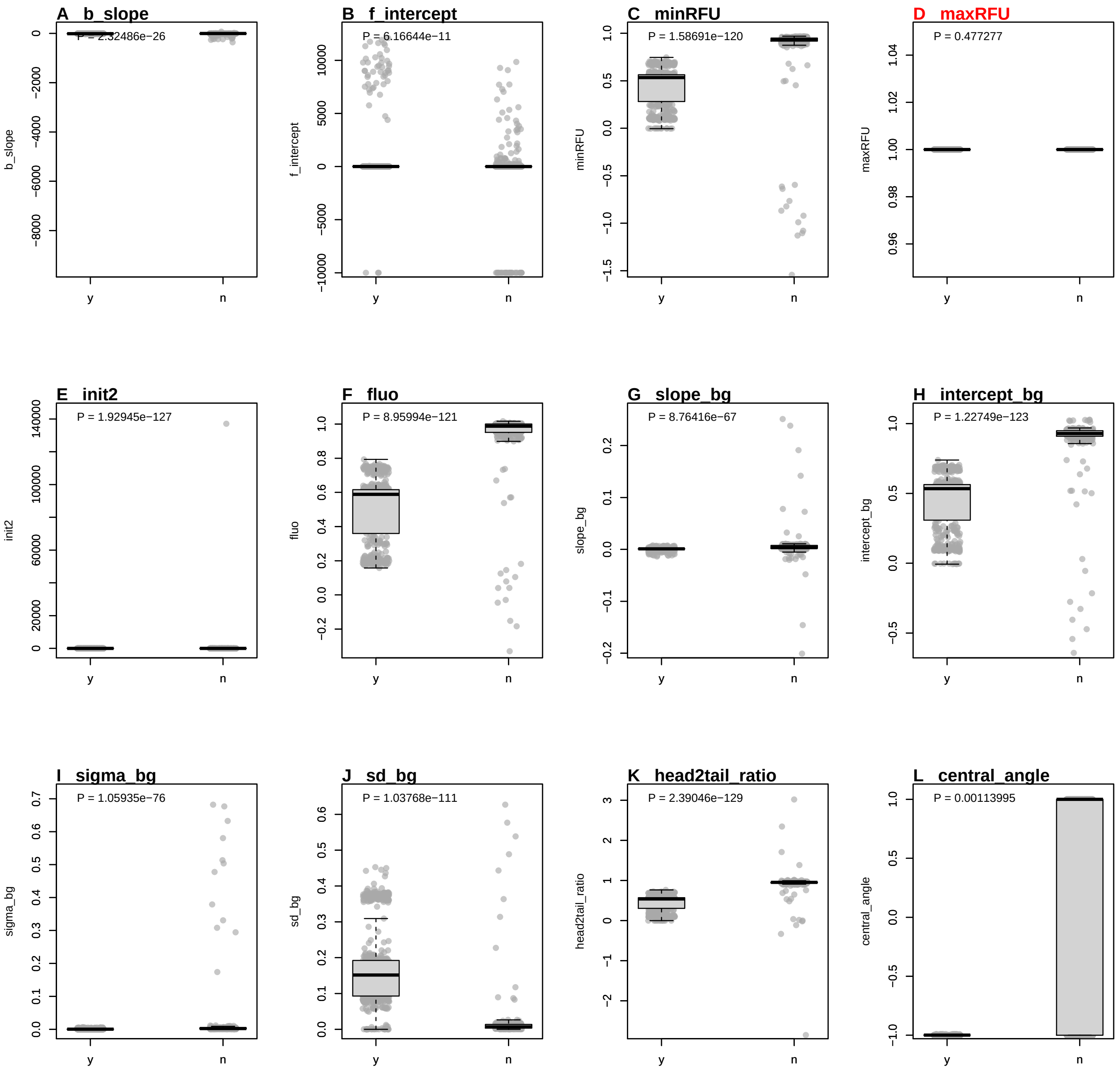

Figure 16: Values of predictors calculated from negative and positive amplification curves. Amplification curve predictors from the ‘data_sample_subset‘ data set were used as they contain positive and negative amplification curves, as well as amplification curves that exhibit a *hook effect* or non-sigmoid shapes. A) ‘eff‘, optimized PCR efficiency in a sliding window. B) ‘sliwin‘, PCR efficiency according to the window-of-linearity method. C) ‘cpDdiff‘, difference between the Cq values calculated from the first and the second derivative maximum. D) ‘loglin_slope‘, slope from cycle at second derivative maximum to second derivative minimum. E) ‘cpD2_range‘, absolute difference between the minimum and maximum of the second derivative. F) ‘top‘, takeoff point. G) ‘f.top‘, fluorescence intensity at takeoff point. H) ‘tdp‘, takedown point. I) ‘f.tdp‘, fluorescence intensity at the takedown point. J) ‘bg.stop‘, estimated end of the ground phase. K) ‘amp.stop‘, estimated end of the exponential phase. L) ‘convInfo_iteratons‘, number of iterations until convergence when fitting a multiparametric model. The classes were compared using the Wilcoxon Rank Sum Test.

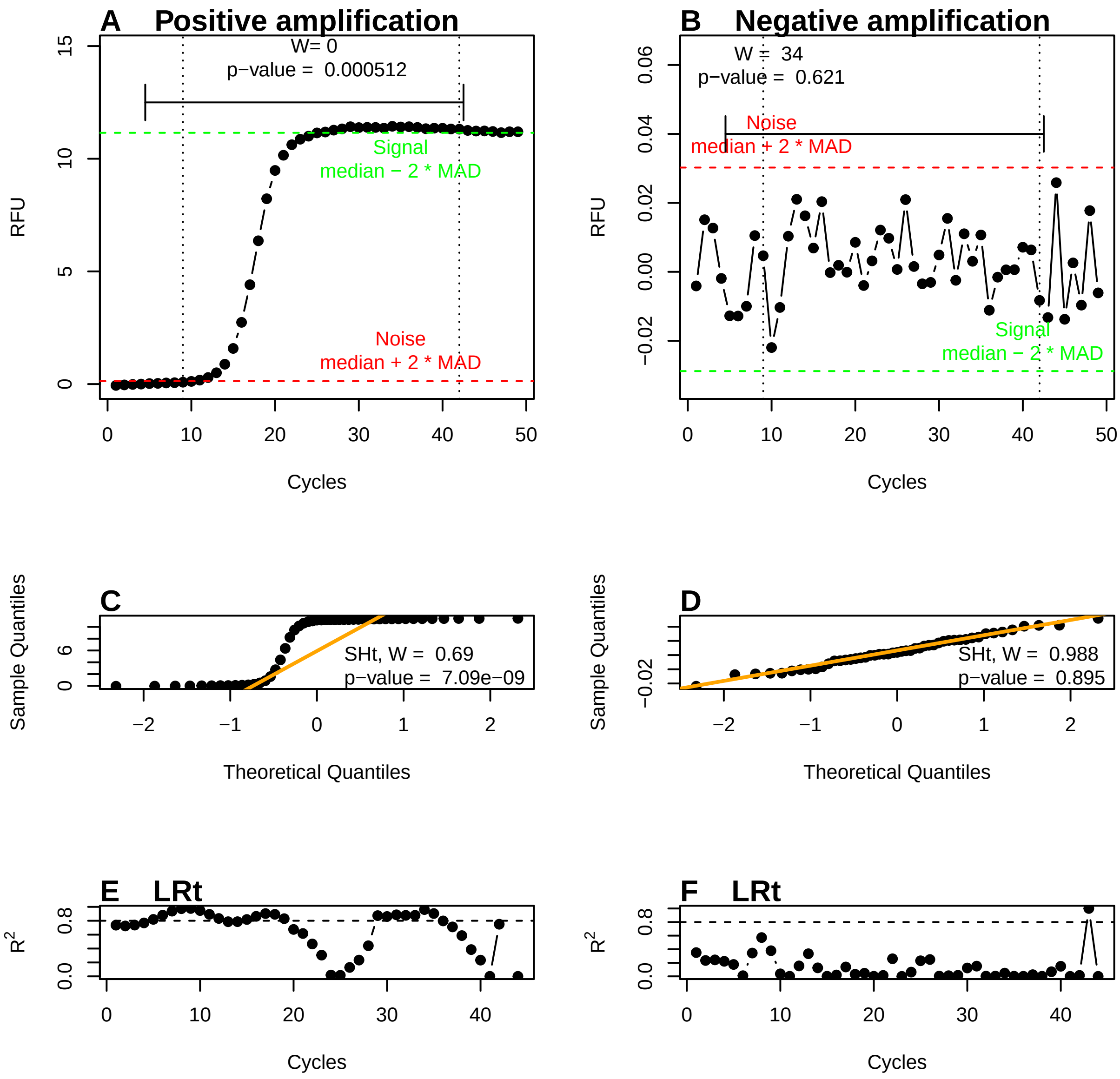

Figure 17: Analysis of amplification curves with the "amptester()" function. A & B) The threshold test (THt) is based on the Wilcoxon ranksum test and compares 20% of the fluorescence values of the ground phase with 15% of the plateau phase. In the example, a significant difference ($p = 0.000512$) was found for the positive amplification curve. However, this did not apply to the negative amplification curve ($p = 0.621$). C & D) A Q-Q diagram is used to graphically compare two probability distributions. In this study the probability distribution of the amplification curve was compared with a theoretical normal distribution. The orange line is the theoretically normal quantil-quantile plot that passes through the probabilities of the first and third quartiles. The Shapiro-Wilk test (SHt) of normality checks whether the underlying measurement data of the amplification curve is significantly normal distributed. Since the p-value of $7.09e^{-9}$ of the positive amplification curve is $\alpha \leq 5e^{-4}$, the null hypothesis is rejected. However, this does not apply to the negative amplification curve ($p = 0.895$). E & F) The linear regression test (LRt) calculates the coefficient of determination ($R^2$) using an ordinary least square regression where all measured values are integrated into the model in a cycle-dependent manner. Experience shows that the non-linear part of an amplification curve has a $R^2$ smaller than 0.8, which is also shown in the example.

the next nine amplitude values are used for the linear regression. The number of cycles can also be adjusted via the parameter `range`. Since all amplification curves are normalized to the 99%-percentile, comparability between the background signals and the slopes is ensured. The output of the `earlyreg()` function is:

- `slope_bg`, which is the slope of the ordinary least squares linear regression model,
- `intercept_bg`, which is the intercept of the linear model and
- `sigma_bg`, which is the square root of the estimated variance of the random error.

The `slope_bg`, `intercept_bg` and `sigma_bg` values differ significantly between negative and positive amplification curves (Figure 16G-I).

The following example illustrates possible usage of `earlyreg()`. For that purpose, amplification curves from the `C127EGHP` data set were analysed (Figure 18A). In figure Figure 18A the amplification curves for all cycles are shown. Next, the `earlyreg()` function was used to determine the `slope_bg`, `intercept_bg` and `sigma_bg` in the range of the first ten PCR cycles. The results were used in a cluster analysis using k-means clustering, demonstrating that the slope seems to be an indicator of differences between the amplification curves. The first 8 cycles were colored according to their cluster (Figure 18B). After cluster analysis, the same could also be observed (Figure 18D-F). Hence, it can be postulated that the slope in the background phase is useful for the amplification curve classification.

```
##      intercept      slope      sigma
## EG1 -0.9996982 -1.0000257 -0.9994170
## EG2 -0.9999187 -0.9999724 -0.9994349
## EG3 -1.0001748 -0.9999236 -0.9995357
## EG4 -1.0000410 -0.9999487 -0.9994961
## EG5 -1.0001467 -0.9999259 -0.9995627
## EG6 -1.0003437 -0.9998825 -0.9995230
```

###### 4.1.10 `head2tailratio()` - A Function to Calculate the Ratio of the Head and the Tail of a Quantitative PCR Amplification Curve

The ratios from the ground and plateau phase can be used to search for patterns in amplification curves. Positive amplification curves have different slopes and intercepts at the start (head, background region) and the end (tail, plateau region) of the amplification curve. Hence, these regions are potentially useful to extract a predictor for amplification curve classification. Negative amplification curves (no slope) are assumed to have a ratio of about 1. In contrast, positive amplification curves should have a ratio of less than 1.

The $n$-dimensional space of all predictor variables $X_{1,2 \ldots n}$ is also called feature space. In the present study the feature space was extended by domain knowledge using known features. The `head2tailratio()`-function is an example for this. Here the feature $X_3 \equiv$ *head2tail_ratio* could be calculated by determining the ratio of the $X_1 \equiv$ *fluorescence intensity in the head region* and $X_2 \equiv$ *fluorescence intensity in the tail region* of a quantitative PCR amplification curve ($X_3 = \frac{X_1}{X_2}$). As ROI, the areas in the ground phase (head) and plateau phases (tail) are used (Figure 5A). For the calculation, the median from the first six data points of the amplification curve and the median from the last six data points are used. The determination of six data points in both regions was made on the basis of *empirical experience*. As a rule, no significant increase in amplification signals can be measured in the first six cycles and in the last six cycles (where the amplification curve usually transitions into the plateau). This assumption is sometimes violated (e. g., *hook effect*) and might lead to false estimates.

```r
library(PCRedux)

# Load the RAS002 amplification curve data set and assign it to the object data

data <- RAS002

# Load the RAS002 decision data set and assign it to the object data_decisions
data_decisions <- RAS002_decisions
```

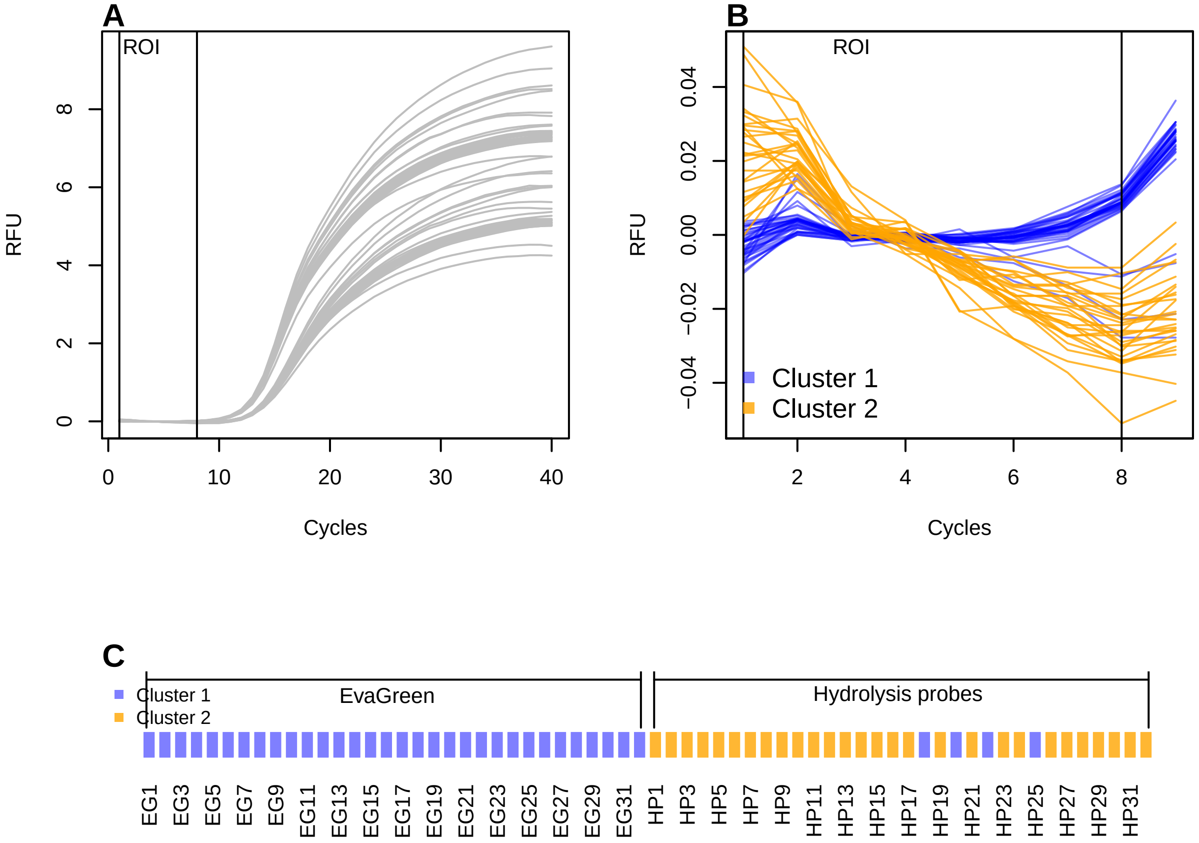

Figure 18: Analysis of the ground phase with the earlyreg() function and the 'C127EGHP' data set (n = 64 amplification curves). This data set consists of 32 samples, which were simultaneously monitored with the intercalator EvaGreen or hydrolysis probes. A) All amplification curves possess slightly different slopes and intercepts in the first cycles of the ground phase (ROI: Cycles 1 to 8). Both the slope and the intercept of each amplification curve were used for cluster analysis (k-means, Hartigan-Wong algorithm, number of centers $k = 2$). B) The amplification curves were assigned to five clusters, depending on their slope and their intersection (red, black). C) Finally, the clusters were associated to the detection chemistries (EvaGreen (EG) or hydrolysis probes (HP)).

```r
# Calculate the head2tailratio of all amplification curves

res_head2tailratio <- lapply(2L:ncol(data), function(i) {
  head2tailratio(
    y = data[, i], normalize = TRUE, slope_normalizer = TRUE,
    verbose = TRUE
  )
})

# Fetch all values of the head2tailratio analysis for a later comparison
# by a boxplot.

res <- sapply(1L:length(res_head2tailratio), function(i)
  res_head2tailratio[[i]]$head_tail_ratio)

data_normalized <- cbind(
  data[, 1],
  sapply(2L:ncol(data), function(i) {
    data[, i] / quantile(data[, i], 0.99)
  })
)

# Assign colors to the classes (n: black, y: green).
colors <- as.character(factor(
  data_decisions, levels = c("y", "n"),
  labels = c(
      adjustcolor("green", alpha.f = 0.5), adjustcolor("black", alpha.f = 0.5))
))

res_wilcox.test <- stats::wilcox.test(res ~ data_decisions)
```

The amplification curves in Figure 19 show a signal increase within the first three cycles, and those in Figure 5C have a negative slope in the tail. The median is used to minimize the influence of outliers.

```r
# Plot the results of the analysis
#
# Position and plot parameters
h <- max(na.omit(res))
h_text <- rep(h * 0.976, 2)

# Create graphic device for the plot(s)
layout(matrix(c(1,1,2), 1, 3, byrow = TRUE))

matplot(
  data_normalized[, 1], data_normalized[, -1],
  xlab = "Cycles", ylab = "normalized RFU", main = "",
  type = "l", lty = 1, lwd = 2, col = colors
)
for (i in 1L:(ncol(data_normalized) - 1)) {
  points(
    res_head2tailratio[[i]]$x_roi, res_head2tailratio[[i]]$y_roi,
    col = colors[i], pch = 19, cex = 1.5
  )
  abline(res_head2tailratio[[i]]$fit, col = colors[i], lwd = 2)
```

```
}
mtext("A", cex = 1, side = 3, adj = 0, font = 2)

# Boxplot of the head2tail ratios of the positive and negative
# amplification curves.

boxplot(res ~ data_decisions, col = unique(colors), ylab = "Head to Tail Ratio")

lines(c(1, 2), rep(h * 0.945, 2))
text(1.5, h_text, paste0("P = ", signif(res_wilcox.test[["p.value"]])),
     cex = 1)

mtext("B", cex = 1, side = 3, adj = 0, font = 2)
```

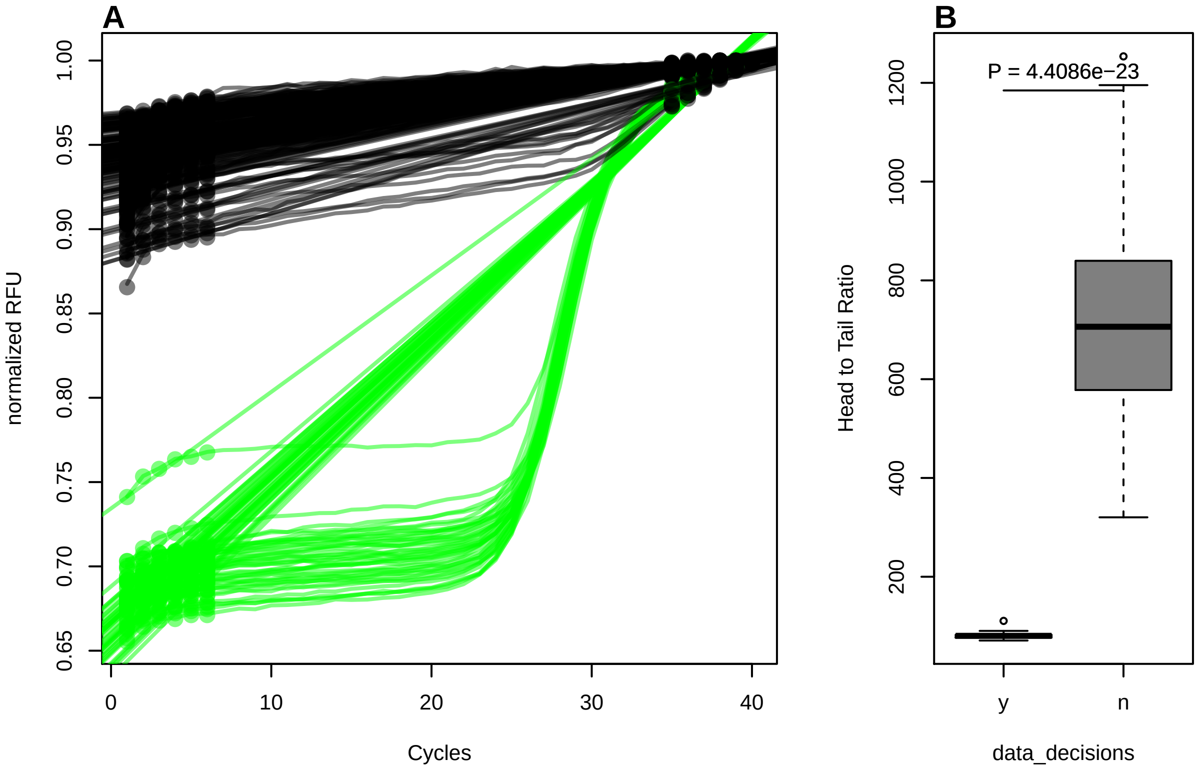

Figure 19: Ratio between the head and the tail of a quantitative PCR amplification curve. A) Plot of quantile normalized amplification curves from the 'RAS002' data set. Data points used in the head and and tail are highlighted by circles. The intervals for the Robust Linear Regression are automatically selected using the 25% and 75% quantiles. Therefore, not all data points are used in the regression model. The straight line is the regression line from the robust linear model. The slopes of the positive and negative amplification curves differ. B) Boxplot for the comparison of the *head/tail* ratio. Positive amplification curves have a lower ratio than negative curves. The difference between the classes is significant.

In section 4 and in Figure 1, it was shown that negative amplification curves may have a slope with a positive or negative sign. There is no consent in the literature and among peers how to deal with this during the processing. One solution is to include the slope as factor in the ratio calculation. The `head2tailratio()` function uses a linear model that calculates the slope between the ground and plateau phases. If the slope of the model is significant, then the ratio from the head and tail is normalized to this slope. This requires setting the `slope_normalizer` parameter in the `head2tailratio()` function. By default, this parameter is not set.

The `head2tail_ratio` values differ significantly between negative and positive amplification curves (Figure 16K).

###### 4.1.11 hookreg() and hookregNL() - Functions to Detect Hook Effect-like Curvatures

hookreg() and hookregNL() are functions to detect amplification curves bearing a *hook effect* (Barratt and Mackay 2002) or negative slope at the end of the amplification curve. Both functions calculate the slope and intercept of an amplification curve data. The assumption is that a strong negative slope at the end of an amplification curve is indicative for a *hook effect*. hookreg() and hookregNL() are part of a peer-reviewed publication (Burdukiewicz et al. 2018). For this reason, the functions will not be discussed here.

###### 4.1.12 mblrr() - A Function to Perform the Quantile-filter Based Local Robust Regression

mblrr() is a function to perform the **m**edian **b**ased **l**ocal **r**obust **r**egression (mblrr) from a quantitative PCR experiment. In detail, this function attempts to break the amplification curve in two ROIs (head (~background) and tail (~plateau)). As opposed to the earlyreg() function, the mblrr() function does not use a fixed interval. Instead, the mblrr() function dynamically determines cut points for each amplification curve. It was defined that:

- the 25% quantile is the value for which 25% of all values are smaller than this value.
- the 75% quantile is the value for which 75% of all values are greater than this value.

Subsequent, a robust linear regression analysis (lmrob()) is preformed individually on both regions of the amplification curve. The rationale behind this analysis is that the slope and intercept of an amplification curve differ in the background and plateau region. This is also shown by the simulations in Figure 3A-C. In the example shown below, the observations "P01.W19", "P06.W35", "P33.W66", "P65.W90", "P71.W23" and "P87.W01" were arbitrarily selected for demonstration purposes Figure 20. Those amplification curves have a slight negative trend in the base-line region and a positive trend in the plateau region.

The correlation coefficient[6] is a measure to quantify the dependence on variables (e. g., number of cycles, signal height). The correlation coefficient is always between -1 and 1, with a value close to -1 describing a strong-negative dependency and close to 1 describing a strong-positive dependency; if the value is 0, there is no dependency between the variables. The most frequently used correlation coefficient to describe a linear dependency is the Pearson correlation coefficient $r$. The correlation coefficient can be used as a predictor. Similar data structures have similar correlation coefficients. However, variables that are not strongly correlated can also be important for modeling.

```
library(PCRedux)

# Select four amplification curves from the RAS002 data set
amplification_curves <- c(2, 3, 4, 5, 44, 45)
data <- RAS002[, c(1, amplification_curves)]

# Load the decision_res_htPCR.csv data set from a csv file.
filename <- system.file("decision_res_RAS002.csv", package = "PCRedux")
decision_res <- read.csv(filename)

# Overview of the amplicon curve classifications
res_decision <- decision_res[amplification_curves - 1, -c(3,4)]

res_decision
```

```
##                      RAS002 test.result.1 conformity
## 1  A01_gDNA.._unkn_B.Globin             y       TRUE
## 2     A01_gDNA.._unkn_HPRT1             n       TRUE
## 3  A02_gDNA.._unkn_B.Globin             y       TRUE
```

[6] Product moment correlation coefficient (Pearson)

```
## 4     A02_gDNA.._unkn_HPRT1              n       TRUE
## 43 B10_gDNA.._unkn_B.Globin              n       TRUE
## 44    B10_gDNA.._unkn_HPRT1              n       TRUE
```

```r
# Plot the regions and the linear regression line in the
# amplification curve plot

colors <- c(
            adjustcolor("blue", alpha.f = 0.5),
            adjustcolor("orange", alpha.f = 0.8)
           )

# Create graphic device for the plot(s)
par(mfrow = c(3, 2))

for (i in 2L:ncol(data)) {
  x <- data[, 1]
  y_tmp <- data[, i] / quantile(data[, i], 0.99)
  res_q25 <- y_tmp < quantile(y_tmp, 0.25)
  res_q75 <- y_tmp > quantile(y_tmp, 0.75)
  res_q25_lm <- try(
    suppressWarnings(lmrob(y_tmp[res_q25] ~ x[res_q25])),
    silent = TRUE
  )
  res_q75_lm <- try(
    suppressWarnings(lmrob(y_tmp[res_q75] ~ x[res_q75])),
    silent = TRUE
  )

  plot(x, y_tmp, xlab = "Cycles", ylab = "RFU (normalized)",
    main = "", type = "b", pch = 19)

  mtext(paste0(LETTERS[i - 1], "    ", colnames(data)[i]), cex = 1,
        side = 3, adj = 0, font = 2)
  legend("topleft", paste0(ifelse(res_decision[i - 1, 2] == "n",
                                  "negative", "positive")),
         bty = "n")
  abline(res_q25_lm, col = colors[1])
  points(x[res_q25], y_tmp[res_q25], cex = 2.5, col = colors[1])
  abline(res_q75_lm, col = colors[2])
  points(x[res_q75], y_tmp[res_q75], cex = 2.5, col = colors[2])
}
```

Finally, the results of the analysis were printed in a tabular format.

```r
# Load the xtable library for an appealing table output
library(xtable)

# Analyze the data via the mblrr() function

res_mblrr <- do.call(cbind, lapply(2L:ncol(data), function(i) {
  suppressMessages(mblrr(
    x = data[, 1], y = data[, i],
    normalize = TRUE
  )) %>% data.frame()
```

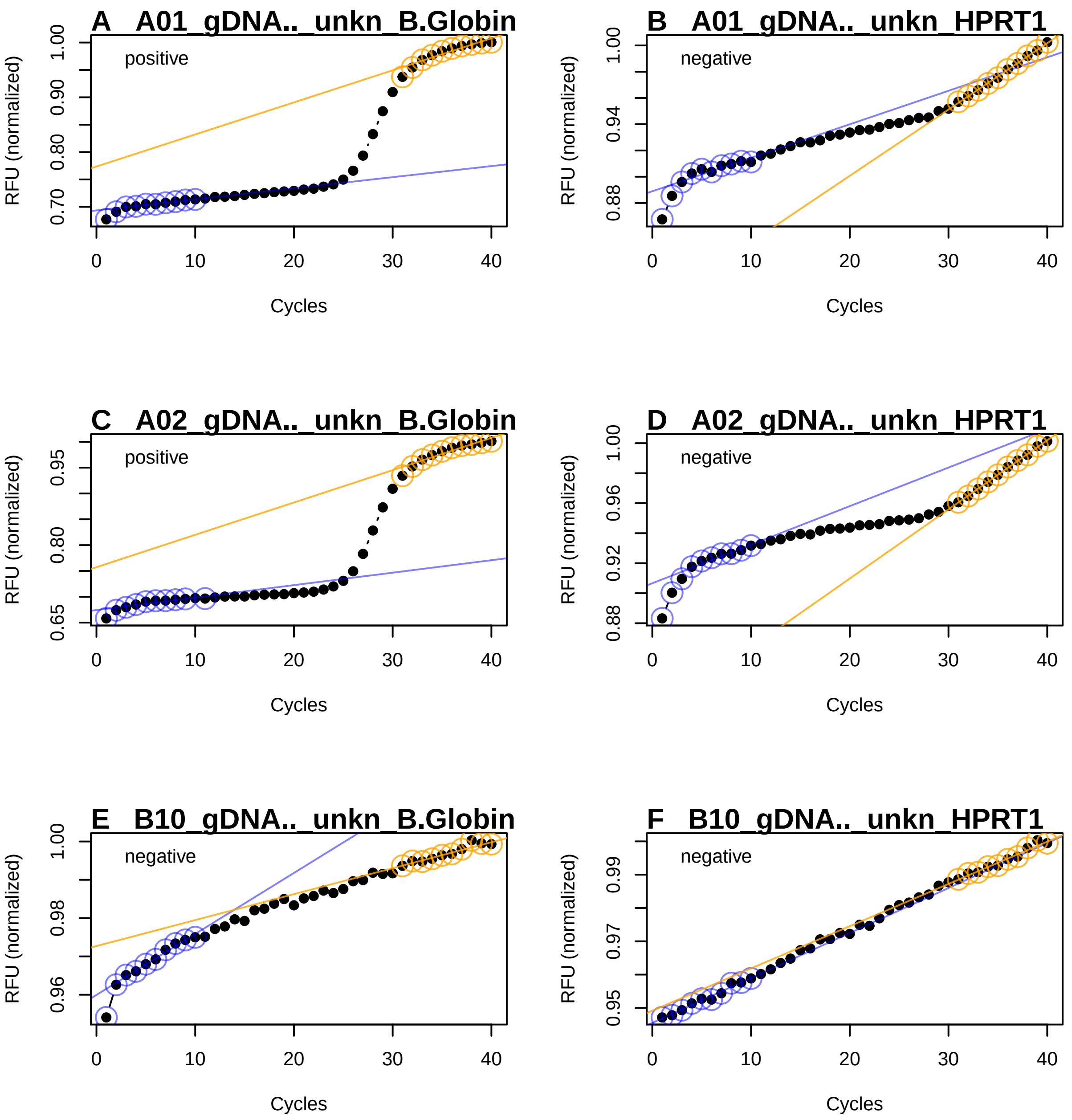

Figure 20: Robust local regression to analyze amplification curves. The amplification curves were arbitrarily selected from the ‘RAS002‘ data set. In the qPCR setup, the target genes beta globin (B. globin) and HPRT1 were simultaneously measured in a PCR cavity using two specific hydrolysis probes (duplex qPCR). Both positive (A, C, E) and negative (B, D, F) amplification curves were used. The amplification curves are normalized to the 99% quantile. The differences in slopes and intercepts (blue and orange lines and dots). The mblrr() function is presumably useful for data sets which are accompanied by noise and artifacts.

```r
}))
colnames(res_mblrr) <- colnames(data)[-1]

# Transform the data for a tabular output and assign the results to the object
# output_res_mblrr.

output_res_mblrr <- res_mblrr %>% t()

# The output variable names of the mblrr() function are rather long. For better
# readability the variable names were changed to "nBG" (intercept of the head
# region), "mBG" (slope of the head region), "rBG" (Pearson correlation of head
# region), "nTP" (intercept of the tail region), "mTP" (slope of tail region),
# "rBG" (Pearson correlation of the tail region)

colnames(output_res_mblrr) <- c(
  "nBG", "mBG", "rBG",
  "nTP", "mTP", "rTP"
)

print(xtable(
  output_res_mblrr, caption = "Selected results of predictors from the mblrr()
             function. nBG, intercept of head region; mBG, slope of head region;
             rBG, Pearson correlation of head region; nTP, intercept of tail
             region; mTP, slope of tail region; rBG, Pearson correlation of
             tail region",
  label = "tablemblrrintroduction"
), comment = FALSE, caption.placement = "top")
```

Table 1: Selected results of predictors from the mblrr() function. nBG, intercept of head region; mBG, slope of head region; rBG, Pearson correlation of head region; nTP, intercept of tail region; mTP, slope of tail region; rBG, Pearson correlation of tail region

| | nBG | mBG | rBG | nTP | mTP | rTP |
|---|---|---|---|---|---|---|
| A01_gDNA.._unkn_B.Globin | 0.69 | 0.00 | 0.91 | 0.77 | 0.01 | 0.94 |
| A01_gDNA.._unkn_HPRT1 | 0.89 | 0.00 | 0.87 | 0.80 | 0.01 | 1.00 |
| A02_gDNA.._unkn_B.Globin | 0.67 | 0.00 | 0.88 | 0.76 | 0.01 | 0.95 |
| A02_gDNA.._unkn_HPRT1 | 0.91 | 0.00 | 0.90 | 0.82 | 0.00 | 1.00 |
| B10_gDNA.._unkn_B.Globin | 0.96 | 0.00 | 0.95 | 0.97 | 0.00 | 0.96 |
| B10_gDNA.._unkn_HPRT1 | 0.95 | 0.00 | 0.99 | 0.95 | 0.00 | 0.98 |

###### 4.1.13 `autocorrelation_test()` - A Function to Detect Positive Amplification Curves

Autocorrelation analysis is a technique that is used in the field of time series analysis. It can be used to reveal regularly occurring patterns in one-dimensional data (Spiess et al. 2016). Autocorrelation measures the correlation of a signal $f(t)$ with itself shifted by some time delay $f(t-\tau)$.

The `autocorrelation_test()` function coerces the amplification curve data to an object of class "zoo" (`zoo` package) as indexed totally ordered observations. Then follows the computation of a lagged version of the amplification curve data. The shifting of the amplification curve data is based on the number of observations (number of cycles 'c') with the following $\tau$.

| Number of Cycles (c) | $\tau$ |
|---|---|
| $c \leq 35$ | 8 |

| Number of Cycles (c) | $\tau$ |
|---|---|
| $35 > c \leq 40$ | 10 |
| $40 < c \leq 45$ | 12 |
| $c > 45$ | 14 |

This is followed by a significance test for correlation between paired observations (original & lagged amplification curve data). The hypothesis is that paired observations of positive amplification curves exhibit significant correlation (`stats::cor.test`, significance level is 0.01) in contrast to negative amplification curves (noise). The application of the `autocorrelation_test()` function is shown in the following example.

In addition, the decisions file `decision_res_RAS002.csv` from the user was analyzed for the most frequent decision (modus) using the `decision_modus()` function (subsubsection 4.3.3).

```
## dec
##   a   n   y
##   1 202 279
```

```
## dec
##   n   y
## 203 279
```

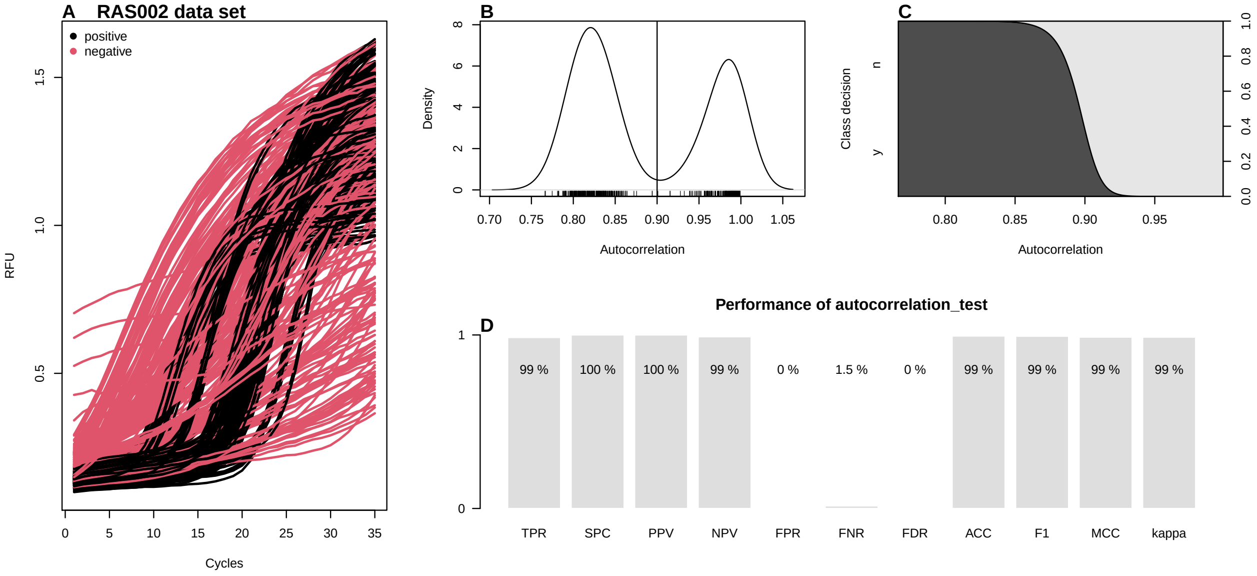

Figure 21: Autocorrelation analysis of the amplification curves of the ‘RAS002‘ data set. A) Display of all amplification curves of the data set ‘RAS002‘. Negative amplification curves are shown in red and positive amplification curves in black. The autocorrelation_test() function was used to analyze all amplification curves. B) The density diagram of the autocorrelation of positive and negative amplification curves shows a bimodal distribution. C) The cdplot calculates the conditional densities of x based on the values of y weighted by the boundary distribution of y. The densities are derived cumulatively via the values of y. The probability that the decision is negative (n) when autocorrelation equals 0.85 is approximately 100%. D) Performance analysis using the performeR() function (see subsubsection 4.1.1 for details).

As shown in this example, the `autocorrelation_test()` function is able to distinguish between positive and negative amplification curves. Negative amplification curves were in all cases non-significant. In contrast, the coefficients of correlation for positive amplification curves ranged between 0.766 and 0.999 at a significance level of 0.01 and a lag of 3.

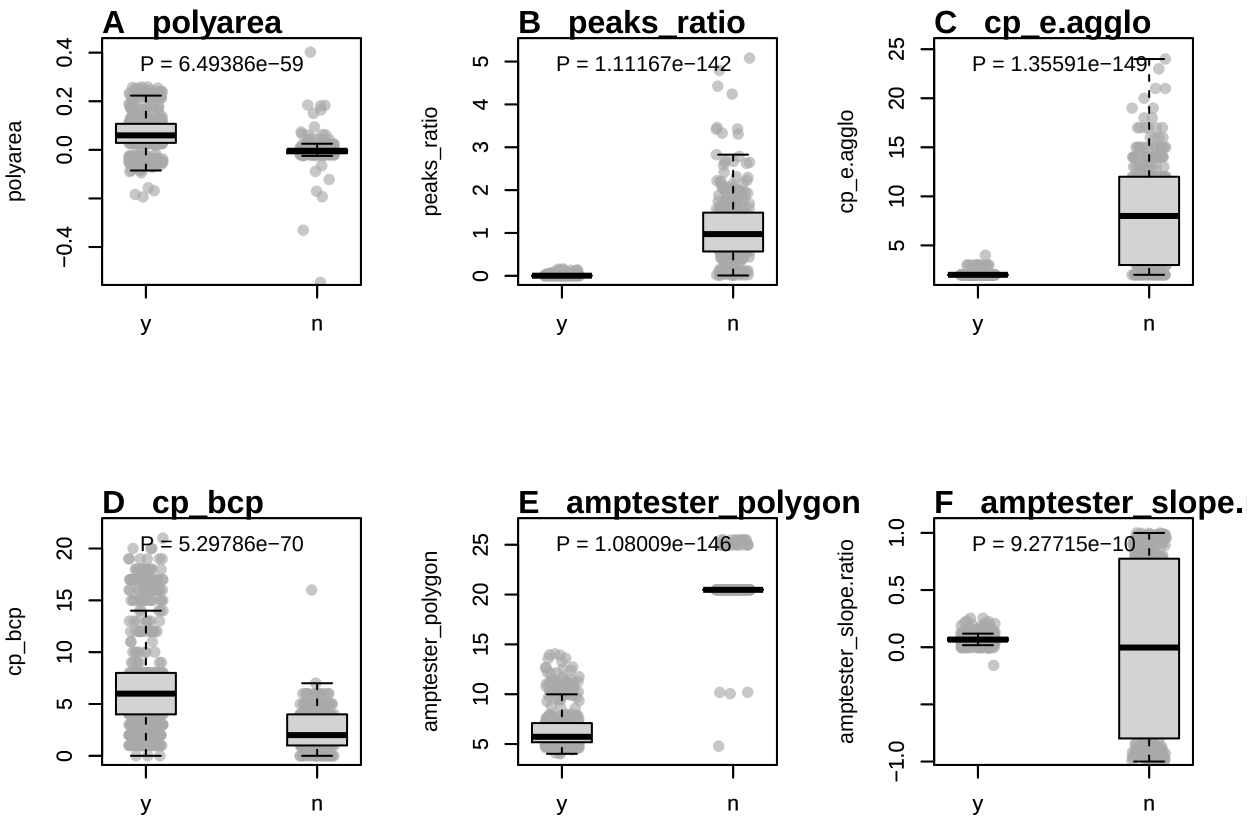

Figure 22: Values of predictors calculated from negative and positive amplification curves. Amplification curves predictors from the ‘data_sample_subset‘ data set were used as they contain positive and negative amplification curves and amplification curves that exhibit a *hook effect* or non-sigmoid shapes. A) ‘polyarea‘, is the area under the amplification curve determined by the Gauss polygon area formula. B) ‘peaks_ratio‘, is the ratio of the local minima and the local maxima. C) ‘cp_e.agglo‘, makes use of energy agglomerative clustering. Positive amplification curves have fewer change points than negative amplification curves. These two change point analyses generally separate positive and negative amplification curves. D) ‘cp_bcp‘, analyses change points by a Bayesian approach. Positive amplification curves appear to contain more change points than negative amplification curves. Nevertheless, there is an overlap between the positive and negative amplification curves in both methods. This can lead to false-positive or false-negative classifications. E) ‘amptester_polygon‘ is the cycle normalized order of a polygon. F) ‘amptester_slope.ratio‘ is the slope (linear model) of the raw fluorescence values at the approximate first derivate maximum, second derivative minimum and second derivative maximum.

###### 4.1.14 Frequentist and Bayesian Change Point Analysis

Change point analysis (CPA) encompasses methods to identify or estimate single or multiple locations of distributional changes in a series of data points indexed in time order. A change herein refers to a statistical property. CPA is used for example in econometrics and bioinformatics (Killick and Eckley 2014; Erdman, Emerson, and others 2007). Several change point algorithms exist, such as the binary segmentation algorithm (Scott and Knott 1974). In change point analysis one assumes independent ordered observations $x_1, x_2, \ldots, x_n \in \mathbb{R}^d$ (James and Matteson 2013). In the case of qPCR, this is simply the cycle-dependent fluorescence, used to create $k$ homogeneous subsets of unknown size (Erdman, Emerson, and others 2007). While frequentist methods make an estimation of the parameter at the location (e. g., mean) of the change points at specific points, change point analysis using Bayesian method produces a probability for the occurrence of a change point at certain points. For the analysis of the amplification curves, it was hypothesized that the number of change points differs between positive (sigmoidal) and negative (noise) amplification curves.

The `pcrfit_single()` function uses two independent approaches for change point analysis. These are the `bcp()` [`bcp`] (Erdman, Emerson, and others 2007) and the `e.agglo()` [`ecp`] function (James and Matteson 2013). The `e.agglo()` function performs a non-parametric change point analysis based on agglomerative hierarchical estimation and is useful to "detect changes within the marginal distributions" (James and Matteson 2013). Measurements from the qPCR systems typically show noise that has rapidly changing components. Differentiators amplify these rapidly changing noise components (Rödiger, Böhm, and Schimke 2013). Therefore, the first derivation of the amplification curve was used for both change point analyses. It was assumed for the change point analysis of amplification curves, that this leads to larger differences between positive and negative amplification curves. An example is shown on Figure 23. In contrast the `bcp()` [`bcp`] function performs a change point analysis based on a Bayesian approach. This method can detect changes in the mean of independent Gaussian observations. As a result, the analysis returns the posterior probability of a change point at each $x_i$. An example is shown on Figure 23. Both the change point analysis methods provide additional information to distinguish positive and negative amplification curves Figure 22E & F).

###### 4.1.15 `windower` - Analysis of multiple regions of interest

The `windower` feature is embedded in the `pcrfit_single()` function. It is a function to calculate the median absolute deviation of several sections of an amplification curve. Therefore, the amplification curve is divided into 10 windows. Each window always contains five data points (cycles) of RFU values, from which the median absolute deviation is calculated. To ensure this, each amplification curve is adjusted with a spline. From this spline a model is calculated from which exactly 50 points are calculated (interpolated), independent of the cycle range.

###### 4.1.16 Frequentist Approaches to Test the Class of an Amplification Reaction and Application of the `amptester()` Predictors

A part of `pcrfit_single()` is the `amptester()` [`chipPCR`] function, which contains tests to determine whether an amplification curve is positive or negative. The input values for the function differ due to the different preprocessing steps in the `pcrfit_single()` function. Therefore, the concepts of the tests are briefly described below.

- The first test, designated as SHt, is based on this Shapiro-Wilk test of normality. This relatively simple procedure can be used to check whether the underlying population of a sample (amplification curve) is significantly ($\alpha \leq 5e-04$) normal distributed. The name of the output of the `pcrfit_single()` function is `amptester_shapiro`.
- The second test is the *Resids growth test* (RGt), which tests if the fluorescence values in linear phase are stable. Whenever no amplification occurs, fluorescence values quickly deviate from linear model. Their standardized residuals will be strongly correlated with their value. For real amplification curves, the situation is much more stable. Noise (meaning deviations from linear model) in background do not correlate strongly with the changes in fluorescence. The decision is based on the threshold value (here 0.5). The output is binary coded (negative = 0, positive = 1). The output name of the `pcrfit_single()` function is `amptester_rgt`.

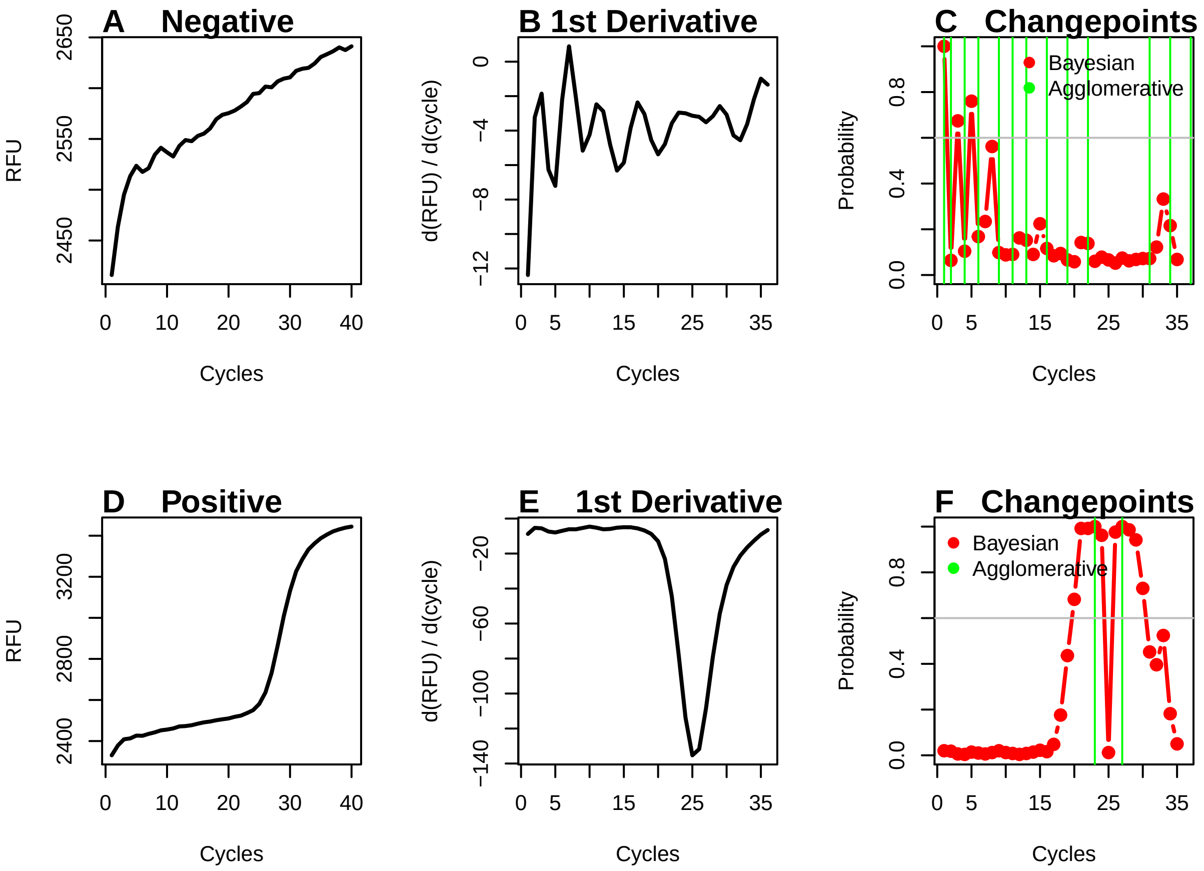

Figure 23: Bayesian and energy agglomerative change point analysis on negative and positive amplification curves. An analysis of a negative and a positive amplification curve from the ‘RAS002‘ data set was performed using the pcrfit_single() function. In this process, the amplification curves were analysed for change points using Bayesian change point analysis and energy agglomerative clustering. A) The negative amplification curve has a base signal of approximately 2450 RFU and only a small signal increase to 2650 RFU. There is a clear indication of the signal variation (noise). B) The first negative derivative amplifies the noise so that some peaks are visible. C) The change point analysis shows changes in energy agglomerative clustering at several positions (green vertical line). The Bayesian change point analysis rarely exceeds a probability of 0.6 (grey vert line). D) The positive amplification curve has a lower base signal ($\sim$ 2450 RFU) and increases up to the 40th cycle ($\sim$ 3400 RFU). A sigmoid shape of the curve is visible. E) The first negative derivation of the positive amplification curve shows a distinctive peak with a minimum at cycle 25. F) The change point analysis in energy agglomerative clustering shows changes (green vertical line) only at two positions. The Bayesian change point analysis shows a probability higher than 0.6 (grey horizontal line) at several positions.

- The third test is the *Linear Regression test* (LRt). This test determines the coefficient of determination ($R^2$) by an ordinary least squares linear (OLS) regression. The $R^2$ are determined from a run of $\sim$ 15% range of the data. If a sequence of more than six $R^2$s larger than 0.8 is found, a nonlinear signal is plausible. This is somewhat counter-intuitive, because $R^2$ of nonlinear data should be low. The output is binary coded (negative = 0, positive = 1). The output name of the `pcrfit_single()` function is `amptester_lrt`.
- The fourth test is called *Threshold test* (THt), based on the Wilcoxon rank sum test. As a simple rule the first 20% (head) and the last 15% (tail) of an amplification curve are used as input data. From this, a one-sided Wilcoxon rank sum tests of the head versus the tail is performed ($\alpha \leq 1e-02$). The output is binary coded (negative = 0, positive = 1). The output name of the `pcrfit_single()` function is `amptester_tht`.
- The fifth test is called *Signal level test* (SLt). The test compares the signals of the head and the tail by a robust "sigma" rule (median + 2 * MAD) and the comparison of the head/tail ratio. If the returned value is less than 1.25 (25 percent), then the amplification curve is likely negative. The output is binary coded (negative = 0, positive = 1). The output name of the `pcrfit_single()` function is `amptester_slt`.
- The sixth test is called *Polygon test* (pco). The pco test determines if the points in an amplification curve (like a polygon) are in a "clockwise" order. The sum over the edges result in a positive value if the amplification curve is "clockwise" and is negative if the curve is counter-clockwise. Experience states that noise is positive and "true" amplification curves are "highly" negative. In contrast to the implementation in the `amptester()` function, the result is normalized by a division to the number of PCR cycles. The output is numeric. The output name of the `pcrfit_single()` function is `amptester_polygon`.
- The seventh test is the *Slope Ratio test* (SlR). This test uses the approximated first derivative maximum, the second derivative minimum and the second derivative maximum of the amplification curve. Next, the raw fluorescence at the approximated second derivative minimum and the second derivative maximum are taken from the original data set. The fluorescence intensities are normalized to the maximum fluorescence of this data and then employed in a linear regression, using the estimated slope. The output is numeric and the output name of the `pcrfit_single()` function is `amptester_slope.ratio`.

#### 4.2 Classified Amplification Curve Datasets

Amplification curves from different sources (e. g., detection chemistries, thermo-cyclers) were manually classified with the `humanrater()` function (subsubsection 4.3.1) or with the `tReem()` function (subsubsection 4.3.2). Raw amplification curve data were exported as comma separated values or in the Real-time PCR Data Markup Language (RDML) format via the `RDML` package. RDML is human readable data exchange format for qPCR experiments. A detailed description can be found in Rödiger et al. (2017). The following code section describes the import of an RDML file from the `PCRedux` package. The RDML file contains amplification curve data of a duplex qPCR (HPV 16 & HPV 18) performed in the CFX96 (Bio-Rad).

At least the following datasets with a dichotomous decision (positive, negative) are included.

- decision_res_batsch1.csv
- decision_res_batsch2.csv
- decision_res_batsch3.csv
- decision_res_batsch4.csv
- decision_res_batsch5.csv
- decision_res_boggy.csv
- decision_res_C126EG595.csv
- decision_res_C127EGHP.csv
- decision_res_C316.amp.csv
- decision_res_C317.amp.csv
- decision_res_C60.amp.csv
- decision_res_CD74.csv
- decision_res_competimer.csv
- decision_res_dil4reps94.csv

- decision_res_guescini1.csv
- decision_res_guescini2.csv
- decision_res_HCU32_aggR.csv
- decision_res_htPCR.csv
- decision_res_karlen1.csv
- decision_res_karlen2.csv
- decision_res_karlen3.csv
- decision_res_lc96_bACTXY.csv
- decision_res_lievens1.csv
- decision_res_lievens2.csv
- decision_res_lievens3.csv
- decision_res_RAS002.csv
- decision_res_RAS003.csv
- decision_res_reps2.csv
- decision_res_reps384.csv
- decision_res_reps3.csv
- decision_res_reps.csv
- decision_res_rutledge.csv
- decision_res_stepone_std.csv
- decision_res_testdat.csv
- decision_res_vermeulen1.csv
- decision_res_vermeulen2.csv
- decision_res_VIMCFX96_60.csv

```
library(RDML)
# Load the RDML package and use its functions to import the amplification curve
#  data
library(RDML)
filename <- system.file("RAS002.rdml", package = "PCRedux")
raw_data <- RDML$new(filename = filename)
```

The following example shows the export of the `RAS002.rdml` file from the RDML format to the csv format.

```
# Export the RDML data from the PCRedux package as the objects RAS002 and RAS003.
library(RDML)
library(PCRedux)

RAS002 <- data.frame(RDML$new(paste0(
    path.package("PCRedux"), "/", "RAS002.rdml"))$GetFData()
    )

# The obbject RAS002 can be stored in the working directory as CSV file with
# the name RAS002_amp.csv.
write.csv(RAS002, "RAS002_amp.csv", row.names = FALSE)
```

| RDML data file | Device | Target gene | Detection chemistry |
|---|---|---|---|
| RAS002.rdml | CFX96 | HPV16, HPV18, HPRT1 | TaqMan |
| RAS003.rdml | CFX96 | HPV16, HPV18, HPRT1 | TaqMan |
| hookreg.rdml | Bio-Rad | various | TaqMan, DNA binding dyes |

Table 4: Classified amplification curve data sets. Decision data sets in `PCRedux`: table with results of manual classification as comma separated values. qPCR data set: name of original amplification curve data set. Package: name of the `R` package containing the amplification curves. Device: is the device used to measure the amplification reaction. **`Note: The original data sets contain information about the detection chemistry used within the corresponding qPCR experiments.`** AB, Applied Biosystems.

| Decision Datasets in PCRedux | qPCR Dataset | Package | Device |
|---|---|---|---|
| decision_res_RAS002.csv | RAS002.rdml | `PCRedux` | CFX96, Bio-Rad |
| decision_res_RAS003.csv | RAS003.rdml | `PCRedux` | CFX96, Bio-Rad |
| decision_res_batsch1.csv | batsch1 | `qpcR` | Light Cycler 1.0, Roche |
| decision_res_batsch2.csv | batsch2 | `qpcR` | Light Cycler 1.0, Roche |
| decision_res_batsch3.csv | batsch3 | `qpcR` | Light Cycler 1.0, Roche |
| decision_res_batsch4.csv | batsch4 | `qpcR` | Light Cycler 1.0, Roche |
| decision_res_batsch5.csv | batsch5 | `qpcR` | Light Cycler 1.0, Roche |
| decision_res_lc96_bACTXY.csv | lc96_bACTXY.rdml | `RDML` | Light Cycler 1.0, Roche |
| decision_res_boggy.csv | boggy | `qpcR` | Light Cycler 96, Roche |
| decision_res_C126EG595.csv | C126EG595 | `chipPCR` | Chromo4, Bio-Rad |
| decision_res_C127EGHP.csv | C127EGHP | `chipPCR` | iQ5, Bio-Rad |
| decision_res_C316.amp.csv | C316.amp | `chipPCR` | iQ5, Bio-Rad |
| decision_res_C317.amp.csv | C317.amp | `chipPCR` | iQ5, Bio-Rad |
| decision_res_C60.amp.csv | C60.amp | `chipPCR` | iQ5, Bio-Rad |
| decision_res_CD74.csv | CD74 | `chipPCR` | iQ5, Bio-Rad |
| decision_res_competimer.csv | competimer | `qpcR` | Light Cycler 480, Roche |
| decision_res_dil4reps94.csv | dil4reps94 | `qpcR` | CFX384, Bio-Rad |
| decision_res_guescini1.csv | guescini1 | `qpcR` | Light Cycler 480, Roche |
| decision_res_guescini2.csv | guescini2 | `qpcR` | Light Cycler 480, Roche |
| decision_res_htPCR.csv | htPCR | `qpcR` | Biomark HD, Fluidigm |
| decision_HCU32_aggR.csv | HCU32_aggR.csv | `PCRedux` | VideoScan |
| decision_res_karlen1.csv | karlen1 | `qpcR` | ABI Prism 7700, AB |
| decision_res_karlen2.csv | karlen2 | `qpcR` | ABI Prism 7700, AB |
| decision_res_karlen3.csv | karlen3 | `qpcR` | ABI Prism 7700, AB |
| decision_res_lievens1.csv | lievens1 | `qpcR` | ABI7300, ABI |
| decision_res_lievens2.csv | lievens2 | `qpcR` | ABI7300, ABI |
| decision_res_lievens3.csv | lievens3 | `qpcR` | ABI7300, ABI |
| decision_res_reps.csv | reps | `qpcR` | MXPro3000P, Stratagene |
| decision_res_reps2.csv | reps2 | `qpcR` | MXPro3000P, Stratagene |
| decision_res_reps3.csv | reps3 | `qpcR` | MXPro3000P, Stratagene |
| decision_res_reps384.csv | reps384 | `qpcR` | CFX384, Bio-Rad |
| decision_res_rutledge.csv | rutledge | `qpcR` | Opticon 2, MJ Research |
| decision_res_stepone_std.csv | stepone_std | `RDML` | StepOne, AB |
| decision_res_testdat.csv | testdat | `qpcR` | Light Cycler 1.0, Roche |
| decision_res_vermeulen1.csv | vermeulen1 | `qpcR` | Light Cycler 480, Roche |
| decision_res_vermeulen2.csv | vermeulen2 | `qpcR` | Light Cycler 480, Roche |
| decision_res_VIMCFX96_60.csv | VIMCFX96_60 | `chipPCR` | CFX96, Bio-Rad |
| decision_res_kbqPCR | kbqPCR | `PCRedux` | CFX96, Bio-Rad |

#### 4.3 Technologies for Amplification Curve Classification and Classified Amplification Curves

Many machine learning concepts exist. One method is supervised machine learning, where the goal is to derive a property from user-defined (classified) training data. Categories such as negative, ambiguous or positive are assigned depending on the form of the amplification curve. An extensive literature research showed that there are no openly accessible classified amplification curve data sets. Open Data is meant in the sense that data are freely available, free of charge, free to use and that data can be republished, without restrictions from copyright, patents or other mechanisms of control (Kitchin 2014).

Therefore, a large number of records with amplification curves and their classification (negative, ambiguous, positive) were added to the `PCRedux` package.

The following is a list of where the labeling data records with the classes negative, ambiguous and positive can be found and called up. In order to save storage capacity, the original amplification data sets were left in the corresponding R-packages. All data are freely available under open source licenses. Note: The **`miRcompData`** package https://bioconductor.org/packages/release/data/experiment/html/miRcompData.html ((Kemperman and McCall 2017)) contains a very large set of gene expression data. However, no label data is available for this one.

For the amplification curves in Table 4, a dichotomous classification was performed (roughly sigmoid or negative amplification reaction with a flat curve shape). Consequently, this does not rule out

- if an unspecific amplification product has been synthesized,
- if a contamination has been amplified or
- if only primer-dimers have been amplified.

To answer this question, other methods such as agarose gel electrophoresis need to be used.

##### 4.3.1 Manual Amplification Curve Classification

For machine learning and method validation, it was important to classify the amplification curves individually. In Rödiger, Burdukiewicz, and Schierack (2015), the **`humanrater()`** [**`chipPCR`**] function was described. This function was developed to help the user during the classification of amplification curves and melting curves. The user has to define classes, which get assigned to an amplification curve after an expert has entered the class in input mask.

> By convention, class labels were specified as e. g., negative ("n"), ambiguous ("a"), positive ("y") in the `PCRedux` package.

All amplification curve data sets listed in Table 4 were classified in interactive, semi-blinded sessions. **`humanrater()`** [**`chipPCR`**] was set to randomly select individual amplification curves. All data sets were manually classified at least three times. The **`htPCR`** data set (Figure 4B) was classified eight times in total (see Figure 24). Most of the amplification curves are neither unequivocal classifiable as positive or negative.

```r
# Suppress messages and load the packages for reading the data of the classified
# amplification curves.
library(PCRedux)

# Load the decision_res_htPCR.csv data set from a csv file.
filename <- system.file("decision_res_htPCR.csv", package = "PCRedux")
decision_res_htPCR <- read.csv(filename)

# Create graphic device for the plot(s)
par(mfrow = c(2, 4))
for (i in 2L:9) {
    data_tmp <- table(as.factor(decision_res_htPCR[, i]))
```

```
    barplot(data_tmp, col = adjustcolor("grey", alpha.f = 0.5),
            xlab = "Class", ylab = "Counts", border = "white")
    text(c(0.7, 1.9, 3.1), rep(quantile(data_tmp, 0.25), 3),
         data_tmp, srt = 90)
    mtext(LETTERS[i - 1], cex = 1, side = 3, adj = 0, font = 2)
}
```

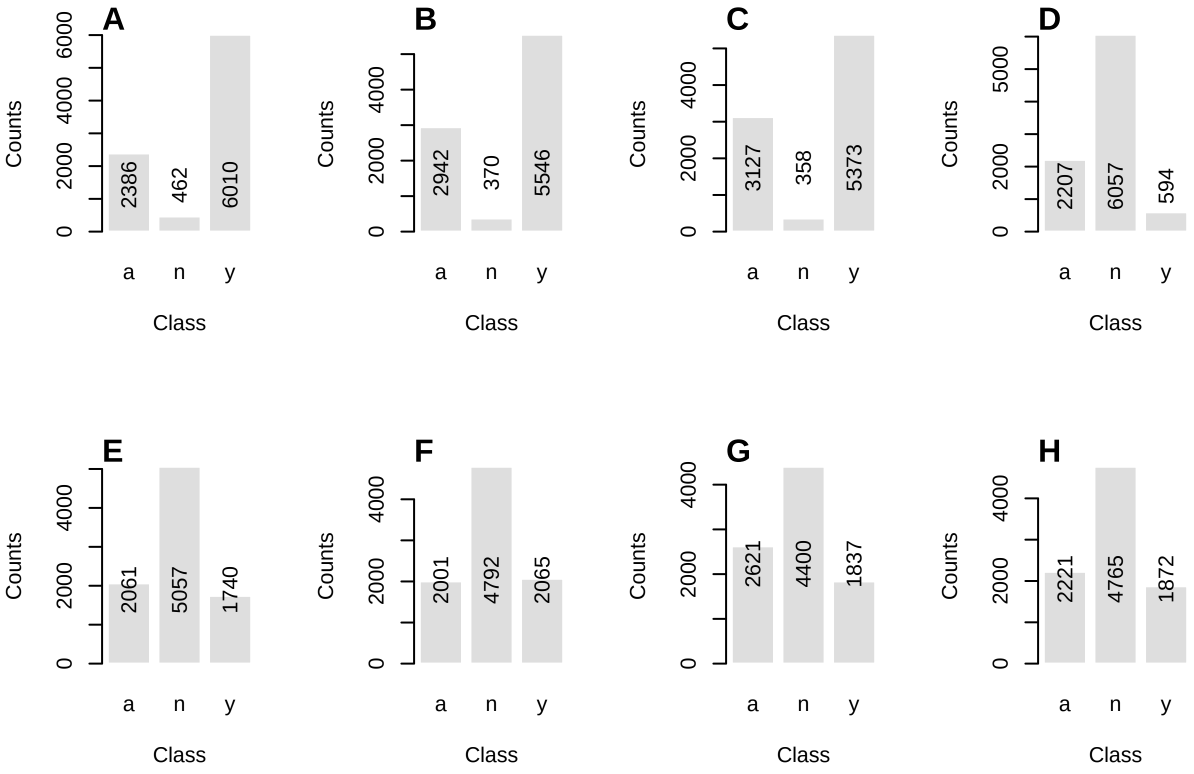

Figure 24: Variations in the classification of amplification curves. A prerequisite for the development of machine-learning models is the availability of manually classified amplification curves. Amplification curves (n = 8858) from the 'htPCR' data set have been classified by one user eight times at different points over time (classes: ambiguous (a), positive (y) or negative (n)). During this process, the amplification curves were presented in random order. The example shows that different (subjective) class mappings may occur for the same data set. While only a few amplification curves were classified as negative in the first three classification cycles (A-C), their proportion increased almost tenfold in later classification cycles (D-H).

This approach is well suited and has been applied to classify a variety of amplification curves during the development of the `PCRedux` package. From experience this is time-consuming and tiring for large data sets, especially when the amplification curves are similar in shape. A high similarity between amplification curves exists, for example, in replicates and negative controls.

###### 4.3.2 `tReem()` - A Function for Shape-based Group-wise Classification of Amplification Curves

The similarity of amplification curves can be used to form groups of similar shapes. The amplification curves in the groups can then be classified in bulk. In this way, a higher throughput can be achieved, a concept yet not described for the analysis of qPCR data in the literature.

The `tReem()` function was developed to perform a *shape-based group classification*. To use the `tReem()` function, the first column must contain the qPCR cycles and all subsequent columns must contain the amplification curves. Two measures of similarity are used within the `tReem()` function.

- In the first measure (default), the Pearson *correlation coefficients* ($r$) are determined in pairs for all combinations of the amplification curves. The correlation coefficient is a statistical measure to describe the strength of the correlation between two or more variables. The correlation coefficient $r$ is regarded as distance between the amplification curves. $r$ is a dimensionless value and only takes values between -1 and 1. If $r = -1$, there is a maximum reciprocal relationship. If $r = 0$ there is no correlation between the two variables. If $r = 1$, there is a maximum rectified correlation.

- In the second measure, the *Hausdorff distance* is used to determine the similarity between amplification curves. The Hausdorff distance is "the maximum of the distances from a point in any of the sets to the nearest point in the other set" (Rote 1991; Herrera et al. 2016). The amplification curves are converted within the `tReem()` function using the `qPCR2data()` function.

Both methods process the distances in the same steps. This involves the calculation of the distance matrix using the Euclidean distances of all distance measures to determine the distance between the lines of the data matrix.

This is used to perform a hierarchical cluster analysis. In the last step, the cluster is divided into groups based on a user-defined $k$ value. For example, two groups are created for $k = 2$. If the amplification curves shapes are highly diverse, a larger $k$ should be used. After a chain of processing steps presents the `tReem()` function a series of plots with grouped of amplification curves. The corresponding classes can then be assigned to the groups of amplification curves by the user using an input mask.

Grouping the amplification curves with the Pearson correlation coefficient as a distance measure is usually faster than the Hausdorff distance. The Hausdorff distance is an approximation of a shape metrics to define similarity measures between shapes. (Charpiat, Faugeras, and Keriven 2003).

```
# Classify amplification curve data by correlation coefficients (r)
data <- qpcR::testdat

classification_result <- tReem(data[, 1:15], k = 3)
classification_result
```

###### 4.3.3 `decision_modus()` - A Function to Get a Decision (Modus) from a Vector of Classes

For the systematic statistical analysis of classification data sets, the `decision_modus()` function has been developed. This allows the most common decision (mode) to be determined. The mode is useful to consolidate large collections of different decisions into a single (most frequent) decision.

> Observed:$a$, $a$, $a$, $a$, $a$, $n$, $n$, $n$ → frequencies 5 x $a$, 3 x $n$ → mode: $a$. Since the class names are known, they only have to be interpreted by the user (e. g., "a",$n$,"y" -> "ambivalent","negative","positive").

A manual classification was performed out for the `htPCR` data set (for an example plot Figure 4B) with the `humanrater()` function. The classification of each amplification curve was performed eight times at different time points since many of the amplification curves did not resemble optimal curvatures (e. g., Figure 1). It is likely that the amplification curve (P06.W47, Figure 1) is considered as ambiguous or even positive (positive ↔ ambivalent) by the users.

Table 5 shows from a total of 8858 amplification curves the first 25 lines classified as negative (*conformity=TRUE*) and the first 25 lines classified as positive. Since in total, the curves were classified eight times (`test.result.1` … `test.result.8`) a whole of 70864 amplification curves was analysed. In this classification experiment the amplification curves have been classified differently in 94.6% of the cases (e. g., line 1 "P01. W01").

The `decision_modus()` function was applied to the record `decision_res_htPCR.csv` with all classification rounds (columns 2 to 9) and the mode was determined for each amplitude curve Figure 25.

```
# Use decision_modus() to go through each row of all classifications done by
# a human.
```

Table 5: Results of the ‘htPCR‘ data set classification. All amplification curves of the ‘htPCR‘ data set were classified as ‘negative‘, ‘ambiguous‘ and ‘positive‘ by individuals in eight analysis cycles (‘test.result.1‘ … ‘test.result.8‘). If an amplification curve was always classified with the same class, the last column (‘conformity‘) shows ‘TRUE‘. As an example, the table shows 25 amplification curves with consistent classes and 25 amplification curves with differing classes (‘conformity = FALSE‘).

| htPCR | test.result.1 | test.result.2 | test.result.3 | test.result.4 | test.result.5 | test.result.6 | test.result.7 | test.result.8 | conformity |
|---|---|---|---|---|---|---|---|---|---|
| P01.W01 | y | a | a | n | n | n | a | n | FALSE |
| P01.W02 | a | y | a | n | n | n | n | n | FALSE |
| P01.W03 | y | a | a | n | n | n | n | n | FALSE |
| P01.W04 | y | a | a | n | n | n | n | n | FALSE |
| P01.W05 | y | a | a | n | n | n | a | n | FALSE |
| P01.W06 | a | a | a | n | n | n | n | n | FALSE |
| P01.W07 | a | a | a | n | n | n | n | n | FALSE |
| P01.W08 | y | a | a | n | n | n | n | n | FALSE |
| P01.W09 | y | a | a | n | n | n | n | n | FALSE |
| P01.W10 | n | a | a | n | n | n | n | n | FALSE |
| P01.W11 | y | y | a | y | y | y | a | y | FALSE |
| P01.W12 | a | a | a | n | n | n | n | n | FALSE |
| P01.W13 | y | a | y | n | n | n | n | n | FALSE |
| P01.W14 | y | a | y | n | n | n | n | n | FALSE |
| P01.W15 | y | a | y | n | n | n | n | n | FALSE |
| P01.W16 | y | a | a | n | n | n | n | n | FALSE |
| P01.W17 | y | a | a | n | n | n | n | n | FALSE |
| P01.W18 | a | a | a | n | n | n | n | n | FALSE |
| P01.W19 | y | a | a | n | n | n | a | n | FALSE |
| P01.W20 | y | a | a | y | a | a | y | a | FALSE |
| P01.W21 | a | a | a | n | n | n | n | n | FALSE |
| P01.W22 | y | a | a | n | n | n | n | n | FALSE |
| P01.W23 | y | a | a | n | n | n | n | n | FALSE |
| P01.W24 | y | a | a | n | n | n | a | n | FALSE |
| P01.W25 | y | a | a | n | n | n | n | n | FALSE |
| P01.W58 | n | n | n | n | n | n | n | n | TRUE |
| P02.W09 | y | y | y | y | y | y | y | y | TRUE |
| P02.W19 | y | y | y | y | y | y | y | y | TRUE |
| P02.W31 | y | y | y | y | y | y | y | y | TRUE |
| P02.W41 | y | y | y | y | y | y | y | y | TRUE |
| P02.W72 | y | y | y | y | y | y | y | y | TRUE |
| P02.W81 | y | y | y | y | y | y | y | y | TRUE |
| P02.W84 | n | n | n | n | n | n | n | n | TRUE |
| P03.W09 | y | y | y | y | y | y | y | y | TRUE |
| P03.W21 | y | y | y | y | y | y | y | y | TRUE |
| P03.W22 | y | y | y | y | y | y | y | y | TRUE |
| P03.W31 | y | y | y | y | y | y | y | y | TRUE |
| P03.W32 | y | y | y | y | y | y | y | y | TRUE |
| P03.W39 | y | y | y | y | y | y | y | y | TRUE |
| P03.W56 | y | y | y | y | y | y | y | y | TRUE |
| P03.W59 | y | y | y | y | y | y | y | y | TRUE |
| P03.W68 | y | y | y | y | y | y | y | y | TRUE |
| P03.W72 | y | y | y | y | y | y | y | y | TRUE |
| P03.W73 | y | y | y | y | y | y | y | y | TRUE |
| P04.W19 | y | y | y | y | y | y | y | y | TRUE |
| P04.W31 | y | y | y | y | y | y | y | y | TRUE |
| P04.W66 | y | y | y | y | y | y | y | y | TRUE |
| P04.W81 | y | y | y | y | y | y | y | y | TRUE |
| P04.W91 | y | y | y | y | y | y | y | y | TRUE |
| P05.W20 | y | y | y | y | y | y | y | y | TRUE |

```r
# Determine the number of observations where all classifications were
# the same (conformity == TRUE).
conformity <- decision_res_htPCR[["conformity"]]

# List all classifications.
dec <- lapply(1L:nrow(decision_res_htPCR), function(i) {
    decision_modus(decision_res_htPCR[i, 2:9])[1]
}) %>% unlist()

# Show statistic of the decisions
summary(dec)
```

```
##    Min. 1st Qu.  Median    Mean 3rd Qu.    Max.    NA's
##   1.000   1.000   2.000   1.779   2.000   3.000    1179
```

Another usage mode of `decision_modus()` is to set the parameter as `max_freq=FALSE`. This option specifies the number of all classifications.

```r
library(PCRedux)
# Decisions for observation P01.W06
res_dec_P01.W06 <- decision_modus(decision_res_htPCR[
which(decision_res_htPCR[["htPCR"]] == "P01.W06"),
```

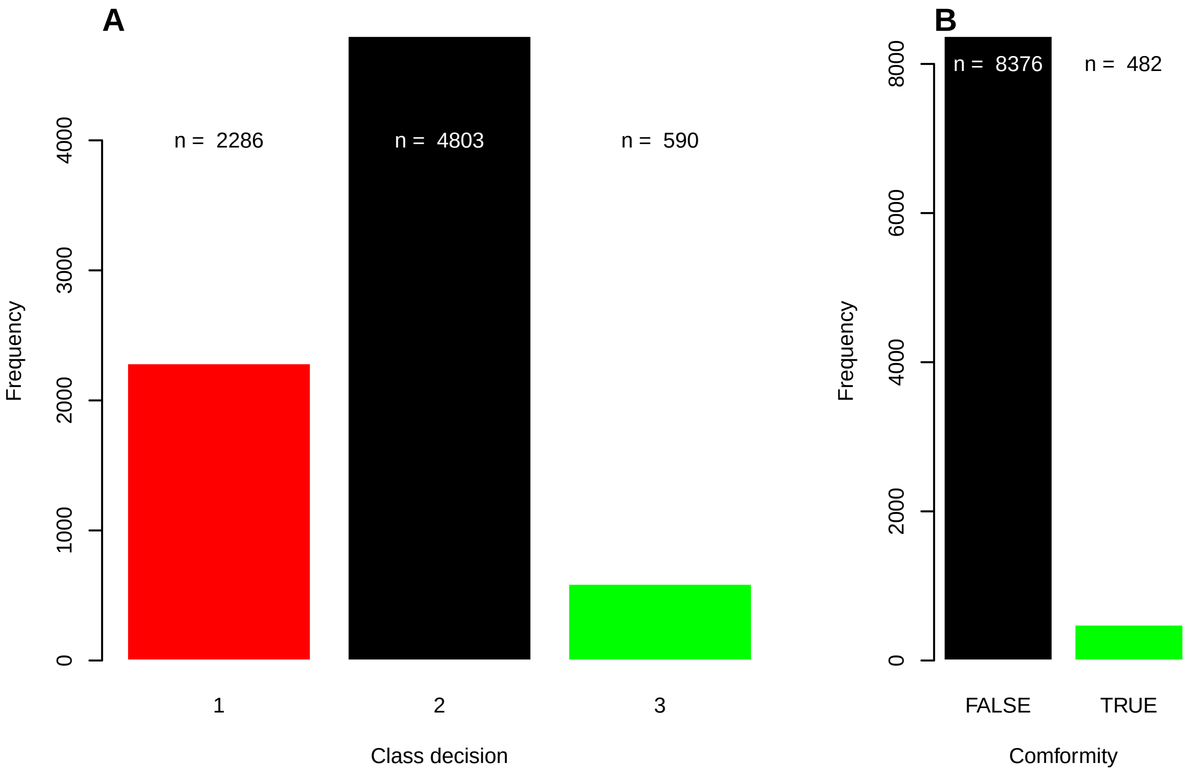

Figure 25: Frequency of amplification curve classes and conformity in the ‘htPCR‘ data set. The ‘htPCR‘ data set was classified by hand eight times. Due to the unusual amplification curve shape and input errors during classification, many amplification curves were classified differently. A) Frequency of negative (black), ambiguous (red) and positive (green) amplification curves in the ‘htPCR‘ data set. The combined number of ambiguous and negative amplification curves appears to be higher, than the number of positive amplification curves. B) The number of observations where all classification cycles made the same decision (conformity == TRUE) accounts for only 5% of the total number of observations. TRUE, all classes of the amplification curve matched. FALSE, at least one in eight observations had a different class.

```
                                        2L:9
], max_freq = FALSE)
print(res_dec_P01.W06)
```

```
##   variable freq
## 1        a    3
## 2        n    5
```

The amplification curve P01. W06 was classified as a=3 times and as n=5 times. Therefore, the decision would turn negative.

##### 4.3.4 PCRedux-app

PCRedux-app is a web server, based on the shiny technology (Chang et al. 2019) wrapped around the encu() function (subsubsection 4.1.2). An user can upload qPCR data and download obtained amplification curve features.

There are different ways to use the function.

- Through RScript (Scripting Front-End for R):
  - Enter the command Rscript -e 'PCRedux::run_PCRedux()' in a console and copy the pasted URL in a browser.
- Through Graphical User Interfaces:
  - The function can be started directly in RStudio or RKWard by:

```
# run the Shiny app
PCRedux::run_PCRedux()
```

### 5 Summary and Conclusions

Extensive amounts of data represent a serious challenge in the analysis of qPCR amplification curves. In a manual classification (e. g. negative, positive), the result is usually characterized by the subjective perception of the experimenter. In addition, the time required for a manual analysis is high. An automatic system for amplification curve analysis might objectify and generalize the decision process. Interestingly, so far most authors focused on the extraction of single predictors from amplification curves. These are mainly the Cq and the amplification efficiency, which are used for the downstream processing such as expression analysis or genotyping (Pabinger et al. 2014).

Numerous software tools were developed, which deal with theses analytical steps. For example Baebler et al. (2017) published quantGenius and Mallona et al. (2017) published Chainy. However, none of them attempts to make use of characteristics of the amplification curve. Especially as this shape relies on the used chemistry and fluorochrome system (Ruijter et al. 2014). Even the cycler used may impact the amplification curve shape (Spiess et al. 2016). Positive amplification curves usually exhibit a sigmoid shape, consisting of a ground phase, exponential phase, and plateau phase. Negative amplification curves resemble flat noisy signal. As a result, the experienced user is usually able to correctly interpret the curve shape. Similarly, outliers and measurement errors can also be identified. When setting up a qPCR assay, manual data analysis is a useful and necessary approach to familiarize oneself with the properties of the qPCR amplification curves. To sum it up in the main document we show that the shape has features that a seen by machine learning models (Random Forest).

It is challenging to analyze and classify amplification curves if they deviate substantially from the sigmoid shape or if their number is no longer feasible for manual analysis. Furthermore, the supposed objectivity of the user must also be questioned. In the scientific environment there is often the temptation - or rather the compulsion - to use all data for publications. As a result, amplification curves of rather poor quality may be provided that are not reproducible and suitable for an objective analysis of amplification curves. For a novice user, the quality of an amplification curve can be acceptable, for an experienced user, not.

Ambiguous amplification curves are a big challenge for the user, as both classes (positive and negative) can be true. In most cases, however, the user is interested in an automatic distinction. For example, between positive and negative samples. This is important for screening applications. In the **online supplement**, further reasons were elaborated on why mathematical/statistical methods are necessary for the objective and reproducible interpretation of the amplification curves.

In addition to the determination of quantification points, the classification of amplification curves is necessary. For example, a diagnostician is interested in whether a sample is positive or negative. In research using high-throughput screening methods, it is important to classify large data sets quickly and cost-effectively. It is important to bear in mind that in manual classification, the classification result is influenced by the subjective perception of the experimenter and that it is comparatively time-consuming.

Here, an automatic computer-assisted classification of amplification curves is feasible because it renders the entire analysis process faster, more objective and more reproducible. The objectives were therefore to

1. create a collection of classified amplification curve data,
2. propose algorithms that can be used to calculate predictors from amplification curves,
3. to develop pipelines (e. g., machine learning, decision trees) that can be used for automatic classification of amplification curves,
4. evaluate pipelines that can be used for an automatic classification of amplification curves based on the curve shape and
5. to bundle and distribute the findings in a public repository open source software and open data package with an open data.class

for an automatic analysis of amplification curve data by machine learning.

For this purpose, the `PCRedux` package was developed. This package contains proof-of-concept algorithms and functions with which predictors (mathematically describable properties) of amplification curves can be calculated. The `pcrfit_single()` function is an extensible wrapper function for all algorithms and functions developed. `PCRedux` version 1.0.7 offers concepts to calculate 90 predictors from an amplification curve. In addition, predictors such as the employed chemistry, the qPCR device and further experimental details can be added by the user. Other parameters (e. g. hydrolysis probe, DNA binding dye) can be converted into binary classifiers and be used for modeling. Machine learning requires predictors to train a model (Saeys, Inza, and Larranaga 2007). The model should then be able to put new unknown data into a meaningful context. Predictors can have a significant influence on the accuracy of the prediction model. The predictors of the `PCRedux` package are a novelty in the literature for the classification of amplification curves, so that the collection probably constitutes the most extensive one at the time of the first release on https://github.com/devSJR/PCRedux (summer 2017).

It can be assumed that not all predictors are suitable or necessary for machine learning. Some **potential** predictors are more likely to be suitable for validation of data integrity (quality management) and data mining. The `maxRFU` predictors and `sigma_bg` (Figure 16) are to be mentioned here as examples. The predictor `maxRFU` should ideally be at a value of 1. If this is not the case, it can be assumed that a preprocessing problem was encountered. The `sigma_bg` value describes the standard deviation of the ground phase. It should be low ($sigma\_bg \leq 0.1$), otherwise it can be assumed that unusually high variations of the intensity values are present. Consequently, the relevance of the predictors must be determined independently by each user on the basis of domain knowledge, the objective, and the given data set.

At this point it should be mentioned that these approaches were deliberately sought and developed to make the approach and implementation more understandable. Self-learning machine-learning methods are expected to be included in the `PCRedux` package in future releases, from the necessity to design and test additional predictors.

To a modest extent, the usefulness of the predictors was tested on exemplary data sets, with the aim of achieving a high degree of objectivity and reproducibility. It is probably not possible to fulfill this ideal completely, since the algorithms were designed from a limited human perspective, which poses a generic problem in machine learning. In particular, data records can be distorted if the user excludes seemingly problematic data. Hence, it is advisable to perform a more comprehensive analysis of all predictors.

Several ROIs (see Figure 5) can be obtained from amplification curves. Sigmoidal amplification curves have turning points that can serve as an indicator of a positive amplification curve, and are mathematical and statistical starting points for calculating predictors. Amplification curves can have unique trajectories and often deviate drastically from ideal sigmoid models (see Figure 4A and Figure 4B). Some amplification curves have only a slight increase with positive or negative signs lacking a sigmoid curvature.

Almost all real-time thermo-cyclers have built-in software that performs preprocessing steps such as smoothing, baseline correction and normalization on the amplification curves (Rödiger, Burdukiewicz, and Schierack 2015; Spiess et al. 2015, 2016). For this reason, it is nearly impossible to get access to informative raw data, so that the impact on the predictor extraction process cannot be adequately estimated.

The volume and classification of data sets needs to be representative (Herrera et al. 2016). The `PCRedux` package contains a large number of manually classified amplification curves (subsection 4.3). Here, data preparation is an important step,as it encompasses data cleansing, data transformation and data integration (Herrera et al. 2016). To assist, the `xray` package (Seibelt 2017) and `assertr` package (Fischetti 2019) can be used to analyze the distribution and variables in records for important anomalies such as missing values, zeros, empty strings (Blank) and infinite numbers (Inf). Users of the`PCRedux` package should use such tools before proceeding with the analysis. Even though the data records (qPCR runs) in the `PCRedux` package have a very similar structure, some records contain missing values or have different dimensions. For example, the data set `C127EGHP` comprises a matrix of 40 x 66 (35 cycles x observations (65 amplification curves)), while the data set `htPCR` comprises a matrix of 35 x 8859. Talking about data sets, it is important to make sure that the volume of the data is representative. In this study amplification curves were categorized by hand and processed with the algorithms described in this study. Although this data set is fairly large in comparison to what existed before, there is no numeric evidence how well it reflects amplification curves in general. In particular, not all data sets have comparable case numbers. For example, the `htPCR` data set (Biomark HD, Fluidigm) encompass in total 8858 amplification curves, while the `C127EGHP` data set (iQ5, Bio-Rad) encompass in total 64 amplification curves. While most of the amplification curves of the `C127EGHP` data set have a classical sigmoid curve shape, amplification curves from the `htPCR` data set largely display noisy curvatures with non-sigmoid shape. One may question if the larger data set introduces a confirmation bias.

Although labeling and amplification curve data sets in the `PCRedux` package are large (see subsection 4.3 how to obtain the open data (open source licenced)) compared to what has been previously available, there is no information how well these represent amplification curves in other settings. Bellman coined the term "Curse of Dimensionality" in 1961 when he dealt with adaptive control processes, vaguely describing the practical difficulties of high-dimensional analysis and estimation. There is only a maximum number of predictors for a sample of a certain size. If there are too many, the performance of an algorithm decreases rather than improves. As a result, many data mining algorithms fail with high dimensionality because the data points are sparse (Herrera et al. 2016). In addition, the following applies:

- The user's knowledge, prejudices, and skills are reflected in the classified data. For example, an amplification can be classified as "ambiguous" by one and "positive" by another user.
- Not all data sets have a comparable number of cases as described above. The most of the amplification curves of the `C127EGHP` data set have an *ideal* sigmoid waveform. In contrast, the amplification curves from the `htPCR` data set are noisy and difficult to classify.
- The user decides
  - how the data is preprocessed,
  - which predictors are determined by the amplification curves,
  - which data set is used for machine learning and to what extent,
  - how the models are tested and
  - which results are reported.

Accordingly, the domain knowledge, biases and competences of the human operator are reflected by the software. For example, amplification that are classified by one human as 'ambiguous' might be classified as 'positive' by another human operator. This will affect (bias) the characteristics of the training data set. The model is intended to be in accordance with the human operator. The implications can be problematic. For example, amplification curves rated as false negative might lead to an adverse evidence in a forensic setting. As illustrated in subsection 2.3, every human operator will make an association between the shape of an amplification curve and the class it belongs to.

Measures to minimize errors in manual classification include the implementation of algorithms to detect them. This should preferably be done in the form of open source packages with publicly accessible data sets. In contrast to black box algorithms and hidden data sets, third parties can review and modify all elements. Many qPCR devices have built-in software that performs preprocessing steps like smoothing, base-lining and normalization and the data sets (Rödiger, Burdukiewicz, and Schierack 2015; Spiess et al. 2015, 2016). This will have an impact on the predictor extraction process. Same applies to the preprocessing steps in the `PCRedux` package, which have not been studied thoroughly.

For the examples in the `PCRedux` package, the question could not be clarified whether class imbalances in the data sets cause a confirmation distortion. However, a class imbalance may result in loss of prediction strength, because some classifiers make the assumption of similar class distributions (Herrera et al. 2016).

A new concept for the fast group-wise classification of amplification curves was introduced within the `PCRedux` package. Based on experience with the collected data sets it can be stated that only a few iterations are necessary to classify a large data set. This part of the software is intended as an assistance tool for the users of the package.

Measures to minimize these sources of error are based on the implementation of algorithms in the form of an open source package, and the publication of the data sets. Unlike black box algorithms and inaccessible data sets, every user can check and correct all steps. However, it is advised that users of the `PCRedux` package verify independently whether their models are objective. Failure to do so may have severe implications, for instance, amplification curves classified as false negative may lead to misleading inferences in a forensic analysis.

Machine-learning algorithms require careful data preprocessing and quality management. In a first step, relatively large data sets of known characteristic vectors have to be collected and depending on the machine learning approach, predictors calculated. In a second step, these characteristics are used to classify unknown characteristic vectors using the automatic learning algorithm. As an example, the amplification curves would need to be randomly split into training data and test data. Some examples were used in the `PCRedux` package to show how predictors of an amplification curve data set can be calculated and used for classifications.

In this study, it was shown how a given data set of amplification curves with known classifications can be used to build a system that can predict the classification of amplification curves. The algorithms provide means for a sensitive and specific classification of amplification curves from qPCR experiments in both supervised an unsupervised analysis mode. However, this method might be applicable to melting analysis too. This needs to be investigated in further studies.

Importantly, the concepts elucidated in this work may also be applied to other bioanalytical methods (e. g. enzyme kinetics, receptor binding studies, ELISA results, biological growth curves) with sigmoidal structure, however this requires more detailed interrogation. The `PCRedux` software may also be coupled with other technologies like Next Generation Sequencing. qPCR is used for pretesting (DNA quality) and as a confirmation test for RNA-Seq quantification (Nassirpour et al. 2014). For this purpose, automated quality control and decision support are conceivable.

Hothorn, Torsten, and Brian S. Everitt. 2014. *A Handbook of Statistical Analyses using R, Third Edition.* 3rd ed. Oakville: Chapman; Hall/CRC.

Isaac, Peter G. 2009. "Essentials of nucleic acid analysis: a robust approach." *Annals of Botany* 104 (2): vi–vi. https://doi.org/10.1093/aob/mcp135.

James, Nicholas A., and David S. Matteson. 2013. "ecp: An R package for nonparametric multiple change point analysis of multivariate data." *arXiv Preprint arXiv:1309.3295.* https://arxiv.org/abs/1309.3295.

Kemperman, Lauren, and Matthew N. McCall. 2017. "miRcomp-Shiny: Interactive assessment of qPCR-based microRNA quantification and quality control algorithms." *F1000Research* 6 (November): 2046. https://f1000research.com/articles/6-2046/v1.

Killick, Rebecca, and Idris A. Eckley. 2014. "changepoint: An R Package for Changepoint Analysis." *Journal of Statistical Software* 58 (3): 1–19. http://www.jstatsoft.org/v58/i03/.

Kitchin, Rob. 2014. *The data revolution : big data, open data, data infrastructures & their consequences.* Los Angeles, California London: SAGE Publications.

Kruppa, Jochen, Yufeng Liu, Gérard Biau, Michael Kohler, Inke R. König, James D. Malley, and Andreas Ziegler. 2014. "Probability Estimation with Machine Learning Methods for Dichotomous and Multicategory Outcome: Theory." *Biometrical Journal*, no. 4 (July): 534–63. https://doi.org/10.1002/bimj.201300068.

Kuhn, Max. 2008. "Building Predictive Models in R Using the caret Package." *Journal of Statistical Software* 28 (5). http://www.jstatsoft.org/v28/i05/.

Lee, Jae K. 2010. *Statistical Bioinformatics: For Biomedical and Life Science Researchers.* Wiley-Blackwell. https://www.wiley.com/en-sg/Statistical+Bioinformatics%3A+For+Biomedical+and+Life+Science+Researchers-p-9780471692720.

Mallona, Izaskun, Anna Díez-Villanueva, Berta Martín, and Miguel A. Peinado. 2017. "Chainy: an universal tool for standardized relative quantification in real-time PCR." *Bioinformatics.* https://doi.org/10.1093/bioinformatics/btw839.

Mallona, Izaskun, Julia Weiss, and Marcos Egea-Cortines. 2011. "pcrEfficiency: a Web tool for PCR amplification efficiency prediction." *BMC Bioinformatics* 12: 404. https://doi.org/10.1186/1471-2105-12-404.

McCall, Matthew N., Helene R. McMurray, Hartmut Land, and Anthony Almudevar. 2014. "On non-detects in qPCR data." *Bioinformatics* 30 (16): 2310–6. https://doi.org/10.1093/bioinformatics/btu239.

McFadden, Daniel L. 1974. "Conditional Logit Analysis of Qualitative Choice Behavior." In *Frontiers in Economics*, Frontiers in Economics:105–42. P. Zarembka (ed.). New York: Academic Press. https://eml.berkeley.edu/reprints/mcfadden/zarembka.pdf.

Nassirpour, Rounak, Sachin Mathur, Mark M. Gosink, Yizheng Li, Ahmed M. Shoieb, Joanna Wood, Shawn P. O'Neil, Bruce L. Homer, and Laurence O. Whiteley. 2014. "Identification of Tubular Injury microRNA Biomarkers in Urine: Comparison of Next-Generation Sequencing and qPCR-Based Profiling Platforms." *BMC Genomics* 15 (June): 485. https://doi.org/10.1186/1471-2164-15-485.

Neve, Jan De, Joris Meys, Jean-Pierre Ottoy, Lieven Clement, and Olivier Thas. 2014. "unifiedWMWqPCR: the unified Wilcoxon–Mann–Whitney test for analyzing RT-qPCR data in R." *Bioinformatics* 30 (17): 2494–5. https://doi.org/10.1093/bioinformatics/btu313.

Nolan, Tania, Rebecca E Hands, and Stephen A Bustin. 2006. "Quantification of mRNA using real-time RT-PCR." *Nature Protocols* 1 (November): 1559. https://doi.org/10.1038/nprot.2006.236.

Pabinger, Stephan, Stefan Rödiger, Albert Kriegner, Klemens Vierlinger, and Andreas Weinhäusel. 2014. "A survey of tools for the analysis of quantitative PCR (qPCR) data." *Biomolecular Detection and Quantification* 1 (1): 23–33. https://doi.org/10.1016/j.bdq.2014.08.002.

Pabinger, Stephan, Gerhard G. Thallinger, René Snajder, Heiko Eichhorn, Robert Rader, and Zlatko Trajanoski. 2009. "QPCR: Application for real-time PCR data management and analysis." *BMC Bioinformatics* 10 (1): 268. https://doi.org/10.1186/1471-2105-10-268.

Richards, F. J. 1959. "A Flexible Growth Function for Empirical Use." *Journal of Experimental Botany* 10 (2): 290–301. https://doi.org/10.1093/jxb/10.2.290.

Ritz, Christian, and Andrej-Nikolai Spiess. 2008. "qpcR: an R package for sigmoidal model selection in quantitative real-time polymerase chain reaction analysis." *Bioinformatics* 24 (13): 1549–51. https://doi.org/10.1093/bioinformatics/btn227.

Ronde, Maurice W. J. de, Jan M. Ruijter, David Lanfear, Antoni Bayes-Genis, Maayke G. M. Kok, Esther E. Creemers, Yigal M. Pinto, and Sara-Joan Pinto-Sietsma. 2017. "Practical data handling pipeline improves performance of qPCR-based circulating miRNA measurements." *RNA* 23 (5): 811–21. https://doi.org/10.1261/rna.059063.116.

Rote, Günter. 1991. "Computing the minimum Hausdorff distance between two point sets on a line under translation." *Information Processing Letters* 38 (3): 123–27. https://doi.org/10.1016/0020-0190(91)90233-8.

Rödiger, Stefan, Alexander Böhm, and Ingolf Schimke. 2013. "Surface Melting Curve Analysis with R." *The R Journal* 5 (2): 37–53. http://journal.r-project.org/archive/2013-2/roediger-bohm-schimke.pdf.

Rödiger, Stefan, Michał Burdukiewicz, Konstantin A. Blagodatskikh, and Peter Schierack. 2015. "R as an Environment for the Reproducible Analysis of DNA Amplification Experiments." *The R Journal* 7 (2): 127–50. http://journal.r-project.org/archive/2015-1/RJ-2015-1.pdf.

Rödiger, Stefan, Michał Burdukiewicz, and Peter Schierack. 2015. "chipPCR: an R package to pre-process raw data of amplification curves." *Bioinformatics* 31 (17): 2900–2902. https://doi.org/10.1093/bioinformatics/btv205.

Rödiger, Stefan, Michał Burdukiewicz, Andrej-Nikolai Spiess, and Konstantin Blagodatskikh. 2017. "Enabling reproducible real-time quantitative PCR research: the RDML package." *Bioinformatics*, August. https://doi.org/10.1093/bioinformatics/btx528.

Rödiger, Stefan, Thomas Friedrichsmeier, Prasenjit Kapat, and Meik Michalke. 2012. "RKWard: a comprehensive graphical user interface and integrated development environment for statistical analysis with R." *Journal of Statistical Software* 49 (9): 1–34. https://www.jstatsoft.org/article/view/v049i09/v49i09.pdf.

Rödiger, Stefan, Peter Schierack, Alexander Böhm, Jörg Nitschke, Ingo Berger, Ulrike Frömmel, Carsten Schmidt, et al. 2013. "A highly versatile microscope imaging technology platform for the multiplex real-time detection of biomolecules and autoimmune antibodies." *Advances in Biochemical Engineering/Biotechnology* 133: 35–74. https://doi.org/10.1007/10_2011_132.

Ruijter, Jan M., Peter Lorenz, Jari M. Tuomi, Michael Hecker, and Maurice J. B. van den Hoff. 2014. "Fluorescent-increase kinetics of different fluorescent reporters used for qPCR depend on monitoring chemistry, targeted sequence, type of DNA input and PCR efficiency." *Microchimica Acta*, 1–8. https://doi.org/10.1007/s00604-013-1155-8.

Ruijter, Jan M., Michael W. Pfaffl, Sheng Zhao, Andrej N. Spiess, Gregory Boggy, Jochen Blom, Robert G. Rutledge, et al. 2013. "Evaluation of qPCR curve analysis methods for reliable biomarker discovery: Bias, resolution, precision, and implications." *Methods* 59 (1): 32–46. https://doi.org/10.1016/j.ymeth.2012.08.011.

Ruijter, J M, C Ramakers, W M H Hoogaars, Y Karlen, O Bakker, M J B van den Hoff, and A F M Moorman. 2009. "Amplification efficiency: linking baseline and bias in the analysis of quantitative PCR data." *Nucleic Acids Research* 37 (6): e45. https://doi.org/10.1093/nar/gkp045.

Saeys, Y., I. Inza, and P. Larranaga. 2007. "A review of feature selection techniques in bioinformatics." *Bioinformatics* 23 (19): 2507–17. https://doi.org/10.1093/bioinformatics/btm344.

Scott, A. J., and M. Knott. 1974. "A Cluster Analysis Method for Grouping Means in the Analysis of Variance." *Biometrics* 30 (3): 507. https://doi.org/10.2307/2529204.

Seibelt, Pablo. 2017. *xray: X Ray Vision on your Datasets*. https://CRAN.R-project.org/package=xray.

Shin, Hoo-Chang, Holger R. Roth, Mingchen Gao, Le Lu, Ziyue Xu, Isabella Nogues, Jianhua Yao, Daniel Mollura, and Ronald M. Summers. 2016. "Deep Convolutional Neural Networks for Computer-Aided Detection: CNN Architectures, Dataset Characteristics and Transfer Learning," February. http://arxiv.org/abs/1602.03409.

Sing, Tobias, Oliver Sander, Niko Beerenwinkel, and Thomas Lengauer. 2005. "ROCR: visualizing classifier performance in R." *Bioinformatics* 21 (20): 3940–1. https://doi.org/10.1093/bioinformatics/bti623.

Spiess, Andrej-Nikolai, Claudia Deutschmann, Michał Burdukiewicz, Ralf Himmelreich, Katharina Klat, Peter Schierack, and Stefan Rödiger. 2015. "Impact of Smoothing on Parameter Estimation in Quantitative DNA Amplification Experiments." *Clinical Chemistry* 61 (2): 379–88. https://doi.org/10.1373/clinchem.2014.230656.

Spiess, Andrej-Nikolai, Caroline Feig, and Christian Ritz. 2008. "Highly accurate sigmoidal fitting of real-time PCR data by introducing a parameter for asymmetry." *BMC Bioinformatics* 9 (1): 221. https://doi.org/10.1186/1471-2105-9-221.

Spiess, Andrej-Nikolai, Stefan Rödiger, Michał Burdukiewicz, Thomas Volksdorf, and Joel Tellinghuisen. 2016. "System-specific periodicity in quantitative real-time polymerase chain reaction data questions threshold-based quantitation." *Scientific Reports* 6 (December): 38951. https://doi.org/10.1038/srep38951.

Tichopad, Ales, Michael Dilger, Gerhard Schwarz, and Michael W Pfaffl. 2003. "Standardized determination of real-time PCR efficiency from a single reaction set-up." *Nucleic Acids Research* 31 (20): e122.

Tierney, Nicholas. 2017. "Visdat: Visualising Whole Data Frames." *The Journal of Open Source Software* 2 (16).

Tolson, Edward. 2001. "Machine Learning in the area of image analysis and pattern recognition." *Advanced Undergraduate Project, Spring*. https://stuff.mit.edu/afs/athena/course/urop/profit/PDFS/EdwardTolson.pdf.

Walsh, Ian, Gianluca Pollastri, and Silvio C. E. Tosatto. 2015. "Correct machine learning on protein sequences: a peer-reviewing perspective." *Briefings in Bioinformatics*, September, bbv082. https://doi.org/10.1093/bib/bbv082.

Westermeier, Reiner. 2004. *Electrophoresis in Practice: A Guide to Methods and Applications of DNA and Protein Separations, Fourth Edition*. Wiley-Blackwell. https://doi.org/10.1002/3527603468.

Williams, Graham J. 2009. "Rattle: A Data Mining GUI for R." *The R Journal* 1 (2): 45–55. http://journal.r-project.org/archive/2009-2/RJournal_2009-2_Williams.pdf.

Zielesny, Achim. 2011. *From Curve Fitting to Machine Learning*. Edited by Janusz Kacprzyk and Lakhmi C. Jain. Vol. 18. Intelligent Systems Reference Library. Berlin, Heidelberg: Springer Berlin Heidelberg. https://doi.org/10.1007/978-3-642-21280-2.
